# Electrophysiological monitoring of plants: an exploratory study on drought stress

**DOI:** 10.64898/2025.12.03.692091

**Authors:** Raffaele Zappala, Lorenzo Gavazzeni, Matteo Gandolfi, Javier Babí Almenar, Maurizio Magarini, Andrzej Kurenda, Renato Casagrandi

## Abstract

Climate change is increasing environmental stress, particularly rising temperatures and water scarcity, in both natural and human-managed systems such as agroecosystems and urban environments. Traditional methods for monitoring plant health in human-managed systems remain limited, underscoring the need for novel approaches. This study explores the potential of plant electrophysiological signals (EPS) and derived statistical features for the early detection of drought stress. The two main objectives of this research are: i) to identify EPS features that are both ecologically relevant and statistically robust for detecting drought stress, and ii) to develop statistical models that integrate these features. EPS data was collected from two drought-stress experiments, one on tomato plants and one on apricot trees. Sixteen features from both time and frequency domains were selected and evaluated. Two models, a logistic and a machine learning classifier, were developed and compared using accuracy, precision, and recall metrics. In apricots, ten time-domain features (Frequency Center, Generalized Hurst Exponent, Hjorth Complexity, Hjorth Mobility, Kurtosis, Root Mean Squared Frequency, Root Variance Frequency, Shape Factor, Skewness and Standard Deviation) showed significant differences between stressed and control groups. In tomatoes, four frequency-domain features (Frequency Centre, Root Variance Frequency, Root Mean Squared Frequency, and Power Law Distribution Exponent) were significantly different. Model accuracy was approximately 50% for apricots and 66% for tomatoes, insufficient for practical deployment but indicative of potential. This study illustrates the potential value of plant EPS data, its derived statistical features, and models for developing early drought stress detection systems in both agricultural and urban plant management contexts.

## 1 Introduction

Climate change is significantly raising temperatures and exacerbating water scarcity, causing environmental stress in both natural and human-modified ecosystems, including agroecosystems and urban systems. Between 2000 to 2030, the extent of drought-affected regions is projected to nearly double (Güneralp et al., 2015). By 2050, southern and central Europe, in particular, are expected to experience unprecedented droughts under a 2°C global temperature rise (Guerreiro et al., 2018), with mid-latitude cities experiencing twice the levels of heat stress compared to their rural counterparts, regardless of climate change scenarios (Wouters et al., 2017). Business-as-usual planning and management actions exacerbate these challenges by altering biotic and abiotic factors, disrupting natural air and water flows, reducing heat dissipation, and increasing temperatures (Susca and Pomponi, 2020). Therefore, climate adaptation actions, combined with innovative monitoring techniques, becomes crucial to alleviate environmental stress and protect human health in urban and agricultural systems.

In urban systems, a recurrent climate adaptation and mitigation action focuses on the increase of urban greenery such as urban forests, green roofs, and constructed wetlands, which also help in restoring biodiversity (Seddon et al., 2020). Research, including the Intergovernmental Panel on Climate Change Assessment Report 6, shows that urban greenery can reduce heat through shading and evapotranspiration, lowering energy costs and reducing the demand for air conditioning by up to 30% (Ferreira and Duarte, 2019; Zardo et al., 2017). Prioritizing tree planting in areas with limited canopy coverage can maximize these benefits (Ziter et al., 2019). Instead, in agroecosystems, common climate adaptation measures include actions such as replacing water-intensive crops with drought-resistant species and adopting water-saving irrigation systems to maintain yields and revenue (Labeyrie et al., 2021). Despite the value of both sets of climate actions, rising temperatures and increasing water scarcity still can place substantial stress on the health of plant species (Filho et al., 2019).

A critical factor in maintaining vegetation health, especially during droughts and heatwaves, is ensuring adequate water supply. When management actions, such as irrigation, fail to meet plants’ water needs, drought stress occurs, often becoming evident only after visible symptoms like leaf wilting or discoloration appear (Osakabe et al., 2014). To prevent this, various methods have been developed to monitor plant physiological responses to water stress. For instance, a pressure chamber is commonly used to measure xylem water potential (Dale and Frank, 2022), but it is a destructive and time-consuming technique that requires strong technical expertise. Remote sensing is becoming widely adopted, particularly for monitoring large or inaccessible areas (Elsharkawy et al., 2022). It helps assess vegetation health using indices like the Normalized Difference Vegetation Index, which reflects the status of plant health and water content (Becker-Reshef et al., 2010). However, the fragmented and small-scale nature of urban green spaces, combined with their heterogenous land cover, requires remote sensing products with high temporal and spatial resolution (Ju et al., 2022; Neyns and Canters, 2022). This makes difficult to rely on remote sensing as a primary monitoring method in urban environments. As an emerging method, monitoring of electrophysiological signals (EPSs) is gaining attention for detecting plant stress, including drought, and has also been extensively studied in the animal kingdom and human medicine, where it is employed to detect brain, heart, stomach and muscle diseases (Rangayyan, 2015).

EPSs are also key components of plant communication systems, where they reflect responses to internal stimuli, such as water scarcity or nutrient deficiency, through changes in electrical potential (Najdenovska et al., 2021a). Although the underlying mechanisms are not fully understood, EPS are believed to result from changes in the electrical potential gradient across the plasma membrane of plant cells, often associated with hydraulic and chemical signals such as reactive oxygen species and calcium ion waves (Hilleary and Gilroy, 2018; Mudrilov et al., 2021; Sukhov et al., 2019). Furthermore, water deficiency alters the osmotic relations between different plant organs and cellular compartments, and affects the electrical properties of the tissues (Serrano-Finetti et al., 2023). Therefore, it is expected to have direct reflections in the electrophysiology recordings. Among the various types of plant EPSs, action potential (AP), variation potential (VP), and system potential (SP) are the most extensively studied (Vodeneev et al., 2016; Wairimu et al., 2021; Zimmermann et al., 2009). APs are typically associated with short-term stress signals and triggered by non-damaging stimuli, whereas VPs and SPs are more indicative of long-term stress responses usually triggered by damaging stimuli (Wairimu et al., 2021). Studies on drought stress in plants have already explored changes in AP, VP and SP propagation under different levels of water deficit (Yuan et al., 2023; Zhou et al., 2022a). Precedent studies showcase that EPS may adequately reflect physiological processes in plants, including respiration and photosynthesis, which has sparked increasing interest in their use for monitoring plant responses to environmental stress.

To understand plant stimuli or stress through EPS observation, researchers often analyse their statistical features in both time and frequency domains (García-Servín et al., 2021). This kind of analysis is frequently complemented by techniques such as machine learning (ML) algorithms (Damm et al., 2018; Rouse et al., 1973). EPS statistical features, referred to as “features” from here on, often serve as inputs for ML models that classify plants as healthy or stressed with high accuracy (González I Juclà et al., 2023; Najdenovska et al., 2021c; Tran et al., 2019a; Tran and Camps, 2021; Wen et al., 2022). ML models integrating EPS features have been applied to a wide range of abiotic stressors, like lack of light, cold stress, osmotic stress (Pereira et al., 2018a), pollution (e.g., sodium chloride, sulfuric acid, ozone) (Damm et al., 2018; Rouse et al., 1973), drought (Tran et al., 2019a), and salt tolerance (Qin et al., 2020). They have been also applied to detect biotic stressors like fungal infections (Simmi et al., 2020a), spider mite infestations (Najdenovska et al., 2021a), and herbivore-induced wounds (Choi et al., 2016; Farmer et al., 2020; Zimmermann et al., 2009). Common ML models used in EPS research include Linear Regression, Deep Learning, Decision Trees, Random Forests, Gradient Boost, and Extreme Gradient Boost (Najdenovska et al., 2021c; Tran et al., 2019a).

A close examination of the literature highlights the proven utility of specific features in both the time and frequency domains for evaluating plant stress and stimuli. In the time domain, minimum and maximum signal values, combined with variance, have been used to detect long-term signal changes, such as decreases in EPS over time (Tran et al., 2019a) and the attenuation of a burning-induced VP (Yudina et al., 2022), both under drought regime. The interquartile range has shown strong discriminative power in identifying biotic stressors, like spider mite infestations in tomato plants (Najdenovska et al., 2021a), and has been identified as a dominant feature for distinguishing environmental stimuli (Chatterjee et al., 2015a). Skewness has proven effective in classifying ozone and osmotic stress (Chatterjee et al., 2015a), while General Hurst Exponent has been used to assess osmotic stress and various biotic and abiotic stressors (Chatterjee et al., 2015a; Tran et al., 2019a). Hjorth Complexity showed promising results for classifying osmotic stress (Chatterjee et al., 2015a), spider mites’ infestation (Najdenovska et al., 2021a) and to analyse electrical responses to critical levels of ozone pollution (Dugelay and Slock, 2015). Hjorth Mobility has been applied to study biotic and abiotic stressors in tomato plants (Chatterjee et al., 2015a; Tran et al., 2019a). In the frequency domain, Fourier Transform has been used to study responses under osmotic stress in maize (Zhang et al., 2012), while power frequency analysis has shown promise in studying electrical extracellular recordings (Zhang et al., 2012). Power Spectrum Analysis has been used to simulate electrical models in Araceae plants (Cabral et al., 2011). Wavelet analysis has proven useful for investigating non-stationary EPSs (Li et al., 2015), with wavelet decomposition, employed for signal denoising (Jinli et al., n.d.; Tian et al., 2021). Specific features, such as gravity frequency and Power Spectrum Entropy, along with peak-to-peak values, in the time-domain, have shown strong potential for capturing the non-linearities in plant electrical signals (Yuan et al., 2023).

Despite advancements in plant EPS research, several challenges remain. *One major gap* is the *limited biophysical interpretation of EPS features in terms of their ecological meaning*, as using these features to indicate plant health is still a relatively new area of study. For instance, in the context of drought stress, only a small number of studies have investigated plant responses (Bekkari et al., 2024; Cattani et al., 2024a; Dolfi et al., 2021; Tran et al., 2019a; Wen et al., 2022), and even fewer have attempted to *provide a biophysical or ecological explanation for the extracted feature values* (Chatterjee et al., 2015a; Yuan et al., 2023). *As a second gap*, *feature selection in plant EPS research has not always been guided by plant-specific ecophysiological considerations*. In some studies, features are chosen based on evidence from medical studies, due to the shared biological origins, similarities in EPS generation processes, and the diagnostic power of certain features in medical applications (Alotaiby et al., 2019). As a result, *the features used may not be the most suitable to capture plant stress responses*. *A third gap* concerns the *widespread use of ML models without a clear understanding of the causal relationships between EPS features and plant stress responses*. While some EPS studies have demonstrated the discriminative power of EPS features within ML models, achieving promising results in binary classification tasks (Chatterjee et al., 2015a; Yuan et al., 2023), these models often function as ‘*black boxes*,’. This further limits our understanding of the *causal or ecological significance of input features*, and hampers efforts to link EPS features with underlying stress mechanisms.

In light of these research gaps, this study investigates drought stress responses in plants:

I. to identify EPS features that are both ecologically meaningful and statistically robust, and
II. to develop statistical models that can inform early-warning tools for plant stress monitoring.

From a practical standpoint, building on emerging work on plant-related EPS, this study further explores the potential of EPS analysis as a monitoring technique for the early detection of plant stress, with an initial focus on drought. The practical aim is to contribute to the growing body of knowledge on how EPS analysis might complement existing plant-monitoring approaches in human-managed ecosystems.

## 2 Methods and Materials

### 2.1 Description of the two datasets and EPS monitoring device

The datasets corresponded to plant drought stress experiments on tomato (*Solanum lycopersicum*) and apricot (*Prunus armeniaca*) plants. Both datasets were provided by Vivent S.A. and were collected using a PhytlSigns® device, a tool capable of recording high-quality signals in both controlled and outdoor environments. The signal was sampled at 256 Hz, a frequency that was considered too fine for the problems at hands, thus the signals were then down sampled at 1 Hz. This frequency is indeed considered high enough to detect the relevant patterns in the plant’s EPS, because APs, VPs and SPs usually last a few minutes (Li et al., 2021).

The tomato experiment dataset included the electrical potential recordings (in mV) of 16 tomato plants: 8 control plants and 8 drought-stressed plants. The experiment was conducted indoors, in a commercial greenhouse condition, under controlled light and temperature. Each recording channel recorded electrical potential from one plant with use of Ag/AgCl electrodes. For each channel reference electrodes were inserted into lower part of the stem about 10 cm above substrate level. Active electrodes were inserted about 1 m above reference electrodes. Data were recorded for nine days (between 29/10/2020 and 07/11/2020), and the analysis focussed in particular on the days from 02/11/2020 to 07/11/2020, as only then drought symptoms were manifested in the stressed plants. One out of these of stressed plants was excluded from the analysis, as its signal was clearly affected by a disturbance. There were some gaps in the data during the days of 02/11, 04/11 and 06/11, possibly due to a malfunction of the recording device. For the control plants, the gaps lasted approximately one minute, while for the stressed plants these gaps lasted approximately one hour.

The apricot experiment dataset consists of electrical potential recordings from 16 young apricot trees (4 years old), with 8 control plants and 8 drought-stressed plants. Electrodes were connected similarly as in the case of tomato plants: reference electrodes were connected in the shaded side of the tree trunk about 10 cm above soil. Active electrodes were connected in an illuminated side of trunk about 1m above reference electrode, usually the region of first branches. Experiment was conducted outdoors in a residential backyard in Renens, Switzerland, from 01/05/2022 to 15/07/2022. The plants were kept in 10-liter pots, with soil covered to reduce water evaporation. The control group was watered every 3-5 days, depending on weather conditions, while the stress group was not watered after 15/05/2022. In addition, differently from the tomatoes of the previous experiment, every day the plants were visually inspected and classified according to their appearance in “normal development”, “wilting” and “strong wilting”. Even in this experiment two plants, one from the control group and the other from the stressed group, were also excluded due to a disease. Another plant from the control group was excluded due to several large gaps in the recording data. The first day included in the analysis was 02/06/2022, as early visual signals of drought stress were already confirmed. The signals of all these plants were studied until 17/06/2022.

### 2.2 Assessment of the ecological relevance and statistical significance of selected features in time-based and frequency-based domains

We selected 16 features from the time and frequency domain that could be of potential value for detecting plant drought stress (Table 1). The selection of features was informed by a scoping literature review of precedent plant EPS studies. We selected features that appeared to be statistically relevant to identify different types of stressors in other plant studies, even if those studies were not related to drought stress.

**Table 1.** The features we selected to analyse the signal, listed together with a brief description, their formula, and the reference to previous uses. The Generalized Hurst Exponent is computed through several steps, which, due to space limitations, are reported in Supplementary Material 2.

| Feature | Domain | Description | Formula | Refs |
| --- | --- | --- | --- | --- |
| Minimum (*Min*) | Time | The lowest value of the signal in the window. In the formula, $y(t)$ indicates the signal. | $Min = \min(y(t))$ | (Tran et al., 2019b; Yudina et al., 2022) |
| Maximum (*Max*) | Time | The highest value of the signal in the window. | $Max = \max(y(t))$ | (Tran et al., 2019b; Yudina et al., 2022) |
| Standard deviation (*Sd*) | Time | The amount of variation or dispersion of the signal in the window. $E[\cdot]$ indicates the average function, while $\mu$ indicates the average of the signal. | $sd = \sqrt{E[(y(t) - \mu)^2]}$ | (Tian et al., 2015; Zhou et al., 2022b) |
| Skewness (*Skw*) | Time | A measure of the asymmetry about its mean of the probability distribution of a windowed signal. | $Skw = E\left[\left(\frac{y(t) - \mu}{sd}\right)^3\right]$ | (Chatterjee et al., 2017, 2015b) |
| Kurtosis (*Krt*) | Time | A measure of the dispersion, or "tailedness", of the probability distribution of the signal. | $Krt = E\left[\left(\frac{y(t) - \mu}{sd}\right)^4\right]$ | (Chatterjee et al., 2017, 2015b) |
| Interquartile range (*IQR*) | Time | A measure of dispersion, evaluated as the difference between third ($Q_3$) and first ($Q_1$) quartile. | $IQR = Q_3 - Q_1$ | (Chatterjee et al., 2017, 2015b) |
| Hjorth mobility (*Hm*) | Time | A measure of the rapidity of changes of a signal. $y'(t)$ is the first derivative of the signal, while $\text{var}(\cdot)$ is the variance. | $HM = \sqrt{\frac{\text{var}(y'(t))}{\text{var}(y(t))}}$ | (Bhadra et al., 2023; Najdenovska et al., 2021b) |
| Hjorth complexity (*HC*) | Time | A measure of signal complexity, reflecting the irregularity or intricacy of a signal. $y''(t)$ is the second derivative of the signal. | $HC = \frac{\sqrt{y''(t)/y'(t)}}{\sqrt{y'(t)/y(t)}}$ | (Bhadra et al., 2023; Najdenovska et al., 2021b) |
| Impulse factor (*IF*) | Time | The ratio of the peak value to the mean value. It is a measurement that provides information on the extremes of the signal. | $IF = \frac{\max(\vert y(t)\vert )}{E[\vert y(t)\vert ]}$ | (Tran et al., 2024, 2019b) |
| Crest factor (*CF*) | Time | Similarly to the *IF*, it measures the presence of peaks. While the *IF* confronts the peaks with the average, the *CF* confronts it with the standard deviation. | $CF = \frac{\max(\vert y(t)\vert )}{\sqrt{E[y(t)^2]}}$ | (Tran et al., 2024, 2019b) |
| Margine factor (*MF*) | Time | Another metric used to compare peak values with the remaining part of the signal. | $MF = \frac{\max(\vert y(t)\vert )}{(E[\sqrt{\vert y(t)\vert }])^2}$ | (Tran et al., 2024, 2019b) |
| Generalised Hurst Exponent (*GHE*) | Time | It computes the long-range correlation in the time series, that is its long-term memory. | Formulation in the Supplementary Material 2 | (Najdenovska et al., 2021b; Tran et al., 2019b) |
| Frequency Centre (*FC*) | Time / Frequency | The average frequency value over the spectrum, weighted for the respective amplitude. In the formula, $w_k$ represents the frequency (Hz) and $A_k$ the relative amplitude (dB) | $FC = \frac{\sum_{k=0}^{N-1} \omega_k A_k}{\sum_{k=0}^{N-1} A_k}$ | (Najdenovska et al., 2021b, 2021d; Tran et al., 2024) |
| Root Variance Frequency (*RVF*) | Time / Frequency | It is the variance of the signal's frequencies weighted for their amplitude values. It measures the dispersion of the signal in the frequency domain. In the formula, $\bar{\omega}$ is the average frequency. | $RVF = \frac{\sum_{k=0}^{N-1} (\omega_k - \bar{\omega})^2 A_k}{\sum_{k=0}^{N-1} A_k}$ | (Najdenovska et al., 2021b, 2021d; Tran et al., 2024) |
| Root Mean Square Frequency (*RMSF*) | Time / Frequency | The RMSF is the amplitude of a direct current that carries the same energy as the signal, so it is a measure of central tendency of the signal. | $RMSF = \sqrt{\frac{\sum_{k=0}^{N-1} \omega_k^2 A_k}{\sum_{k=0}^{N-1} A_k}}$ | (Najdenovska et al., 2021b, 2021d; Tran et al., 2024) |
| Power Spectral Density ($\beta$) | Frequency | The Power Spectral Density (PSD) is a decreasing function of the frequency. When it follows a power law, the exponent $\beta$ of the power law describes how fast the PSD decreases. When $\beta$ is lower, the signal carries more energy in the lower frequencies, and vice versa. | $A_k \sim \omega_k^\beta$ | (Simmi et al., 2020b; Souza et al., 2017) |

Before calculating the features, the EPS data was pre-processed according to well established procedures (Damm et al., 2018; Najdenovska et al., 2021c), which included: i) downsampling; ii) noise filtering; iii) outliers detection and their removal; and iv) missing data imputation. The pre-processing steps are detailed in Supplementary Material (SM) 1.

#### 2.2.1 Comparisons across time

The features in the time domain cannot be computed on each point. As an example, the standard deviation can be computed only on a set of points. For this reason, it was necessary to divide the signal into equal-length windows (*l*), whose size had to be determined, and to compute the features on these windows. Small windows did not adequately capture the signal’s behaviour. Conversely, large windows oversmoothed the signal. After iterative tests, a window length of 400 seconds was considered the most suitable for statistical analysis, providing a clear visual output. Further window lengths were instead used for the machine learning models (as described in Section 2.3). For the statistical analysis, described at the end of this section, the feature values were split into night (night: 9:00 pm to 6:00 am) and day segments. The latter helped to analyse more clearly whether circadian rhythms influenced feature values.

All the features in the frequency domain were computed splitting the data into day and night segments, without the use of windows of 400 seconds, except for FC, RVF and RMSF, for which it was possible to apply also the window length of 400 seconds used for time-domain features. Building on past EPS research (Chen et al., 2016), analysis in the frequency domain was focused on the amplitude spectra (*A*_*k*_), namely on the values of the magnitude (dB) associated to each frequency component. First, we computed the Discrete Fourier Transform (DFT) to transform the original signal values into a series of coefficients of a linear combination of complex sinusoids sorted by frequency.

For the Power Spectral Density (PSD), according to the procedure used in (Simmi et al., 2020c), Welch’s algorithm (Averaged Periodograms) was used instead of FFT, because that algorithm works well with evenly spaced samples in time, as our recordings are. The resulting PSD data was log-transformed so as to inspect whether linear regression (Ordinary Least Squares) could permit to estimate a significative slope with respect to the logarithm of the frequency and, possibly, to see whether such a slope could help discriminate between stressed and control plants. This approach was compared to a non-linear regression on the original variables, according to the Akaike Information Criterion (AIC) (Xiao et al., 2011). The method favoured by the AIC was employed. After estimating α and β (see Eq. 1), it was possible to represent the interpolating curve in the original plane with the inverse logarithmic transform:

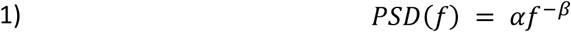

Where:

- PSD is the Power Spectral Density
- *f* is the frequency
- α and β are the parameters to be estimated.

In order to obtain meaningful values for the features, we normalised the data. This was necessary because initial calculations revealed large value ranges for most features, which prevented meaningful comparisons. Three normalization approaches were tested: i) normalization of feature values over the entire sample, ii) normalization of feature values per individual plant, and iii) normalization of the signal over the entire sample before calculation of the features. We chose the third approach, which provided more comparable data.

#### 2.2.2 Comparisons based on the plants’ health status

The EPS data from both tomato and apricot tree experiments were analysed by plant. However, the duration of the apricot tree experiment, combined with the visual categorization of plants into three groups (normal development, wilting, and strong wilting), enabled investigation of: i) whether certain feature values change gradually or abruptly as plants undergo increasing drought stress; and ii) whether statistical analysis of these features can reliably distinguish between the three groups. For conducting these additional analyses it was necessary to reorder the timeline and the aggregation of individual plant data.

In particular, the first analysis involved adjusting the data for stressed plants based on the first signs of wilting, rather than using calendar days. This means that, among stressed plants, the features’ values corresponding to the first day with visible signs of wilting were aggregated, although they corresponded to different calendar days, before comparison to the control plants. The same happened to the second day after the onset of wilting, and so on. This allowed for a consistent comparison between stressed and control plants by aligning the data within a time window of five days before and five days after the onset of wilting. For the second analysis, feature values were aggregated according to the visual classification groups, allowing for a comparison based on group-level averages rather than individual plants. During these two analyses, a visual inspection suggested that plants under drought stress might exhibit changes in the variability of some feature values. To confirm this in our result, we ran a third type of analysis, in which the interquartile range of each feature during the wilting period was compared to the period before visible wilting occurred to quantitatively assess whether there was an increase in variability. More specifically, for each feature and for each day, the interquartile range of the seven values (one for each stressed plant) was computed, and the interquartile ranges of the two sets of days – those before and those after the start of wilting – were compared through a Mann-Whitney test.

#### 2.2.3 Statistical methods

The Fisher’s Discriminant Ratio (FDR) and the Mann-Whitney tests were used to evaluate if each feature had the power of significantly discerning between control and stressed plants. The FDR measures how well a feature separates two or more groups, in this case control and stressed plants. It has previously been applied to classify plant health status (Chatterjee et al., 2015b). An FDR threshold of 1 was used to identify features with good potential for group differentiation. The exceedance of this threshold means that the average difference between sample features from two different classes is greater than the within-class standard deviation (Xu and Lu, 2006). This means that the classes’ centres are further apart than the classes are wide. The Mann-Whitney test was used to determine whether there were statistically significant differences in the distributions of feature values between control and stressed groups. This non-parametric test was preferred over the t-test because we were not a priori guaranteed that the data could consistently follow a normal distribution. Features with a p-value of 0.05 or below were considered statistically significant and potentially useful for distinguishing between the two groups. For the analyses of data aggregated into groups of different wilting levels only the Mann-Whitney test was applied. The outcomes of the statistical analyses provided a basis for discussing the biophysical interpretation of EPS features, and their potential ecological relevance for detecting drought stress.

### 2.3 Development of logistic and machine Learning models and comparison of their performance

A logistic classifier was developed using the features presented in the previous section. They were used as input for the logistic model, maintaining the window length of 400 seconds. The logistic non-linear regression algorithm (Equation 2) was employed, where *Y* represents the class type (control or stressed) and equals one if the window of the signal belongs to a stressed plant and zero otherwise, while *X_i_* ∈ *n* refers to the input data of each selected feature, computed on a window of the signal. Pr[*Y*=1] is the estimated probability that the window of the signal belongs to a stressed plant. γ_0_ and γ_i_ are the model parameters, and they were estimated using Maximum Likelihood Estimation.

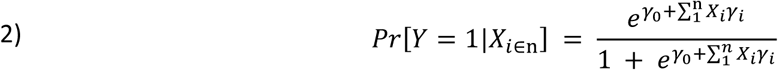

For the machine learning approach, we selected the Extreme Gradient Boosting (XGB) algorithm, as it has shown strong performance in previous EPS studies (Najdenovska et al., 2021c; Tran and Camps, 2021). XGB builds an ensemble of decision trees, where each successive tree attempts to reduce the errors made by the ensemble of the previously selected trees. During the training phase, the algorithm adds new trees to the model to minimize the loss function, which is a representation of the model’s error. Each new tree is built to reduce the residual loss of the previous trees. For further details on XGB algorithm can be found in (Chen and Guestrin, 2016).

In the XGB model, different temporal window lengths were tested, in addition to the default window of 400 seconds. Besides the use of windows, for this kind of model the signal needs to be segmented into intervals, each containing a set number of windows. To do this, the signal was firstly divided into windows *w_i_* of length *l*, that is window *w_i_* included all time steps between *l*·*(i-1)* and *l*·*i*. Secondly, the intervals were defined, where the first interval included all windows from *w_1_* to *w_N_*, the second interval were shifted by one window, including all windows from *w_2_* to *w_N+1_*, the third interval included all windows from *w_3_* to *w_N+2_*, and so on. Note that neighbouring intervals partly overlap, while neighbouring windows were always separated.

Once features were computed, their mean value per interval was estimated as a statistical aggregator, which served as the input for the XGB model. This feature value aggregation helps to reduce storage requirements and mitigates overfitting risks.

The outputs of both models (the logistic regression and the XGB) were evaluated using three metrics: accuracy, precision and recall (Eq. 3). The first measures the ratio of correctly predicted samples to the total number of samples. The second is the percentage of positive samples, i.e. samples from drought-stressed plants, among the one that were considered drought-stressed by the model. The third represents the fraction of correctly predicted samples of a chosen class among all the samples from drought-stressed plants.

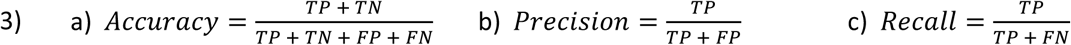

In Eq. 3, TP (true positive) refers to plants that have been correctly classified as drought stressed. TN (true negative) refers to plants that have been correctly classified as control cases. FP (false positive) refers to plants that are healthy but have been classified as stressed. FN (false negative) refer to plants that suffer drought stress but are classified as healthy.

To validate both models, leave-one-out cross-validation was employed. In each trial, one channel was used for testing while the remaining channels formed the training set, resulting in seven different runs. Accuracy, precision and recall metrics were averaged from all the trials through the median and the interquartile range.

Two hyperparameters are important for the XGB: the number of windows in each interval (*N*) and the window length (*l*). They were manually varied: *N* assumed values among (5, 10, 15, 20, 30, 35) and *l* assumed values among (120, 180, 240, 300, 660, 780, 900). The other hyperparameters (learning rate, number of decision trees, maximum tree depth and the number of features considered by each tree) were calibrated through Bayesian optimization. The performance of each (*N,l)* value was assessed according to the average accuracy and to the standard deviation of the accuracy across the leave-one-out validation. Since the average and the standard deviation of the accuracy are two different metrics, it is not possible to rank (*N,l)* values according to their performance. For example, one (*N,l)* value might outperform another one according to one metric, e.g., the average accuracy, and underperform it according to the other metric, e.g., the standard deviation of the accuracy. However, there can be (*N,l)* values that are outperformed by another (*N,l)* value according to both metrics. These are called “dominated solutions”. When these solutions are discarded, the remaining ones are called the “Pareto solutions”. When two Pareto solutions are compared, neither is strictly better than the other: each performs better on one metric and worse on the other. For this reason, the Pareto solutions express a trade-off between the average of the accuracy and its standard deviation.

In addition to predicting the plant’s status (control or stressed), the features of the models were also ranked according to their relevance. The relevance was calculated according to the Shapley values (Shapley, 1951), which are based on several runs of the models, each with a different subset of the features. For each run, the features were deleted one by one, to assess the decrease of the accuracy (as indicated in Eq. 3a) that followed the deletion of that feature. This decrease is considered a proxy to the contribution from that feature. The Shapley value for a feature is obtained by averaging, across the runs, the value of this proxy.

## 3 Results

### 3.1 Statistical significance of selected features

#### 3.1.1 Time domain analysis

In the comparison across time for the tomato plants the FDR was above one only in few cases, without an apparent pattern. This is illustrated in Figure 1, which shows the results for the features that had an FDR above one for any daylight or nighttime period. Only three features (KRT, RVF and GHE) out of sixteen exceeded the threshold for the FDR at least once. Furthermore, each of these features show either one or two instances of the exceedance of the threshold, out of eleven days or nights. Lastly, these features do not show any relevant pattern, such as an increasing or decreasing FDR across time. Also for the apricot, only in few cases the FDR was above one: nine features (FC, GHE, HM, RMSF, MAX, MIN, RVF, SF and SKW) exceeded the threshold at least once, and the most distinct results are illustrated in Figure 2. Notably, some features showed more than one exceedance of the threshold, often in consecutive periods. The Generalized Hurst Exponent, in particular, showed four exceedances within four days (Figure 2C). Similar results are obtained through the Mann-Whitney test, shown in Figure 2, on the right. For the apricot experiment, many features also showed an increasing separation between the control and the stressed plants over time, as visualised in Figure 2 for FC, GHE, HM and RMSF. This suggests that the values of the features tend to depart from their baseline values when the plant starts suffering drought stress.

**Figure 1:**
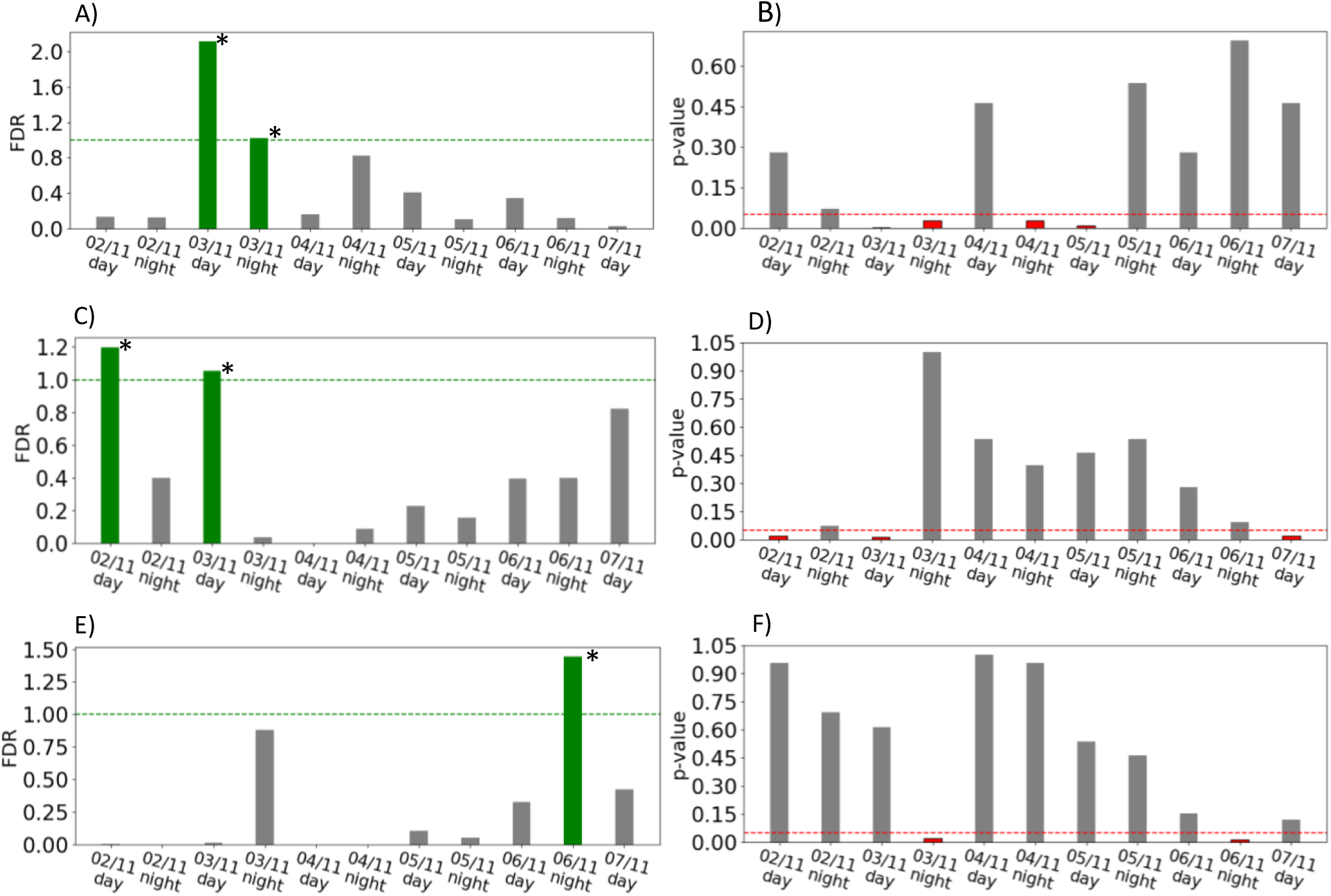
FDR (on the left column) and p-value of the Mann-Whitney test (on the right column), showing the separation between control and stressed tomatoes, for the features which show the greatest separation, in the form of an FDR above one. The features are the Kurtosis (A and B), the Root Variance Frequency (C and D) and the Generalized Hurst Exponent (E and F). An asterisk marks the periods when both the FDR exceeds one and the Mann-Whitney test shows a statistically significant difference. See SM 3 for the results for all features.

**Figure 2:**
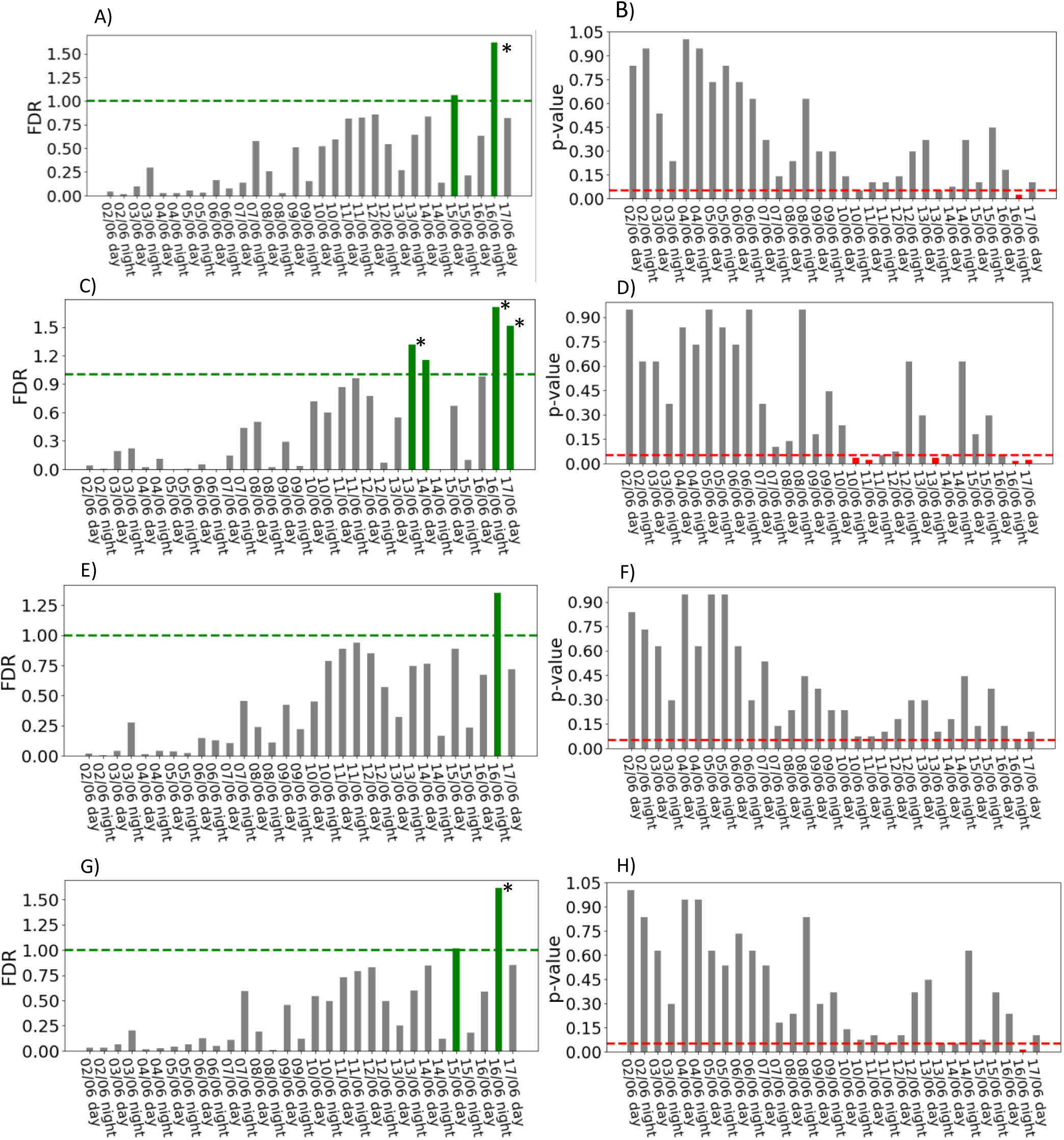
FDR (on the left column) and p-value of the Mann-Whitney test (on the right column), showing the separation between control and stressed apricots, for the most relevant features. The features are the Frequency Center (A and B), the Generalized Hurst Exponent (C and D), the Hjorth Mobility (E and F) and the Root Mean Squared Frequency (G and H). An asterisk marks the periods when both the FDR exceeds one and the Mann-Whitney test shows a statistically significant difference. See SM 3 for the results for further features.

Further understanding of changes in feature values, as apricot plants undergo increasing drought stress, is gathered by conducting analyses according to the plants’ health status, i.e. aggregating them in stressed and control samples. As explained in Section 2.2, the first of these analyses required shifting the data of the stressed plants, in order to match the start of the wilting periods of all of them. The results are shown in Figure 3, which shows that some features, in particular FC and RVF, tended to show a gradual increase in their values as plants underwent drought stress (Figure 3 A-D). Instead, HC showed an abrupt change in values when plants started to undergo drought stress, this was even more clearly visible through the Mann-Whitney test (Figure 3F).

**Figure 3:**
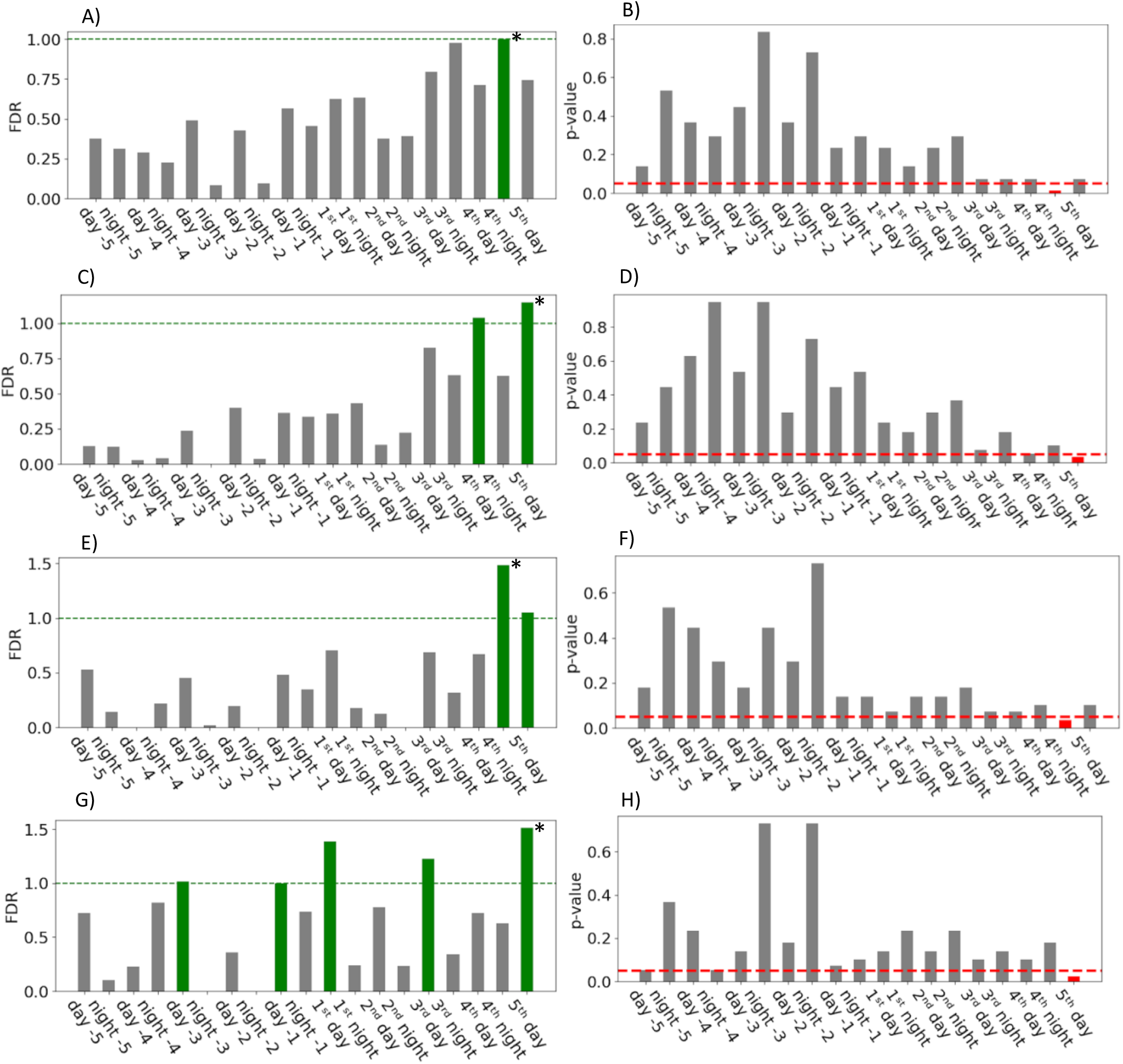
FDR (left column) and Mann-Whitney tests (right column) showing the separation between control and stressed apricots, for the most relevant features. Stressed plants are evaluated in the days around the first visible sign of wilting. In the figure, 1^st^ day refers to the first day of the first signs. The features are the Frequency Center (A and B), the Root Variance Frequency (C and D), the Hjorth Complexity (E and F) and the Generalized Hurst Exponent (G and H). See SM 3 for the results for further features.

The second of the aggregated analyses for apricots compared the categorical groups (control, wilting and strongly wilting) and the results are shown in Figure 4. It shows that several features were significantly different between the control and the stressed (i.e. wilting and strongly wilting) plants. However, feature values of the stressed plants were similar regardless of the intensity of the stress, namely wilting and strongly wilting plants had similar feature values (Figure 4). The values of most of the features (FC, GHE, HM, RMSF, RVF) increase when the plant is wilting, while the values of the HC decreases. In control plants, the GHE assumes values around 0.5 (Figure 4), while in stressed plants it assumes higher values. The boxplots also show that the dispersion around the median value is often larger for the stressed plants.

**Figure 4:**
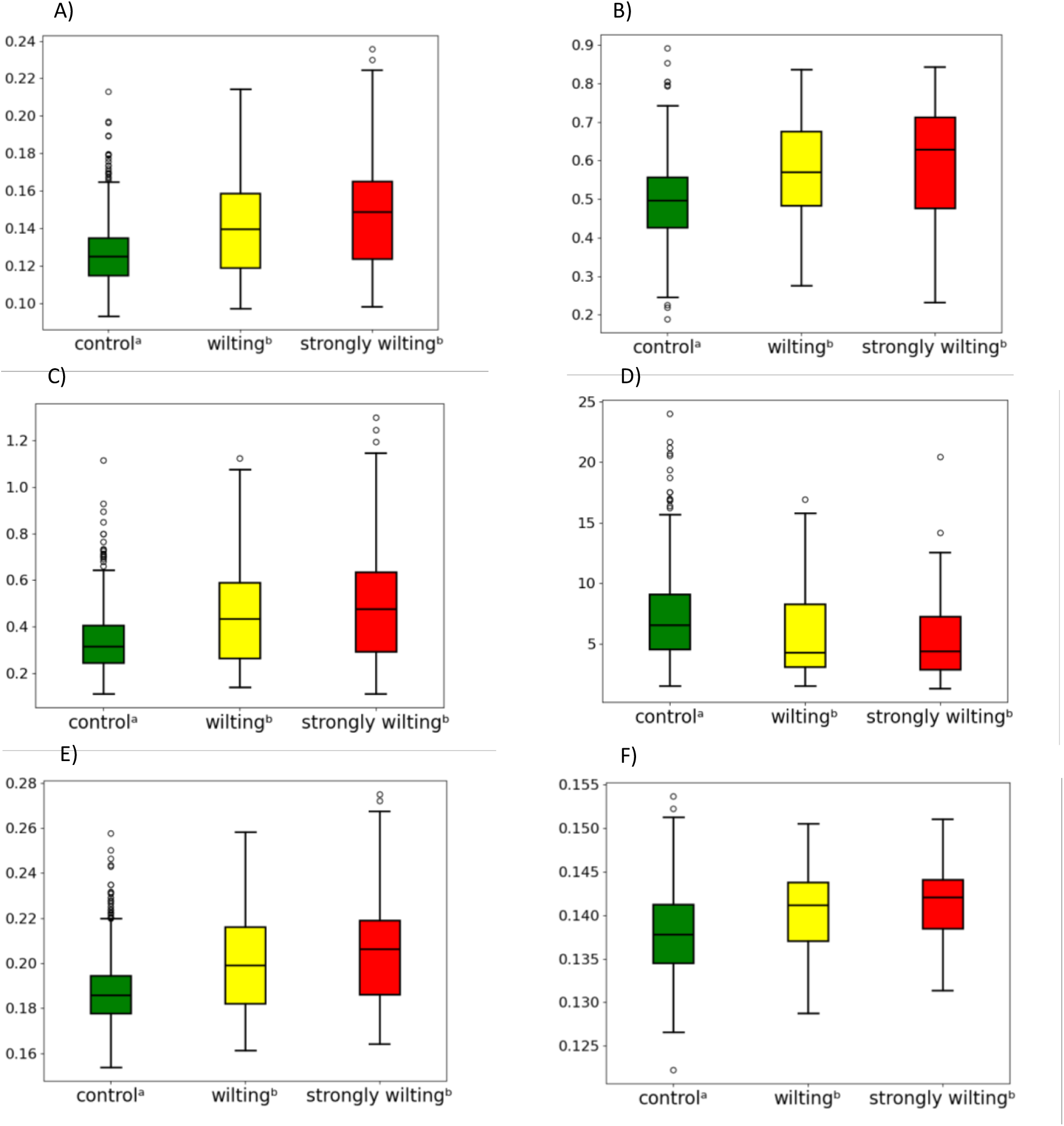
Comparison of the value of the features computed for the control apricots and for the stressed apricots in two different conditions: wilting and strongly wilting. The superscript letters indicate statistically significant differences according to the Mann-Whitney test. The features are the frequency Center (A), the Generalized Hurst Exponent (B), the Hjorth Mobility (C), the Hjorth Complexity (D), the Root Mean Squared Frequency (E) and the Root Variance Frequency (F). See SM 3 for the results for further features.

The third additional analysis on the apricot plants compared the interquartile range of the features’ value before and after the start of wilting, as explained in Section 2.2. Three features showed a significantly higher interquartile range after the plants started wilting: Frequency Centre, Root Variance Frequency and Root Mean Squared Frequency (Figure 5). The other features did not show a significant difference in the interquartile range (boxplots shown in SM3).

**Figure 5:**
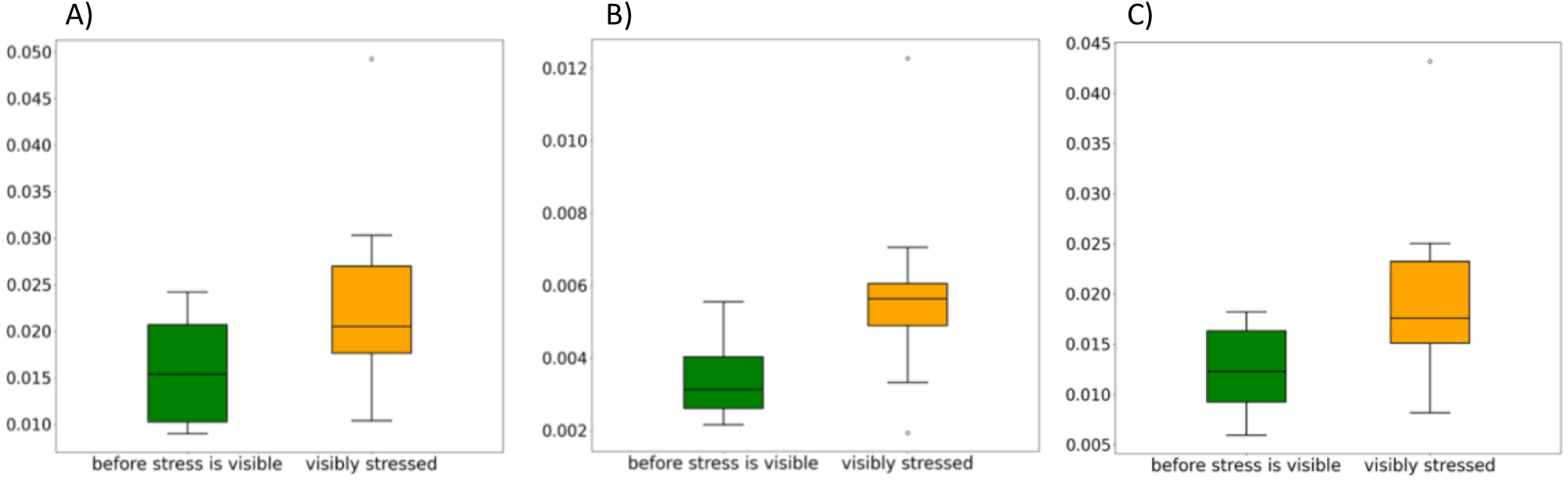
Interquartile range of three features in the stressed apricots, in the six days before the first visible signs of wilting and on the following six days. The features are the Frequency Center (A), the Root Variance Frequency (B) and the Root Mean Squared Frequency (C). See SM 3 for the results for the remaining features.

In the analyses on the apricot plants, FC and RMSF remain consistently among the features that indicate drought stress. Other features were relevant in more than one analysis: RVF, GHE, HM and HC. Three of these features, namely FC, RMSF and RVF, are computed on the frequency spectrum of the signal. This suggests that the impact of drought stress on the signal might be more easily detected in the frequency domain.

Further results from time-domain features, all FDR and Mann–Whitney analyses, by plant and aggregated, are included in SM3 for both datasets.

#### 3.1.2 Frequency domain analysis

For tomato plants, frequency features have FDR values greater than one for all features and across all days during the daytime periods, with the exception of 05/11, when only the RVF shows an FDR greater than one. For the nighttime periods, only the Power Spectral Density exhibits an FDR value greater than one, and only for the nights of 04/11 and 05/11. For apricot plants, the FDR value is greater than one only for Power Spectral Density on the night of 07/06 and the day of 15/06, and for the Root Variance Frequency on the night of 16/06. Further results on the FDR and Mann–Whitney analyses of both datasets for frequency features are included in SM3.

For both control and drought-stressed tomatoes, FC values are higher at night, which means that lower frequencies are stronger during day (Figure 6A1). This change in FC values is larger in control plants, while stressed plants’ response remains more constant over a certain frequency range throughout the 24 hours. The same pattern is observed for the RVF and the RMSF (Figure 6A2-6A3): the features’ values decrease during the daylight for the control tomatoes, while they are constant throughout day and night for the stressed plants.

**Figure 6:**
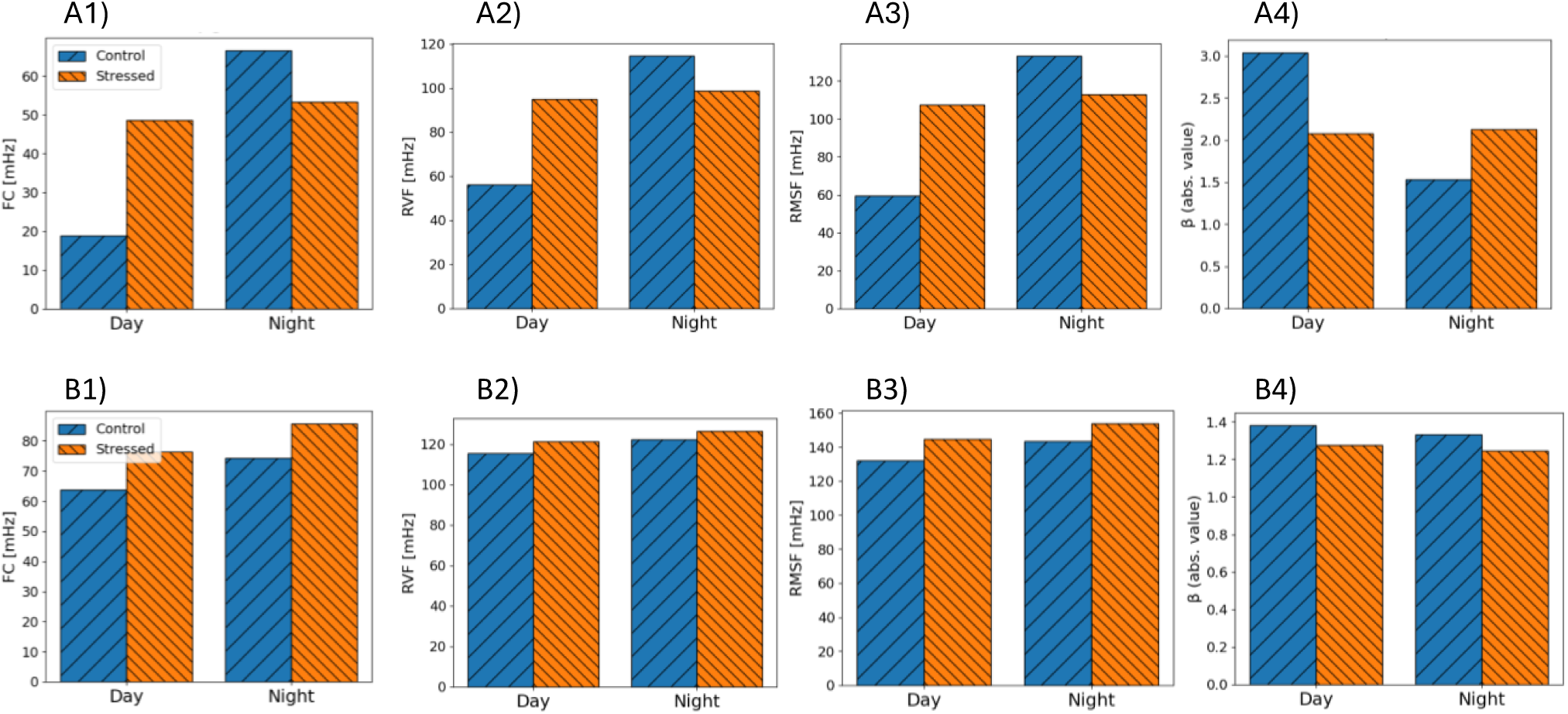
Average frequency features values of the tomato (A) and the apricot (B) plants, shown separately for control and stressed plants, and for daylight and nighttime periods. The frequency features are FC (1), RVF (2), RMSF (3) and β (4).

Regarding PSD values for tomato plants, a visual analysis shows that the amplitude decreases when the frequency increases, following a power law (see Figure 7 and the figures in SM3). Furthermore, during daylight periods control plants present a sharper decrease in the amplitude when increasing the frequency components (they have a higher absolute β value) than the drought-stressed plants. In fact, when considering the data for the day segments, β is lower in the control than in the drought-stressed plants. Instead, when only the night segment is considered, the behaviour is inverted (Figure 6A4). At night, the absolute value of β for the control set significantly decreases, with a value that is 50% lower than that obtained for the daylight. The value of β shows a light non-significant increment for the drought-stressed signals. The frequency features of the apricot plants show only small differences between control and stressed plants, both during days and during night (Figure 6B1-6B4).

**Figure 7:**
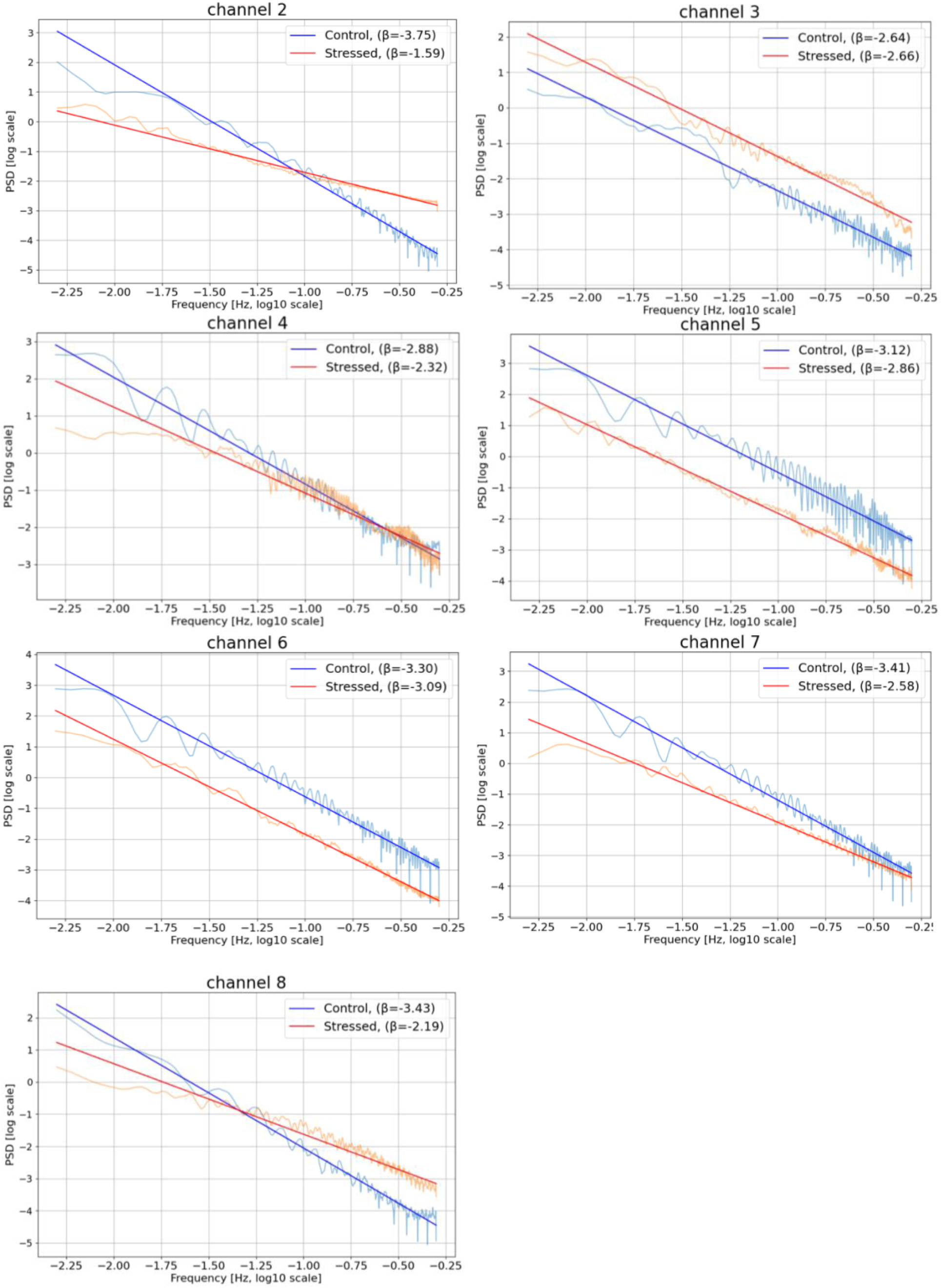
Power Spectral Density of the tomato plants for the signal of the 03/11 day. The scale is bilogarithmic, so a straight line corresponds to a power-law relation between the PSD and the frequency. The images for the other periods are in SM3.

### 3.2 Performance of logistic and machine learning models

For the tomato plants, the logistic model reached an accuracy of 67%, but with a high variability (standard deviation equal to 29%). As shown in Table 2, the precision is higher than the recall, meaning that the model would provide more false negatives (stressed state classified as healthy) than false positives (healthy state classified as stressed). The median is higher than the average, while the interquartile range is lower than the standard deviation. The reason is that one of the channels, when used for validation, gave an extremely low value of accuracy (2%), thus lowering the average and increasing the standard deviation. The other channels gave values between 54% and 98%. Among all the channels, the interquartile range, supported by the standard deviation, describes a higher variability in the recall respect to the precision. The features that provide the highest contributions are the Maximum and the Minimum, followed by Hjorth Complexity and Generalized Hurst Exponent (Figure 8). The performance of the classifier does not decrease when these variables only are used.

**Figure 8.**
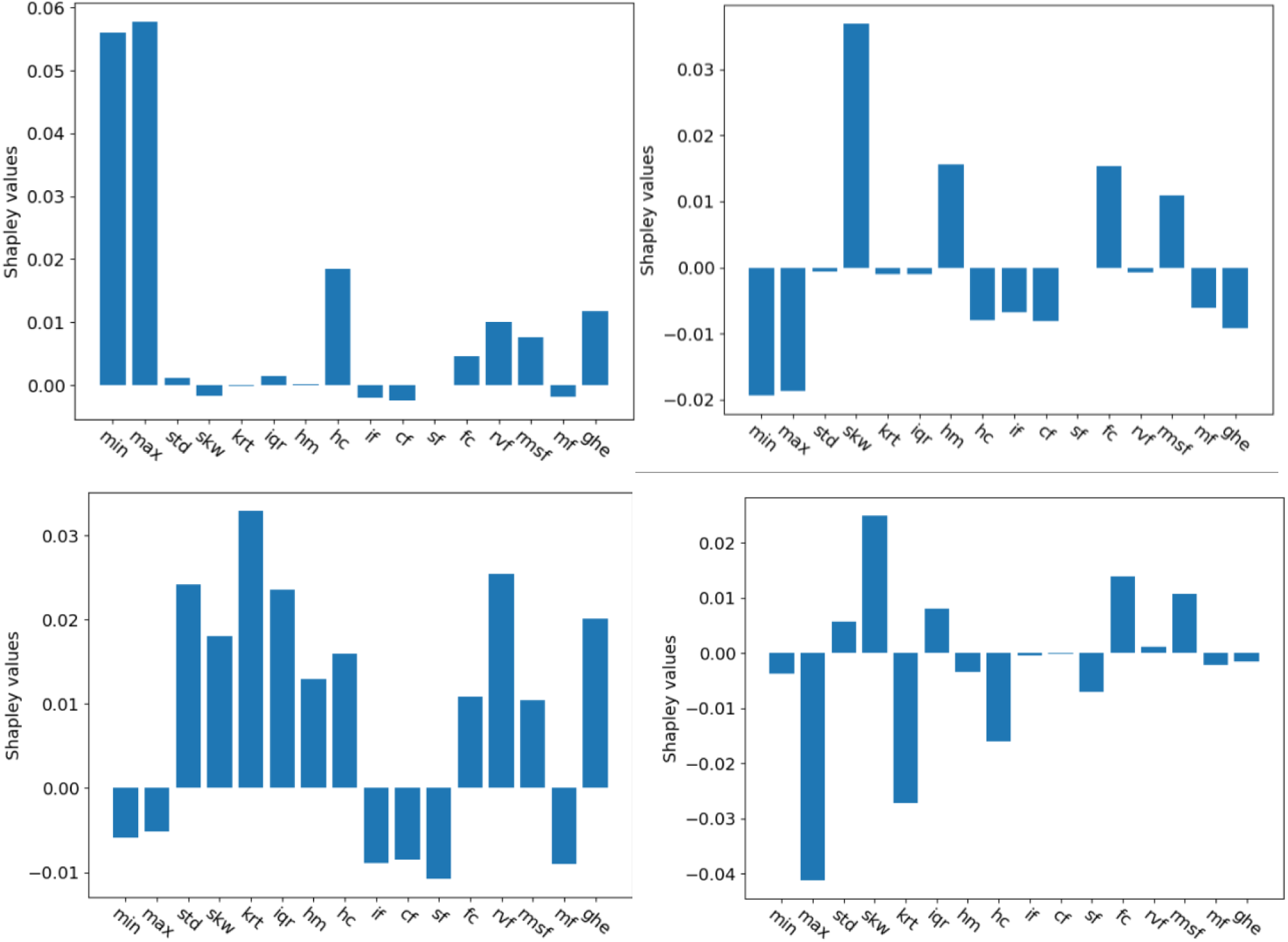
Shapley values of each feature as a regressor in the logistic classifier (top row) and in the XGB model (bottom row). The plots on the left shows the results for the tomato experiment, while the plots on the right show those for the apricot experiment. The Shapley values are averaged across all the trials of the cross-validation.

**Table 2.**
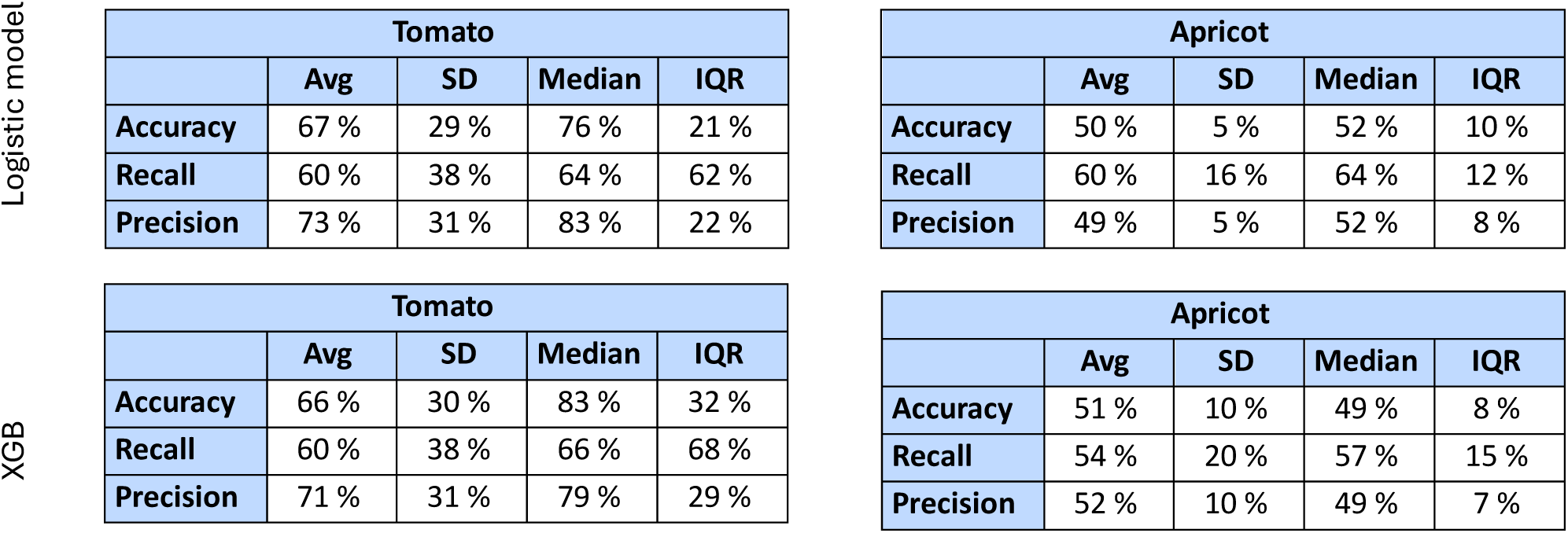
Classifier outputs aggregated by two different metrics, average (Avg) and median, with their dispersion indexes, standard deviation (SD) and interquartile range (IQR). The classifiers are the logistic model and the XGB, which employed the hyperparameters N = 15 and l = 120 for the tomato and N = 30 and l = 660 for the apricot. See SM 4 for the results for further values of the hyperparameters.

For the apricot plants, instead, the logistic model reached an average accuracy of 50% only, with a small variability (standard deviation equal to 5%). This means that the model barely overcomes the performance of a random choice, and in some cases, it performs worse than it. As shown in Table 2, the recall is higher than the precision, meaning that the model would provide more false positives (healthy state classified as stressed) than false negatives (stressed state classified as healthy). The feature with the largest Shapley value is the Skewness, followed by the Hjorth Mobility and the Frequency Center (Figure 8).

The results of the XGB depend on the choice of its hyperparameters, namely the size of the windows *l* and the number of windows employed *N*. Figure 9 shows the average accuracy across the leave-one-out validations and its standard deviations. The Pareto frontier is also highlighted in the figure. A higher standard deviation means that the performance of the XGB model is sensitive to the specific channel used for the validation. The best performance is the one with the highest accuracy and the lowest standard deviation.

**Figure 9.**
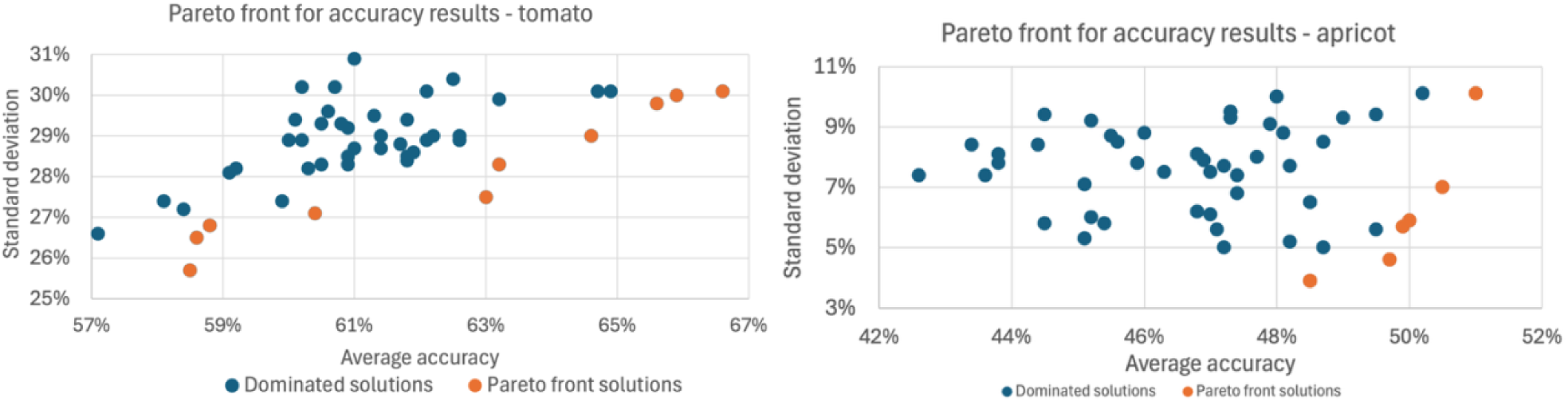
Accuracy results of the XGB models for varying values of N and l. The Pareto front highlights non-dominated solutions. The best points are those nearer to the bottom right edge of the graph.

The solutions with the highest accuracy were analysed more in depth. For the tomato plants, this solution had hyperparameters *N* = 15 and *l* = 120, while for the apricot plants, it had *N* = 30 and *l* = 660. For these solutions, more performance metrics are reported for this model in Table 2.

In summary, the model for the tomato plants reaches an accuracy that, although not optimal, is significantly better than random chance. The precision appears to be significantly higher than the recall, meaning that the model is more prone to classify the stressed state as healthy than the opposite. The model for the apricot plants, instead, barely reaches an accuracy of 50%, similar to the logistic model. Regarding the contributions of the features, the KRT, STD, IQR, and RVF are the most impactful for the tomato’s model’s learning process, as shown in Figure 8. The contributions to the apricot model are not relevant, as the model is not effective in discriminating between control and stressed plants.

## 4 Discussion

This study showed that drought-stressed and healthy plants exhibit differences in their EPS, highlighting the potential of EPS analysis as a valuable emerging technique for monitoring plant drought stress. In particular, from the comprehensive set of statistical features, FC, HM, HC, RVF, RMSF, and GHE consistently showed statistically significant differences between control and stressed apricots. For apricots, Figures 2–3 clearly illustrate that for specific features (e.g. FC and RMSF), the divergence in values between control and stressed plants increased progressively with wilting. In some instances, this trend was accompanied by greater variability in feature values among the stressed plants compared to the controls, as shown in Figure 5. Instead, drought-stressed and healthy tomato plants showed different values for KRT, RVF, and GHE (see Figure 1), although less consistently than apricots. Regarding frequency domain features, tomatoes showed significantly different values for all of them (FC, RVF, RMSF and β) when computed on the daylight periods, but not on the nighttime periods. Lastly, two classifiers, the logistic regression and the XGB, were not effective in discriminating segments coming from either stressed or control apricot plants, as they reached an accuracy of approximately 50%. The same classifiers yielded better results for tomatoes, with an accuracy of approximately 66%, though this level of performance remains inadequate for practical applications. In terms of the contribution of specific features to the classifiers, MIN, MAX, and KRT emerged as the most influential variables. This observation aligns with previous findings (Tran et al., 2019b), (Najdenovska et al., 2021d) which similarly demonstrated that classifiers could distinguish between stressed and control tomato plants, albeit with somewhat higher accuracies.

### 4.1. Time– and frequency-domain-features: statistical significance and ecological relevance

In terms of time-domain features, the results suggest that FC, RVF, RMSF and GHE computed on the 400-seconds segment can discriminate drought-stressed and control plants, but they might not be effective metrics if the stress is not severe enough. FC, RVF and RMSF reflect whether higher or lower frequencies are prevailing in the plants. In particular, natural phenomena generally have a larger power density in lower frequencies, while a random noise has a constant power density across frequencies, so a shift in the plant’s prevailing frequency might be due to the reduction in the intensity of its physiological processes with respect to the baseline noise. GHE indicates whether a signal has short-term correlations only or also long-term ones. Short-term correlations are plausible in a plant constantly responding to the environment, while long-term correlations are plausible in a plant that suffered a loss of adaptation to the environmental conditions and is focused on the processes related to drought resistance. The importance and the direction of change of GHE confirms the observation of a previous study on tomatoes under drought stress (Tran et al., 2019b). Moreover, both GHE and FC were reported as significant in another study involving tomatoes exposed to multiple stressors, including drought (Najdenovska et al., 2021d). These features have also been identified as relevant in studies examining other stress conditions in tomatoes, such as infestations by spider mites (Najdenovska et al., 2021b) and nutrient deficiencies (Tran et al., 2024). In contrast, a previous study (Chatterjee et al., 2015b) reported that exposure of tomato plants to three different stressors (ozone, sulphuric acid, and osmotic stress) consistently led to a decrease in GHE, unlike the increase observed in the present study. Further studies are needed to clarify whether this difference reflects a distinct impact of drought stress on the plant’s EPS compared with the other stressors examined in (Chatterjee et al., 2015b).

Among the time-domain features with good discriminative power, FC, RVF, and RMSF exhibited greater variability among stressed apricots than among controls, as showcased in Figure 5. This pattern suggests that, within the stressed group, some plants displayed markedly higher feature values as wilting advanced, whereas others maintained responses similar to those of the controls. Such heterogeneity could account for both the observed differences between groups and the increased variability. These findings imply that EPS-based monitoring systems may be less effective for individual-plant diagnostics, particularly for certain features, but could perform well in detecting potential collective stress responses at the population level. Accordingly, deploying EPS sensors across multiple plants under similar conditions (e.g., within the same field, greenhouse, or urban green patch) may enhance their capacity to detect environmental stress. In addition, coupling EPS recordings with complementary approaches, such as visual or physiological assessments, could further improve interpretability.

Regarding the frequency-domain features, the results, illustrated in Figure 6, showed significant differences between control and stressed tomato plants during the daylight periods. Control plants exhibited lower FC, RVF, and RMSF values and a higher β coefficient during the day compared with stressed plants. However, at night, control plants displayed feature values similar to those of the stressed plants, which maintained comparable values across both day and night conditions. This pattern suggests that, while control plants modulate their electrophysiological activity between daylight and nighttime periods, i.e., by shifting their frequency spectrum toward lower frequencies, stressed plants lose this diurnal variability and maintain a relatively constant spectral profile. In other words, the key difference lies not in the absolute feature values but in the dynamic behavioural change that is preserved in control plants and seems to be suppressed under stress. These findings contrast with those reported by Simmi *et al*. (Simmi et al., 2020b), who observed a slightly higher β coefficient in stressed plants during daylight. However, their experiment involved tomato plants infected by *Oidium neolycopersici*, suggesting that changes in β coefficient values may depend on the specific nature of the stressor, and values for this feature should therefore be interpreted in a stressor-specific context. Moreover, when applied to apricot plants, no significant difference in frequency-domain feature values were detected, either during the day or at nights. Nevertheless, a previous study on another woody species, grapevine, reported a higher proportion of power at lower frequencies in control plants (Cattani et al., 2024b), consistent with the results obtained for tomatoes in the present study. Further studies are needed to replicate the analysis on apricots and to evaluate the extent to which frequency-domain patterns can be generalized across different species. Therefore, the β coefficient may exhibit plant-specific and stressor-specific behavior, and its reliable interpretation, and potential utility for practical monitoring, likely depends on first identifying the prevailing stressors, either through variations in other electrophysiological features or by integrating EPS analysis with complementary diagnostic approaches.

In terms of the methodological aspects of feature analysis, the effectiveness with which features differentiate stressed from control plants in this study also depends on the statistical test applied. For instance, Figure 3 shows that the separation between HC values for stressed and control plants appears abrupt when evaluated using the Mann–Whitney test, whereas the FDR reveals a more gradual and less distinct variation. This result highlights the importance of applying multiple statistical tests when comparing features across plant groups, as different methods may capture different aspects of their discriminative capacity. In contrast with our study, previous studies have employed ANOVA (Bhadra et al., 2023; Costa et al., 2023) and the Wilcoxon signed-rank test (de Toledo et al., 2024; Pereira et al., 2018b). However, ANOVA assumes normally distributed data, which was generally not the case in the present experiments. On the other hand, the Wilcoxon signed-rank test is appropriate for dependent samples and could be used to assess variations in feature values before and after a stimulus. However, in the current study two independent sets of plants were compared, control and stressed, making the Mann–Whitney test more suitable. Overall, the use of two different statistical tests in this study reduces the risk of overlooking relevant patterns. A potential future improvement would be to compare stressed plants before and after wilting using also Wilcoxon signed-rank test.

### 4.2. Logistic and machine learning classifiers: importance for drought detection tools

Both logistic regression and XGB achieved similar performances, in discriminating signal segments from drought-stressed and healthy plants, with accuracies of approximately 66% for tomatoes and 50% for apricots, as shown in Table 3. For apricots, this value is comparable to random guessing, indicating that the classifiers failed to capture distinctive electrophysiological patterns between control and stressed plants. Despite the accuracy for tomato plants is better than random chance, it is still insufficient for a good classifier. When compared with other EPS studies on tomato plants that also employed classifiers (Chatterjee et al., 2014; González I Juclà et al., 2023; Kurenda et al., 2024; Najdenovska et al., 2021a, 2021c; Tran et al., 2024, 2019a; Tran and Camps, 2021), the performance observed here is lower. However, these results also highlight an opportunity for improvement, proving that EPS-based classifiers have a potential for applied use as stressor detection tools. Additionally, as discussed in Section 4.1, these outcomes may partly reflect plant-specific and stressor-specific behaviors that influence the discriminative power of EPS-based models. Alternatively, the results might be affected by the noise filtering approach used. In particular, the apricots’ signals were affected by a disturbance that recurred at regular intervals and which was removed and substituted by imputed values. The imputation might have altered the signal characteristics, thus lowering the possibility of recognizing the stress pattern. Another explanation for the low accuracy might involve the number of features, which was 16, while other studies employed a larger number. For instance, (Tran et al., 2019b) used 182 features for the detection of drought stress in tomatoes. Future research should aim to clarify these factors and to identify the key methodological aspects, such as optimal feature selection and classifier design, that could substantially enhance the effectiveness of EPS-based monitoring for different plant species and stress conditions.

In the logistic model, the features MIN and MAX were the major contributors to the classification, even though the statistical analysis of individual features did not identify them as significant. In contrast, the XGB model relied on KRT, which showed relevance in the statistical analysis within the time domain (see Figure 1). Despite this apparent inconsistency, similar outcomes were reported by Tran et al. (Tran et al., 2019b), whose classifiers also identified MIN and MAX as key contributing features. This may suggest that the relationship between these features and plant status is complex and possibly involves interactions between them, patterns that cannot be detected through simple univariate comparisons between control and stressed plants. Moreover, the accuracy obtained was highly variable depending on the channel used for validation, which may indicate that the machine learning models captured trends applicable to most plants, while a few exhibited divergent behaviors. This behavior is observed in the greater variability of FC, RVF, and RMSF in stressed apricots (Figure 5), which further supports the idea that the electrophysiological responses among drought-stressed plants are more heterogeneous than those of control plants.

Other relevant features for the classifiers included HC and GHE for the logistic model, and RVF, STD, and IQR for the XGB model. Two of these features, GHE and RVF, also showed statistically significant differences between control and stressed tomato plants, although not in a consistent way across the experiments. On the other hand, all features identified as significant in the statistical analysis (KRT, GHE, and RVF) were also among the most relevant variables for the classifiers. It is likely that the models were able to discriminate between control and stressed plants primarily on the days when these features exhibited clear separation between the two groups, thereby slightly improving accuracy. This finding confirms that KRT, GHE, and RVF are indeed relevant features, but their potential for drought-stress detection remains limited.

### 4.3. Limitations and suggestions for future studies

Despite the value of this study in advancing the understanding of EPS analysis as an emerging technique for monitoring plant drought stress, and potentially other stressors, the experiments had certain limitations that partially constrained the analysis and comparison of the EPS data.

Firstly, the experiment on apricots was conducted outdoors, but apart from visual observations, no additional environmental variables were used in the analysis. Such data could help to interpret variations in feature values and to explain some observed dynamics. Secondly, the EPS signals from both experiments were affected by noise. Although filtering was applied to reduce it, this process may have inadvertently altered aspects of the original signal and, consequently, the derived feature values. Thirdly, the tomato dataset covered a shorter time period than the apricot dataset, which prevented assessment of how feature separation evolved over time, as was done for apricots. This limited the comparability between the two experiments and hindered the identification of potential EPS dynamics that might be generalizable across species under drought stress.

Future pilot studies aiming to detect plant drought stress through EPS, whether in laboratory or field conditions, should aim to minimize these limitations by incorporating complementary sensors to monitor abiotic factors such as radiation, temperature, and humidity; by verifying signal quality throughout data collection; and by ensuring that experiments are of sufficient duration to capture the full transition from healthy to severely wilting states.

## 5 Conclusion

This study identified the electrophysiological signal’s (EPS) features that are most consistently associated with drought stress in tomato and apricot plants, highlighting their potential as key metrics for future EPS-based early-warning tools.

In tomatoes, frequency-domain features, including Frequency Center (FC), Root Variance Frequency (RVF), Root Mean Squared Frequency (RMSF), and the β coefficient, consistently differentiated stressed from control plants during daylight periods. Time-domain features such as Generalized Hurst Exponent (GHE) and Kurtosis (KRT) also showed potential relevance, albeit less consistently. For apricots, several time-domain features, including FC, RVF, RMSF, GHE, Hjorth Mobility (HM), and Hjorth Complexity (HC), changed significantly as wilting progressed, with stressed plants displaying higher variability than controls. This heterogeneity suggests that dynamic changes rather than absolute feature values may be most informative for detecting drought stress. Some features, such as MAX, MIN, and KRT, were major contributors within classifiers, despite the limited significance of MAX and MIN in univariate statistical analyses, highlighting potential complex interactions between features. Overall, the evidence within this study indicates that the most robust features for detecting drought stress seem to be those that capture dynamic, frequency-based, and distributional changes in EPS signals.

Overall, these results expand the evidence base for EPS as a monitoring tool and lay the groundwork for a drought-stress early-warning system capable of tracking vegetation status in diverse contexts. Such a system could support decision-making in agroecosystems and urban green spaces, providing a practical, scalable, and field-applicable approach to proactive plant monitoring.

Future studies should gain a deeper understanding of the biophysical and ecological mechanisms underlying potentially relevant EPS features, to clarify why the features mentioned above emerge as statistically significant and whether they reflect general stress responses or are specific to drought. At the same time, to ensure robustness and generalizability, EPS analyses should be replicated across additional datasets, plant species, and open-field conditions. In the long term, complementary studies should explore the effects of combined biotic and abiotic stressors, ultimately assessing the feasibility of developing a plant-based environmental monitoring network that leverages EPS signals as early-warning tools for multiple stressors.

## 6 Supplementary materials

Supplementary Material 1: Description of the EPS data pre-processing steps

Supplementary Material 2: Description of the General Hurst Exponent

Supplementary Material 3: Additional results on the time-domain and frequency-domain features

Supplementary Material 4: Additional results for the XGB model

## 7 Acknowledgement

This work was supported by the National Biodiversity Future Centre (NBFC) project, funded by the European Union’s NextGenerationEU, National Recovery and Resilience Plan (NRRP), CN00000033, CUP, D43C22001250001.

