## Supplementary Material 1 for "Electrophysiological monitoring of plants: an exploratory study on drought stress"

**Supplementary Material 1 – Description of the EPS data pre-processing steps.**

Before calculating the features, the EPS data was pre-processed following established steps: i) downsampling, ii) noise filtering; iii) outlier detection and removal; and iv) missing data imputation.

While the original data was collected at a sample rate of 256 Hz, the supplier provided pre-processed data downsampled to 1 Hz for the sample of the tomato experiment, but not for the apricot experiment. Downsampling to 1 Hz reduces data volume and computational load without losing significant information. For this reason, a downsampling algorithm was developed and applied to the second dataset. Before applying the downsampling a low pass anti-aliasing filter was applied to prevent bias derived from changing the sampling rate. The algorithm works by dividing the original signal into windows of the same amplitude as the sample rate. For each window the mean value is evaluated bringing the sample rate equal to 1Hz. Further methods for downsampling can be contemplated such as considering the median value instead of the mean or applying a decimation approach where only the first value of each window is taken. In this study, the mean method appeared as the best solution because it made the signal more stable with less spikes.

The apricot plant dataset showed regular peaks, order of magnitudes larger than the rest of the signal (Figure SM1.1). They were caused by a disturbance, possibly caused by the periodic activity of another device which was located near the experiment's site. Outliers were identified as those exceeding the average of the signal of at least 5 times the standard deviation. The average and the standard deviation were computed in a 90-seconds window around the suspected outlier. However, since the spikes lasted a few seconds, the window ignored the 40 adjacent seconds to the suspected outlier. The detected outliers were replaced with a normal random variable of the same mean and standard deviation of the data points in the window, clipped to 1.7 standard deviations over or below the mean, to avoid creating new outliers.

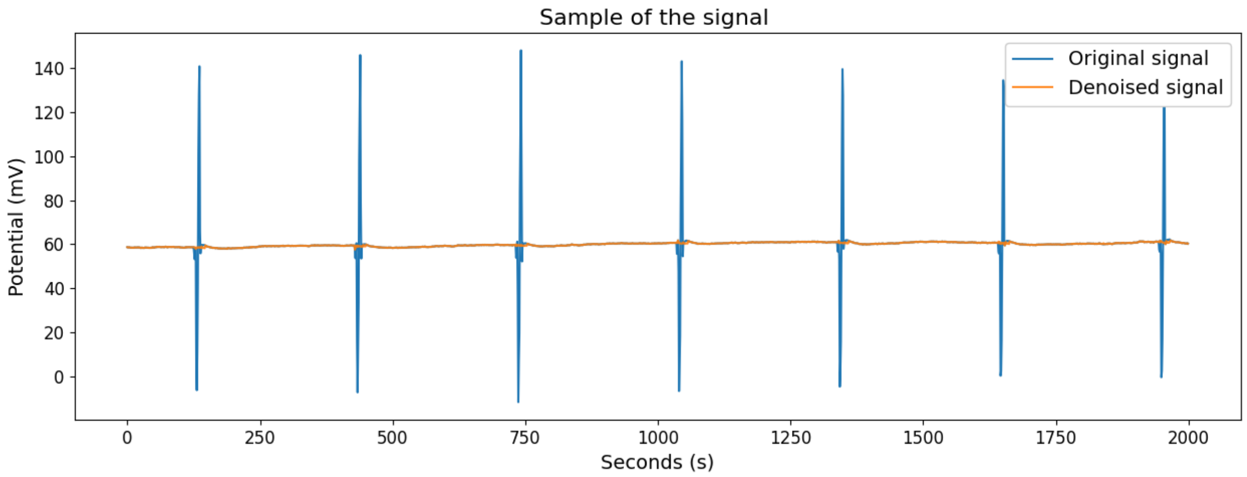


*Figure SM1.1: sample of signal from the apricot plant, both before and after denoising. Note that the noisy peaks are order of magnitude larger than the variations of the signal.*

Afterwards, a notch filter was applied to remove 50 Hz and 100 Hz noise from local power sources. In case there is still noise in the signal a Fast Fourier Transform (FFT) can be performed to detect and eliminate those frequencies. This procedure showed that the control plants in the tomato experiment showed sharp peaks in the frequency domain at approximately 0.05 Hz and upper harmonics. These peaks are too sharp to be plausibly attributed to plants' physiological processes and are likely due to a disturbance during the recording phase. This noise was removed through selective notch filters at the noise base frequency and at its harmonics. Figure SM1.2 shows the impact of the filter on the power spectral density of a signal and on a sample of it.

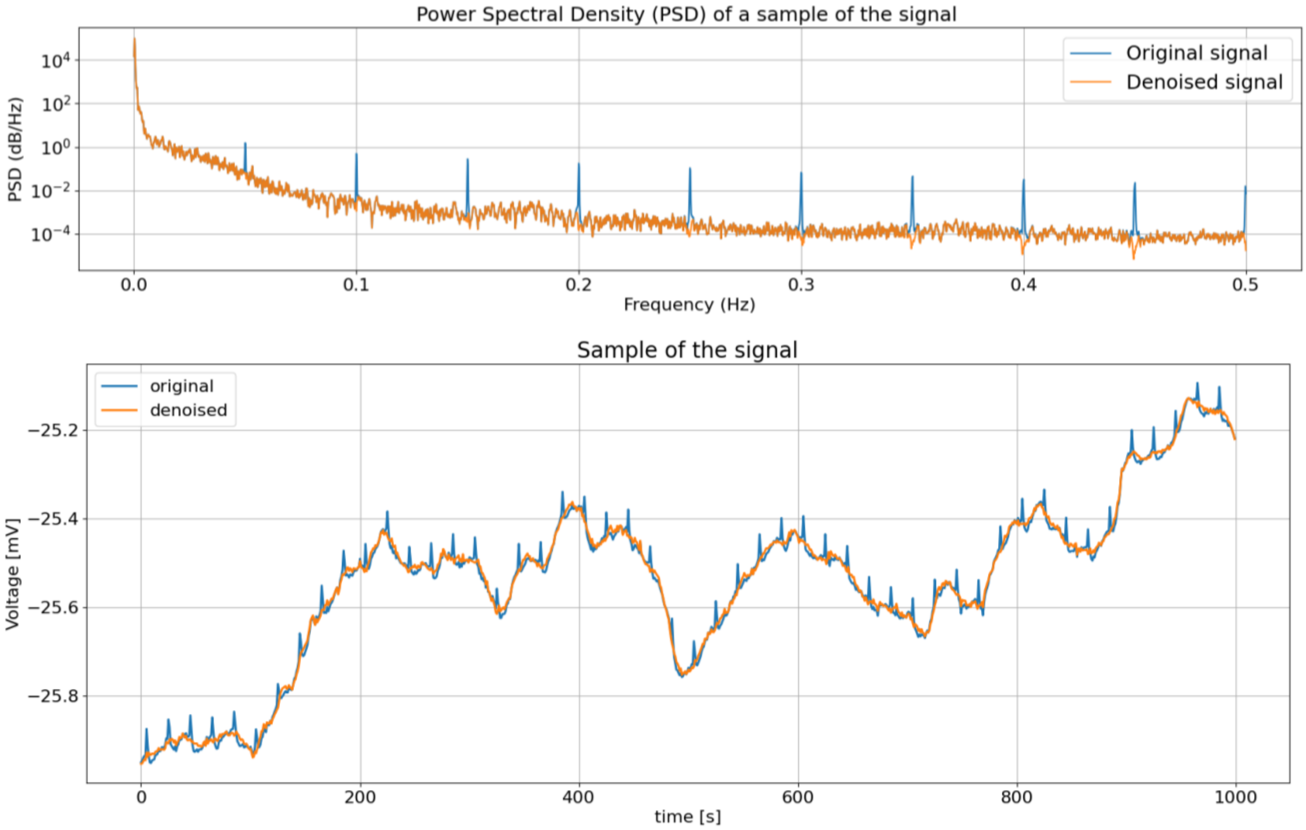


*Figure SM1.2: Impact of the filter of the noise of the tomato control plants. The figure above illustrates the power spectral density, while the figure below shows a sample of the signal.*

The fourth step was the imputation of missing data. The apricot signal had few seconds gaps between the recording of on the day and that of the following one. It also had occasional gaps, at most few minutes long, within each day recordings. Missing data were replaced with a procedure similar to the one used to substitute outliers: each missing point was substituted with a normal random variable that had the same mean and standard deviation as the average and sample standard deviation of 25 consecutive points preceding the missing one, clipped in the same way as was done for the outliers removal.

The tomato dataset had some gaps in the data during the days of 02/11, 04/11 and 06/11. For the control plants, the gaps lasted approximately one minute, while for the stressed plants these gaps lasted approximately one hour. In this case, the gaps were so large that it was not advisable to impute them, as the result would have been strongly affected by the imputation method. Since some analysis, such as those related to the frequency spectrum for a day-long signal, required the completeness of the data, they were restricted only to those days where there was no gap.

After all the preprocessing was completed, it was necessary to determine the optimal window size (*l*) for the analysis of features in the time domain through empirical testing. This should not be confused with making a moving average of the value of the features. Setting a window of size *l*, indeed, means computing one single value every *l* points in the time series and the value of the features are computed based on non-overlapping windows. The moving average, instead, would compute a feature value for each of the points in the time series, and neighbouring feature values would be the averages on largely overlapping windows. After iterative tests, a window length of 400 seconds was considered the most suitable, providing a clear visual output. Figure SM1.3 provides an illustration of different window sizes.

Figure SM1.3 illustrates the effect of window size on a specific feature's visual results.

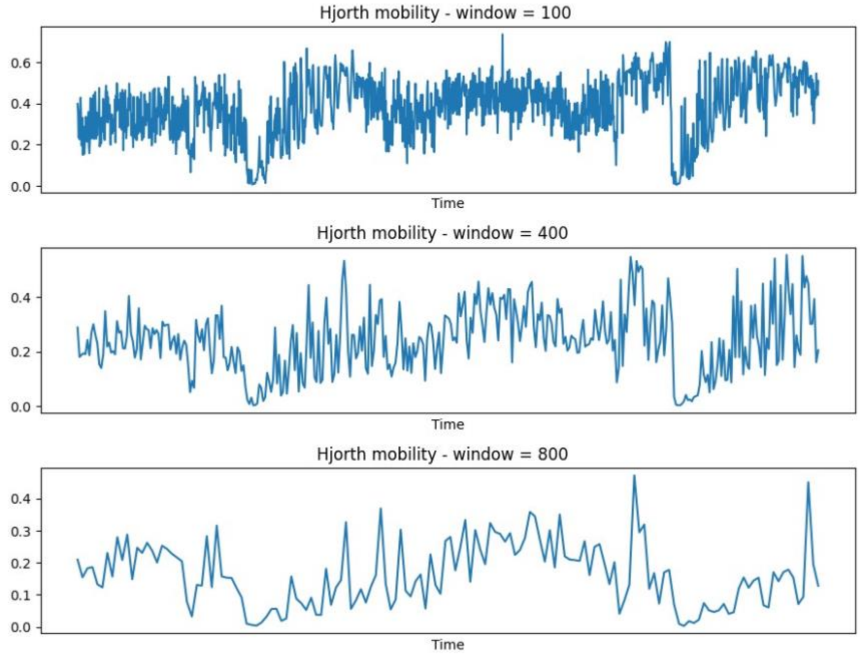


*Figure SM1.3. Visual representation of the effect of the windows length in the values of Hjort Mobility. The values of the windows are in seconds.*
