## Supplementary Material 2 for "Electrophysiological monitoring of plants: an exploratory study on drought stress"

**Supplementary Material 2 – Description of the General Hurst Exponent**

The Hurst Exponent is a powerful index that describes the long-term memory of time series. It evaluates the decrease in autocorrelation as the time lag between two values increases. In fractal geometry, the Hurst exponent is also generalized to account for the fractal dimension's properties. This theory can be adapted to signal analysis to assess the time series long-term correlation.

A common approach for the General Hurst Exponent (GHE) estimation, that we also used in this work, consists in applying the DFA (detrended fluctuation analysis) on the original signal, through the Rescaled Range method.

Firstly, the original time series $X$ is divided into $N$ non overlapping windows of length $n$ ($X^1, X^2, \dots, X^N$). Then, for each window, the average is computed:

$$\mu_k = \frac{1}{n}\sum_{i=1}^{n} X_i^k$$

The following computation is the cumulative deviation series, namely the sum of the deviations from the mean:

$$Z_j^k = \sum_{i=1}^{j}\left(X_i^k - \mu_k\right)$$

Then the rescaled range is computed, that is the range of the first $n$ cumulative deviations from the mean:

$$R_k(n) = \max_{j\in(1,2,\dots,n)}\left(Z_j^k\right) - \min_{j\in(1,2,\dots,n)}\left(Z_j^k\right)$$

The standard deviation of the points in the interval is also computed:

$$S_k(n) = \sqrt{\frac{1}{n}\sum_{i=1}^{n}\left(X_i^k - \mu_k\right)^2}$$

The GHE is thus defined with the following expression:

$$\sum_{k=1}^{N}\frac{R_k(n)}{S_k(n)} = C\, n^{GHE}$$

The averaging operation is performed between all the previously computed non-overlapping windows.
Then, the GHE is estimated by a linear regression on the log-linearization of the reported formula:

$$\log\sum_{k=1}^{N}\frac{R_k(n)}{S_k(n)} = GHE\,\log n + \log C$$

The result for a tomato plant's signal on a whole day is reported in Figure SM2.1. The GHE represents the light-blue line's slope.

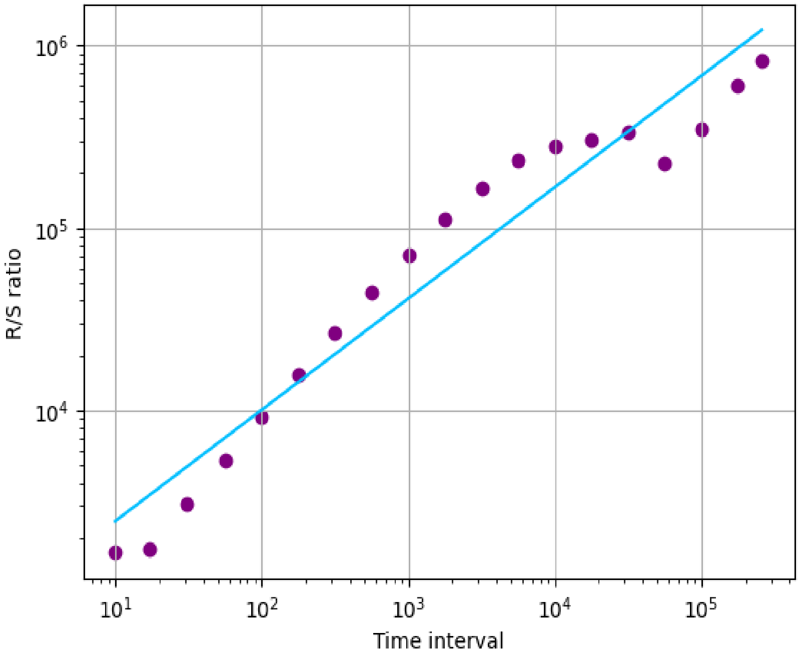


*Figure SM2.1: rescaled range for a tomato plant's signal. The GHE is the slope of the light-blue regression line.*

A random walk has a GHE equal to 0.5 and each increment of the time series is independent of one other. A time series with an anti-persistent behaviour has a lower GHE value, and this means that there's a high tendency of reverting two consecutive values. When the GHE value is higher than 0.5 indicates a persistent behaviour, where an increasing increment has higher probability to be followed by another increasing increment and vice versa. Since the rescaled range is divided by the standard deviation, GHE values are independent from the original signal's amplitude.
