## Supplementary Material 3 for "Electrophysiological monitoring of plants: an exploratory study on drought stress"

**Supplementary Material 3 – Additional results on the time-domain and frequency-domain features**

This SM includes additional results, which were not included in the article because of space reason. This SM is organised into subsections, with the titles following the same structure of those in the Results section:

- SM 3.1: Time domain;
- SM 3.2: Frequency domain analysis .

### SM 3.1 Time domain analysis

#### Tomato plants – time domain

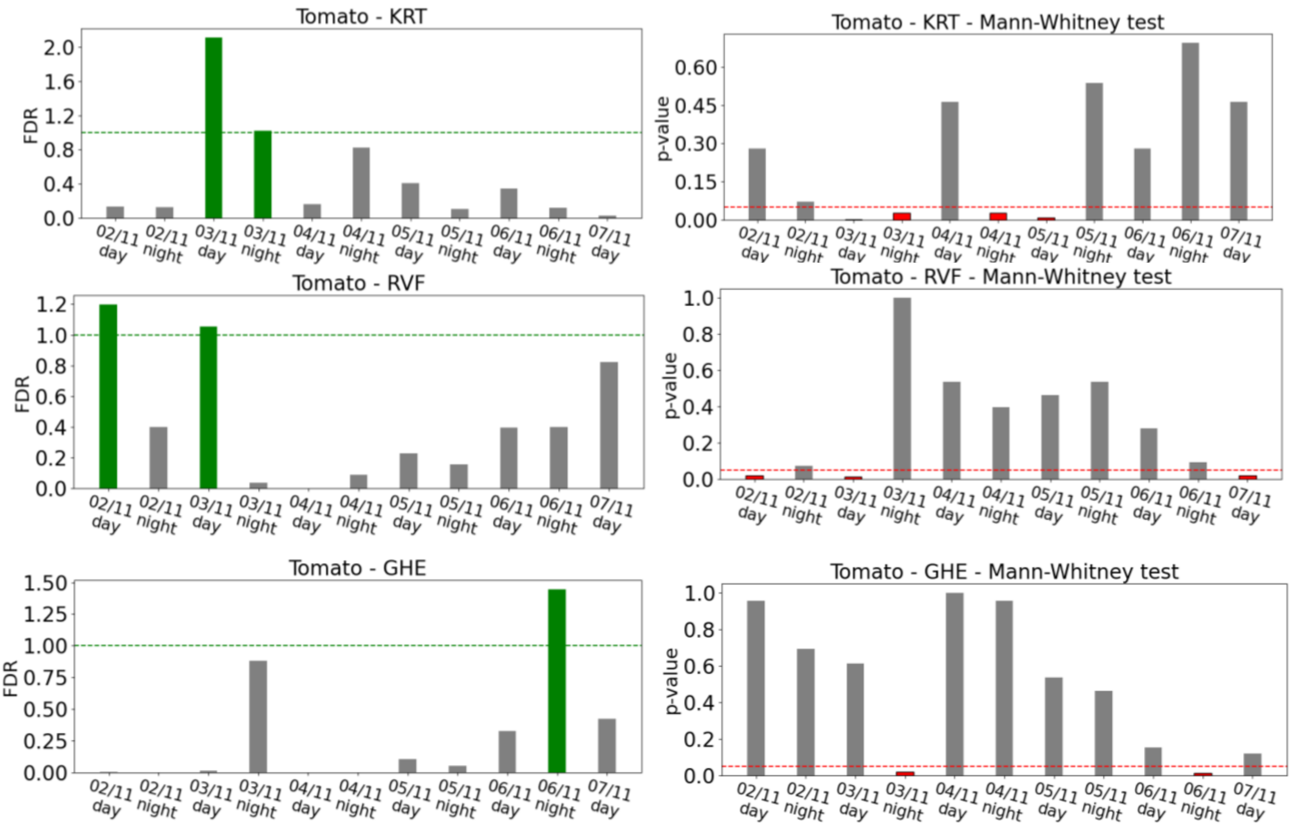


*Figure SM 3.1.1: FDR (on the left column) and p-value of the Mann-Whitney test (on the right column), showing the separation between control and stressed tomatoes, for all the features which exceeded the threshold of one for the FDR.*

### Apricot plants – time domain

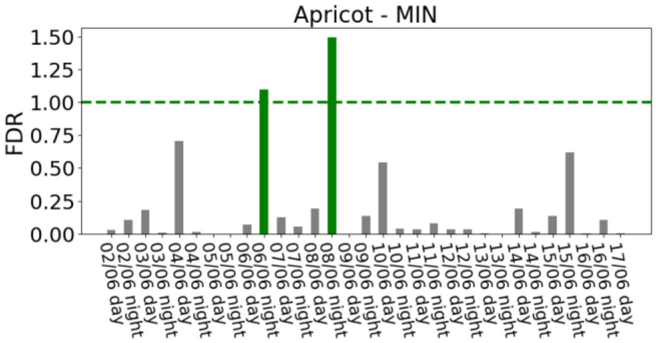


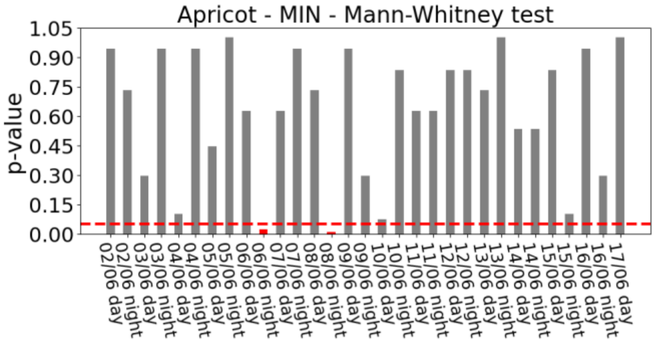


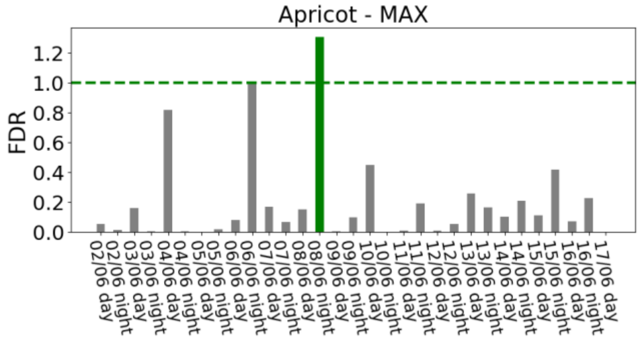


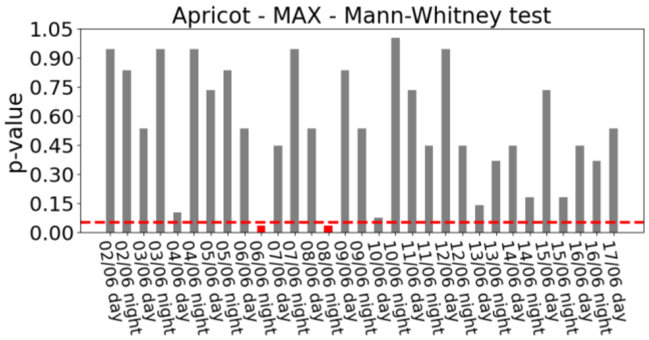


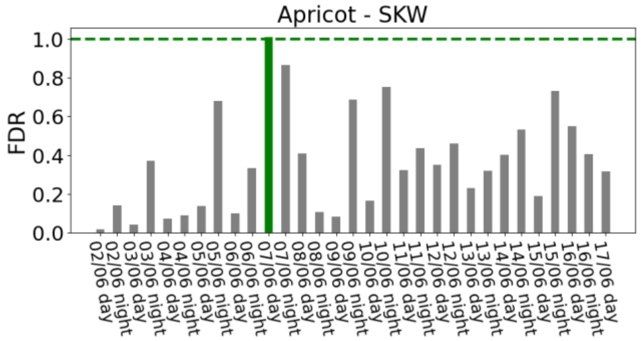


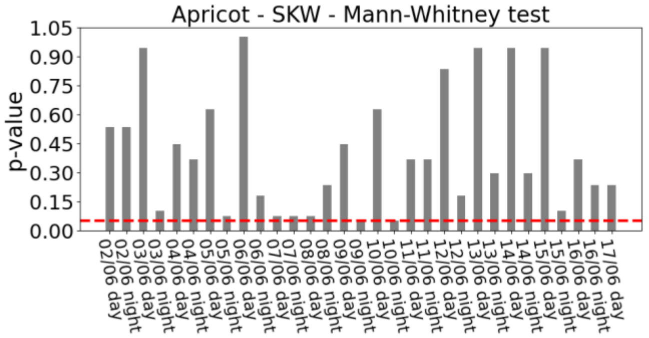


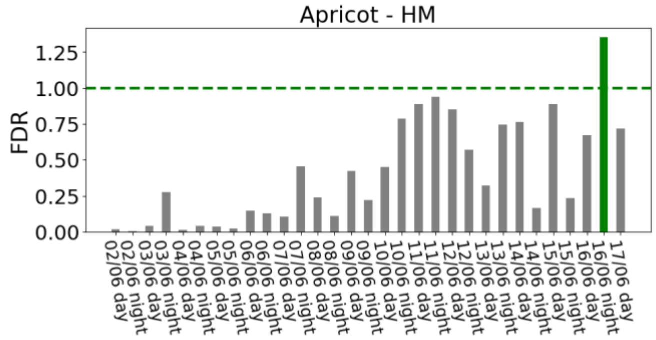


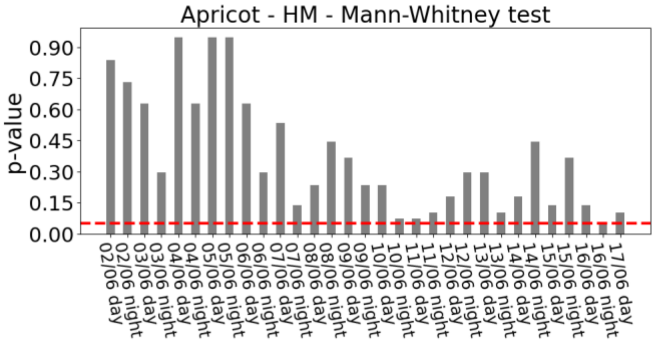

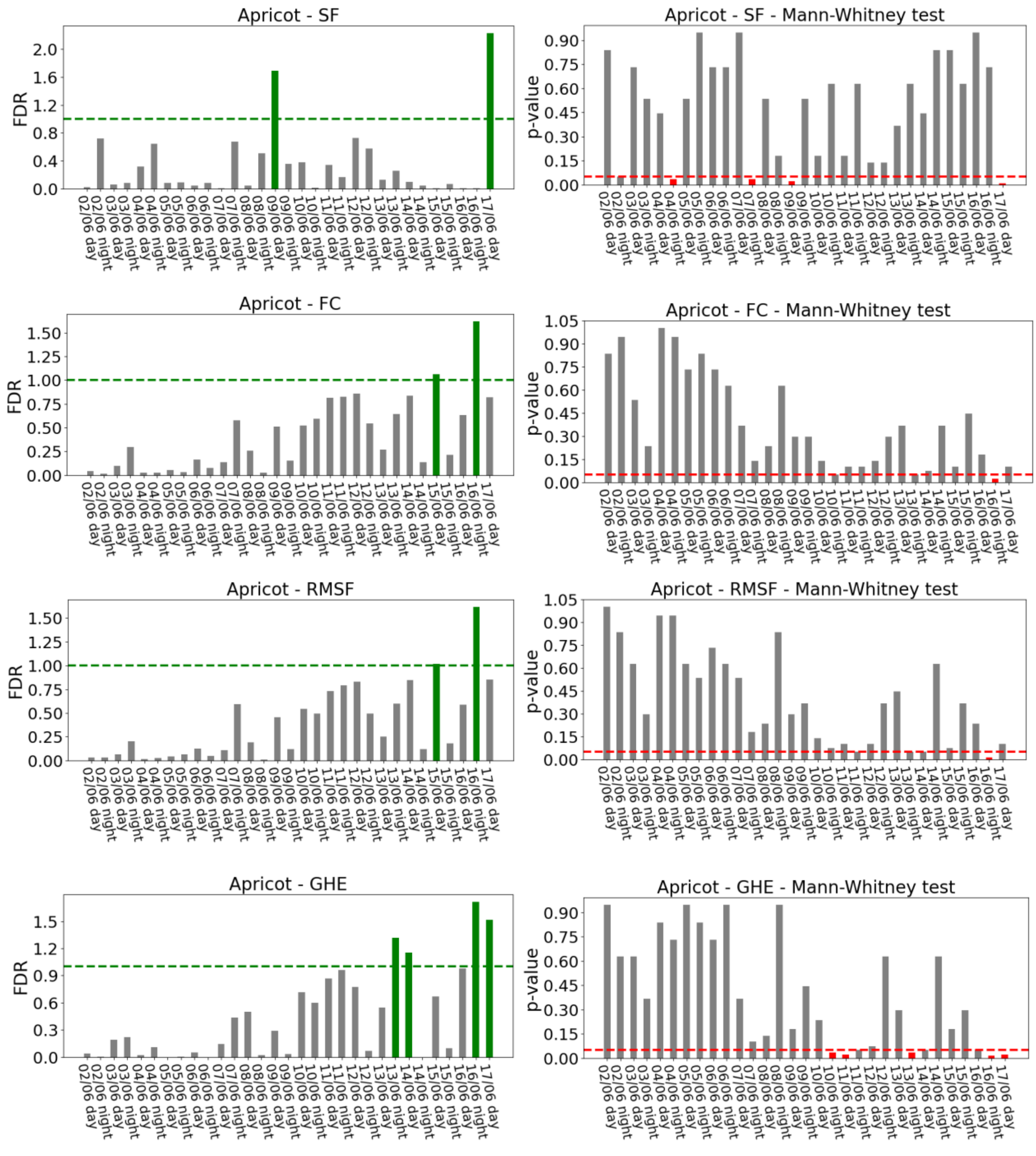


*Figure SM 3.1.2: FDR (on the left column) and p-value of the Mann-Whitney test (on the right column), showing the separation between control and stressed apricots, for all the features which exceeded the threshold of one for the FDR.*

### Apricot plants – comparison around the first appearance of visible signs of wilting

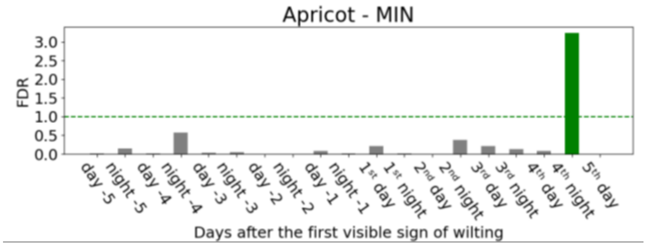


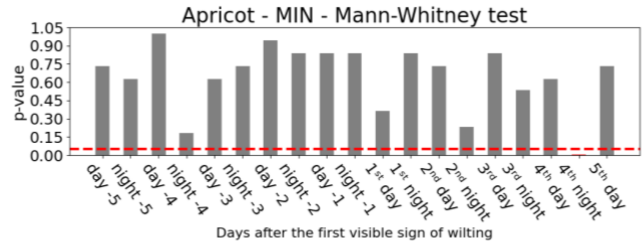


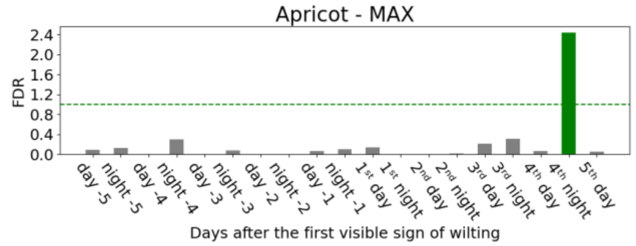


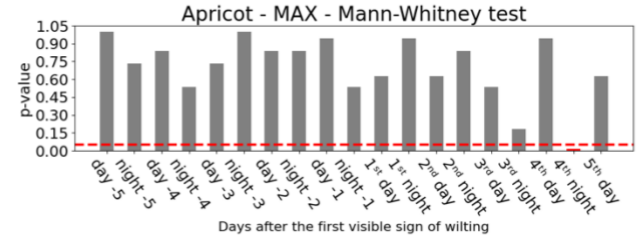


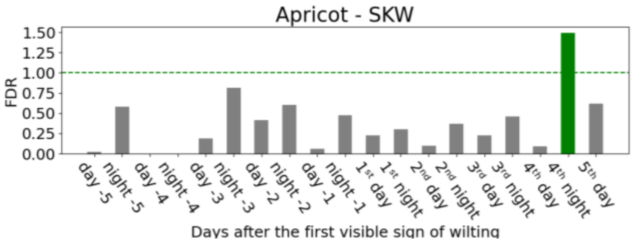


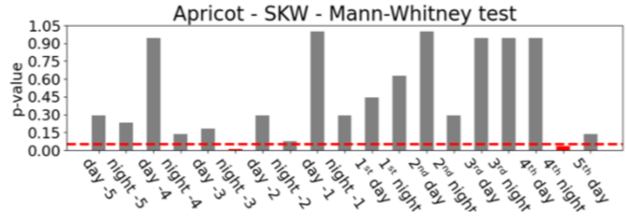


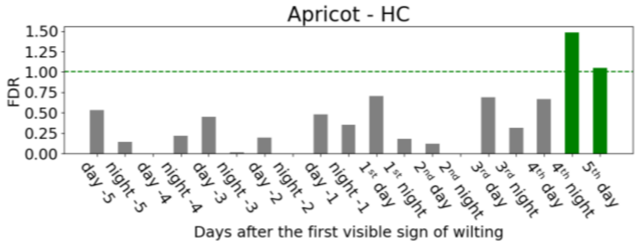


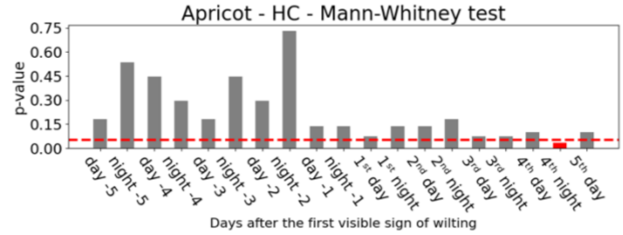


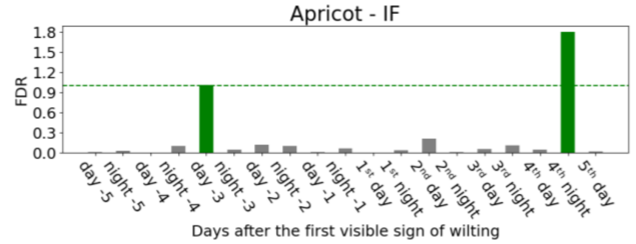


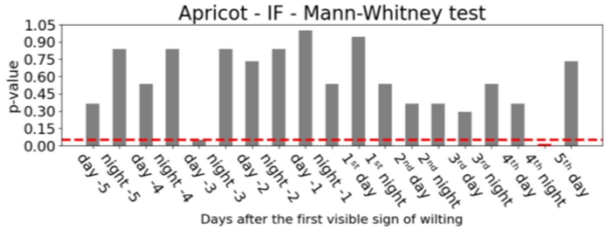


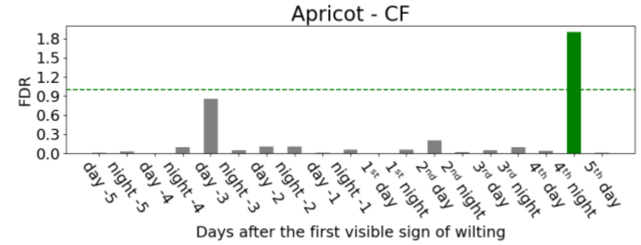


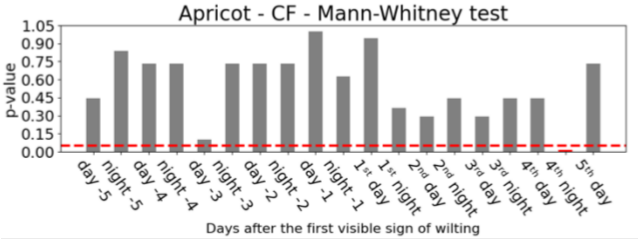

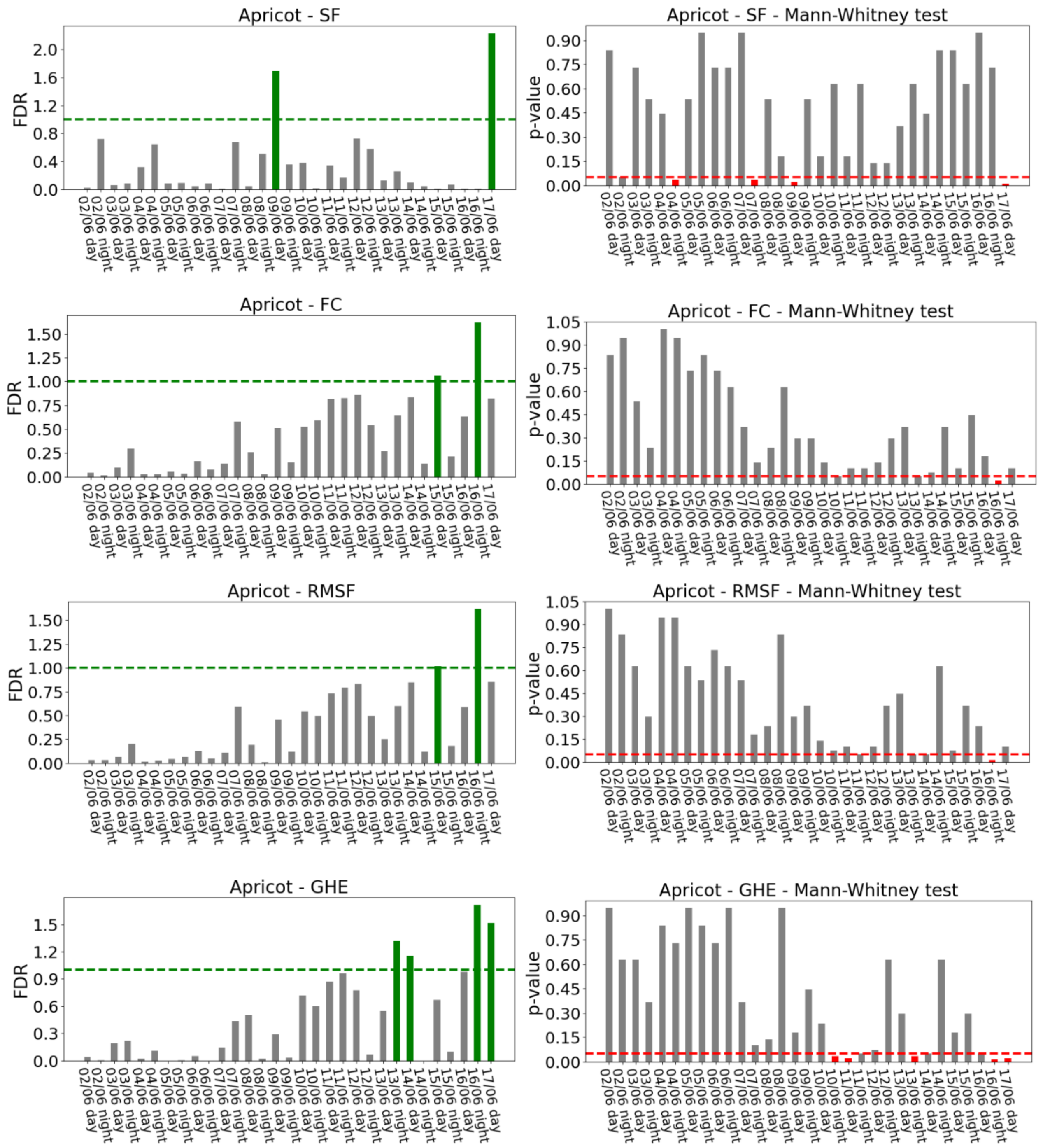


*Figure SM 3.1.2: FDR (on the left column) and p-value of the Mann-Whitney test (on the right column), showing the separation between control and stressed apricots, for all the features which exceeded the threshold of one for the FDR.*

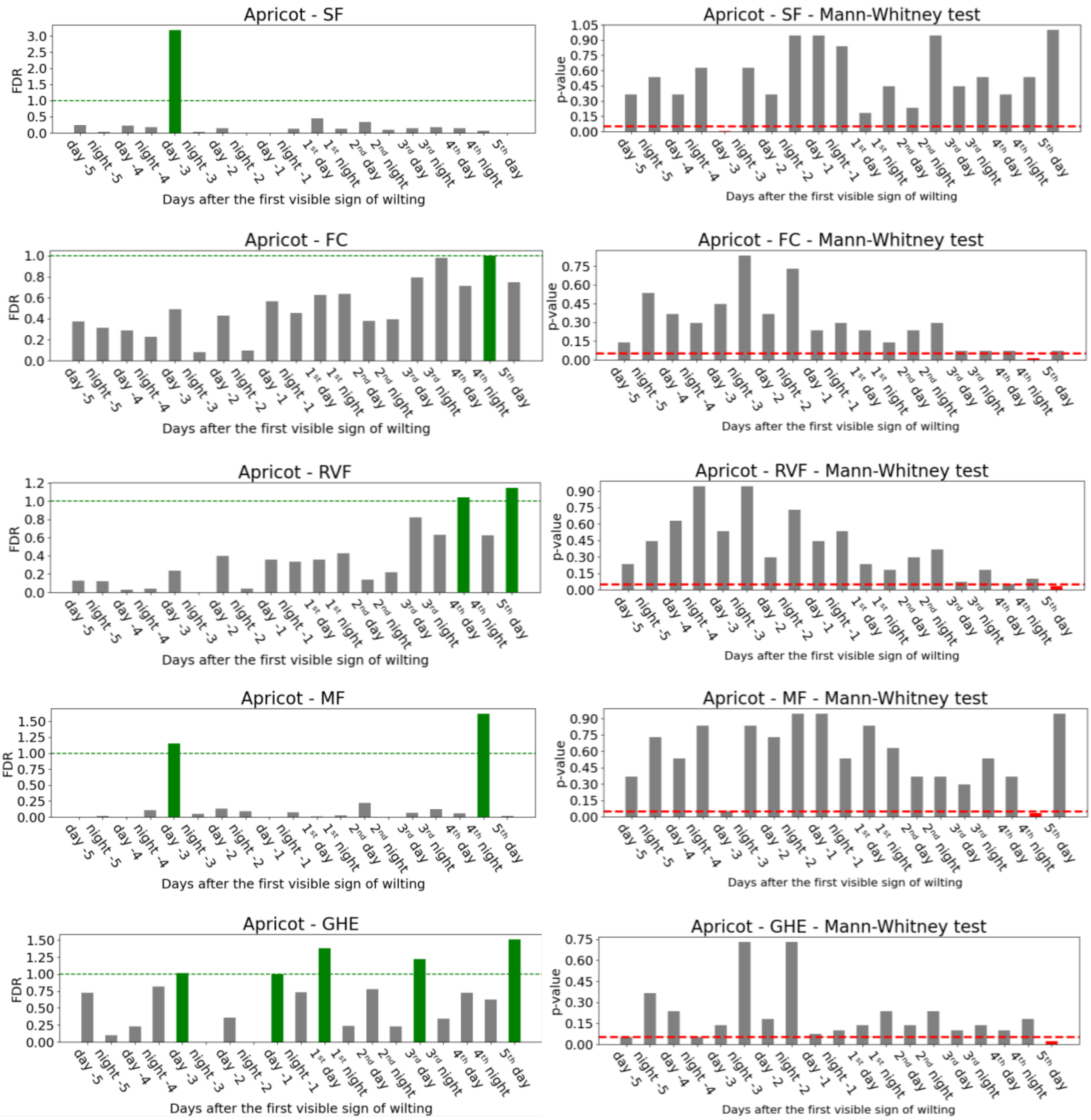

Apricot - SF
Apricot - SF - Mann-Whitney test
Apricot - FC
Apricot - FC - Mann-Whitney test
Apricot - RVF
Apricot - RVF - Mann-Whitney test
Apricot - MF
Apricot - MF - Mann-Whitney test
Apricot - GHE
Apricot - GHE - Mann-Whitney test
FDR
p-value
day -5, night -5, day -4, night -4, day -3, night -3, day -2, night -2, day -1, night -1, 1st day, 1st night, 2nd day, 2nd night, 3rd day, 3rd night, 4th day, 4th night, 5th day
Days after the first visible sign of wilting

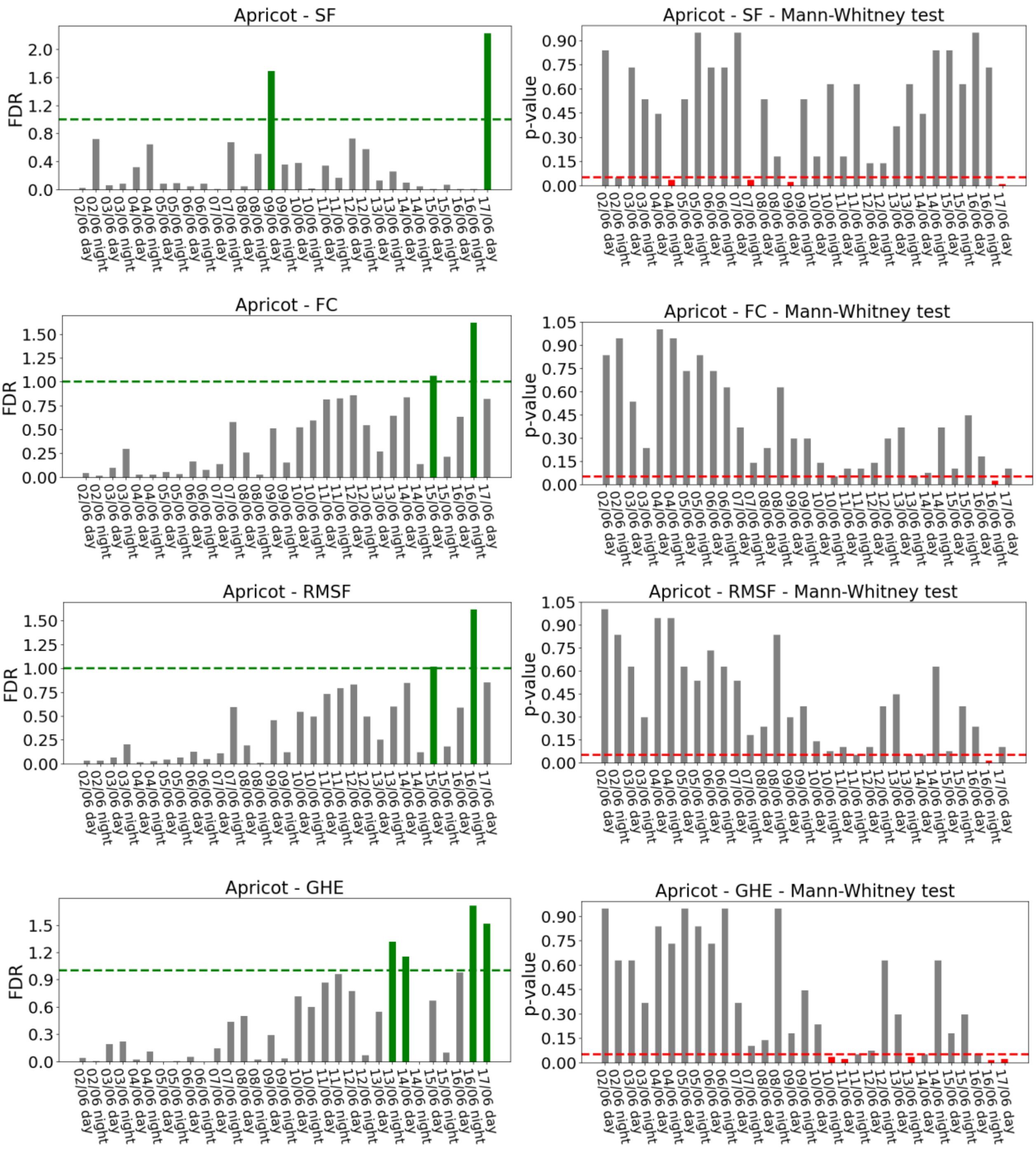


*Figure SM 3.1.3: FDR (on the left column) and p-value of the Mann-Whitney test (on the right column), showing the separation between control and stressed apricots, for all the features which exceeded the threshold of one for the FDR. Stressed plants are evaluated in the days around the first visible sign of wilting. In the figure, 1st day refers to the first day of the first signs.*

**Apricot – comparison based on plant status**

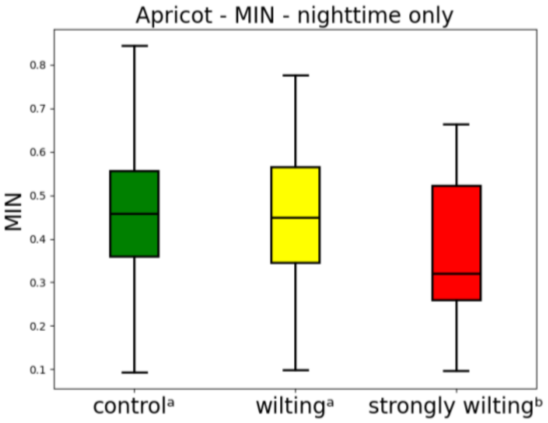


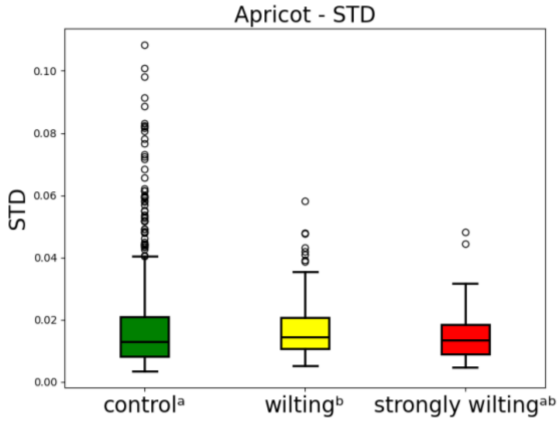


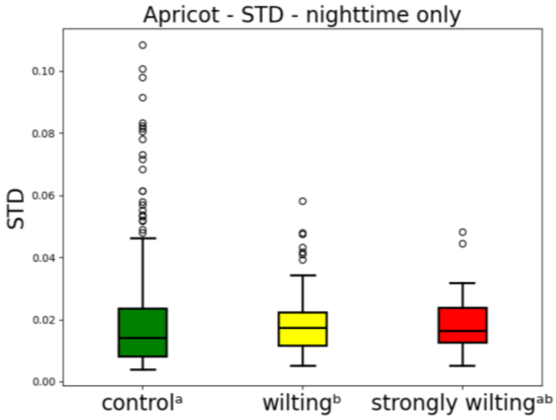


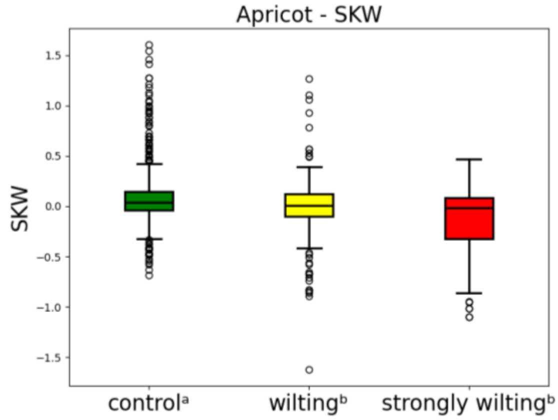


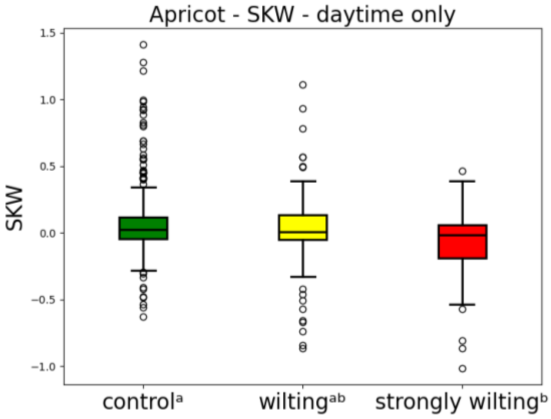


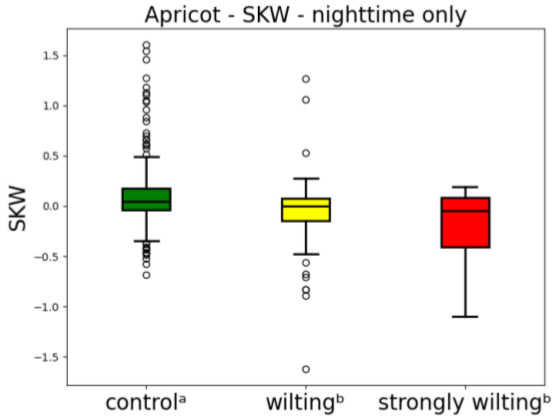


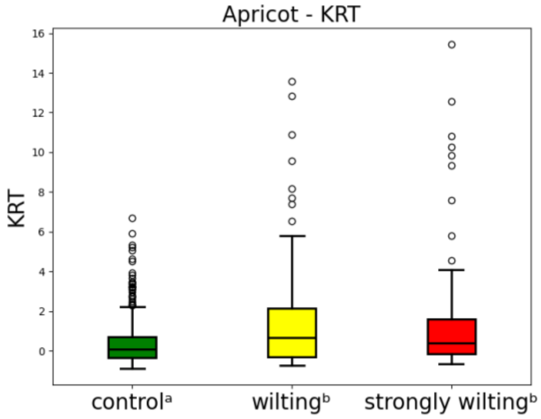


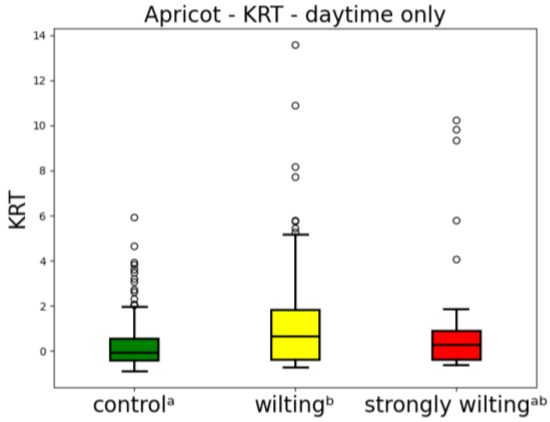

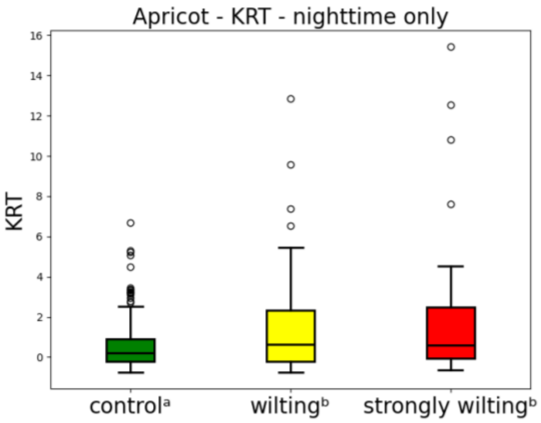
Apricot - KRT - nighttime only
KRT
control$^{a}$
wilting$^{b}$
strongly wilting$^{b}$

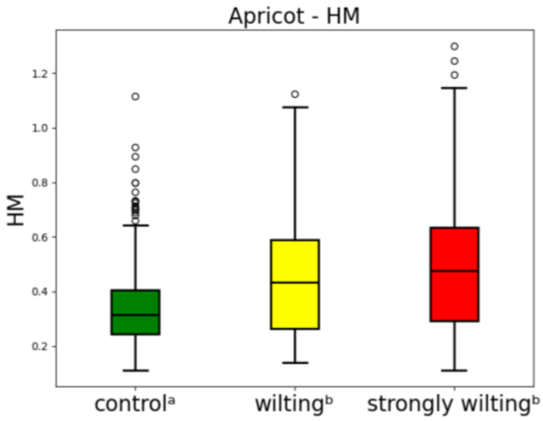
Apricot - HM
HM
control$^{a}$
wilting$^{b}$
strongly wilting$^{b}$

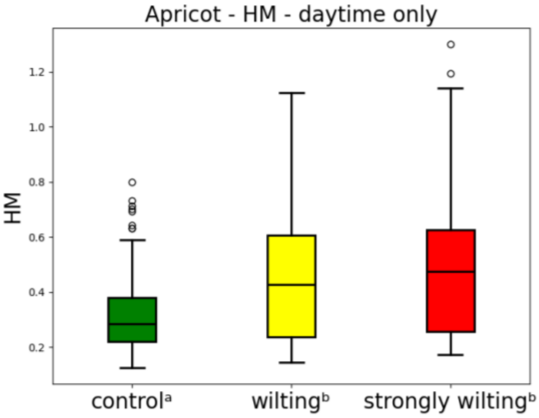
Apricot - HM - daytime only
HM
control$^{a}$
wilting$^{b}$
strongly wilting$^{b}$

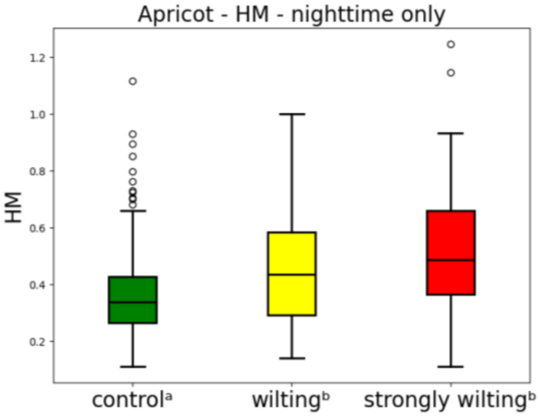
Apricot - HM - nighttime only
HM
control$^{a}$
wilting$^{b}$
strongly wilting$^{b}$

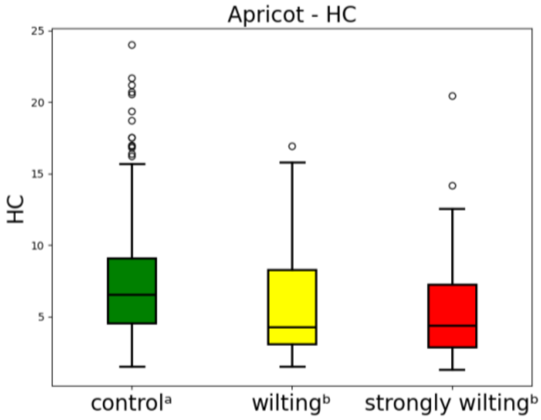
Apricot - HC
HC
control$^{a}$
wilting$^{b}$
strongly wilting$^{b}$

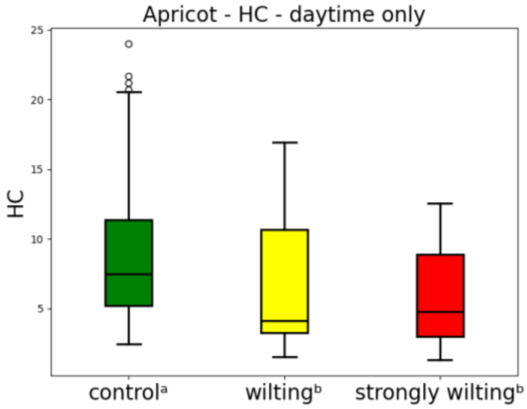
Apricot - HC - daytime only
HC
control$^{a}$
wilting$^{b}$
strongly wilting$^{b}$

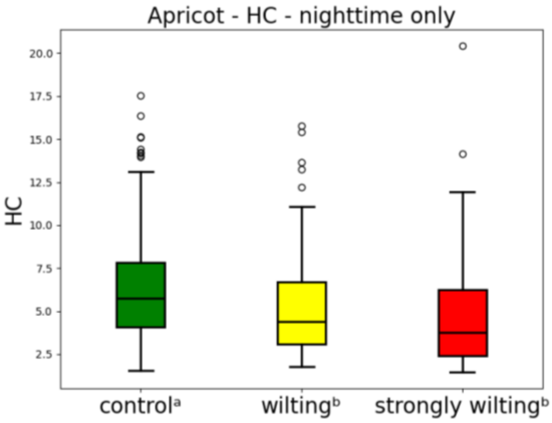
Apricot - HC - nighttime only
HC
control$^{a}$
wilting$^{b}$
strongly wilting$^{b}$

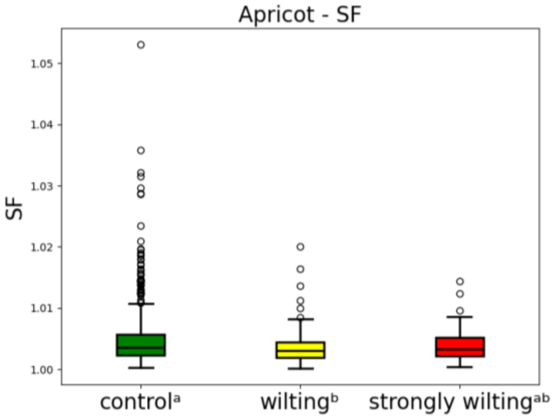
Apricot - SF
SF
control$^{a}$
wilting$^{b}$
strongly wilting$^{ab}$

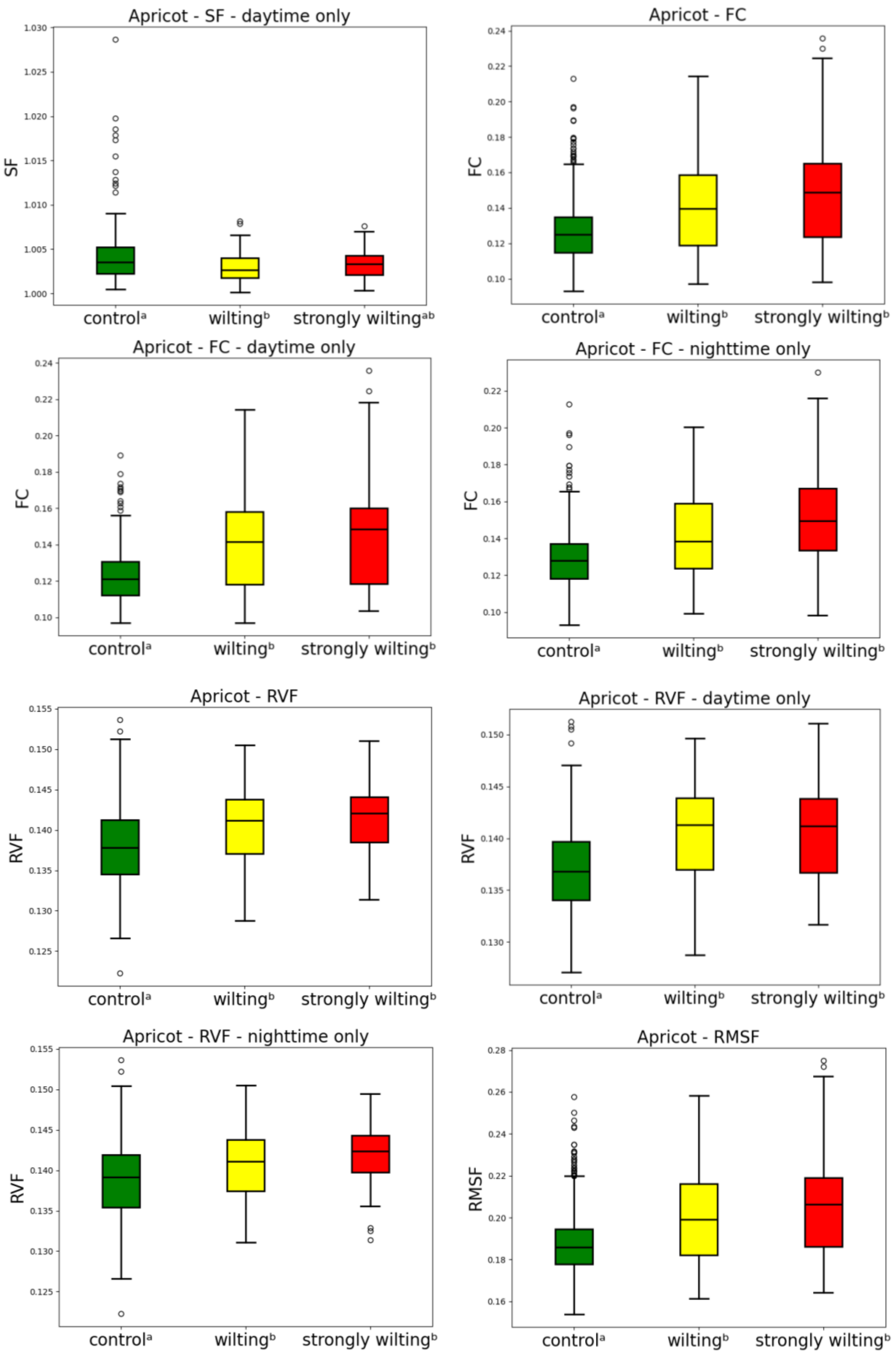

Apricot - SF - daytime only
SF
control[a] wilting[b] strongly wilting[ab]
Apricot - FC
FC
control[a] wilting[b] strongly wilting[b]
Apricot - FC - daytime only
FC
control[a] wilting[b] strongly wilting[b]
Apricot - FC - nighttime only
FC
control[a] wilting[b] strongly wilting[b]
Apricot - RVF
RVF
control[a] wilting[b] strongly wilting[b]
Apricot - RVF - daytime only
RVF
control[a] wilting[b] strongly wilting[b]
Apricot - RVF - nighttime only
RVF
control[a] wilting[b] strongly wilting[b]
Apricot - RMSF
RMSF
control[a] wilting[b] strongly wilting[b]

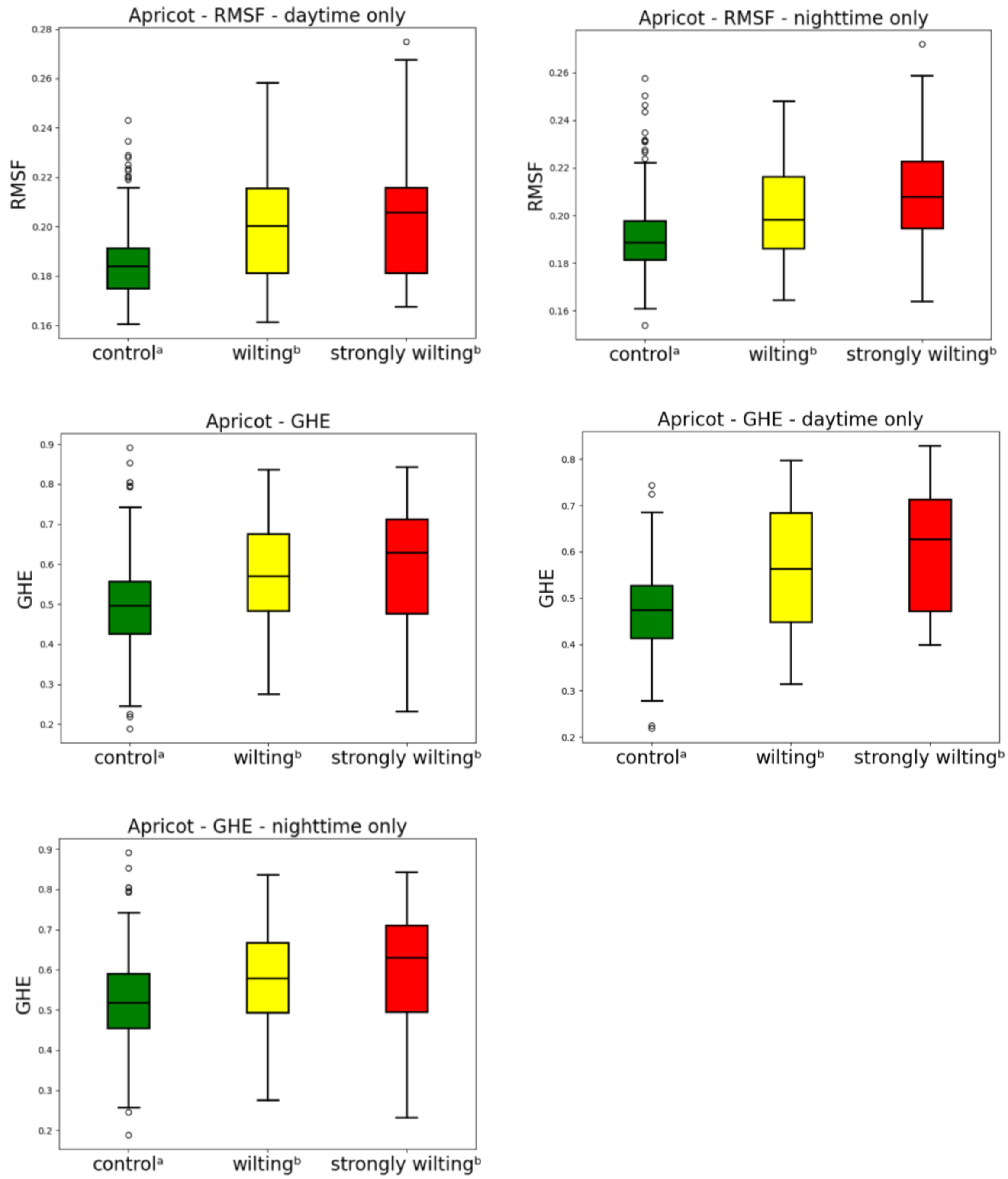


*Figure SM 3.1.4: Comparison of the value of the features computed for the control apricots and for the stressed apricots in two different conditions: wilting and strongly wilting. The superscript letters indicate statistically significant differences according to the Mann-Whitney test. The figure illustrates all features that showed at least one significant difference between two different conditions.*

### Apricot – interquartile range of the features around the first day of visible wilting

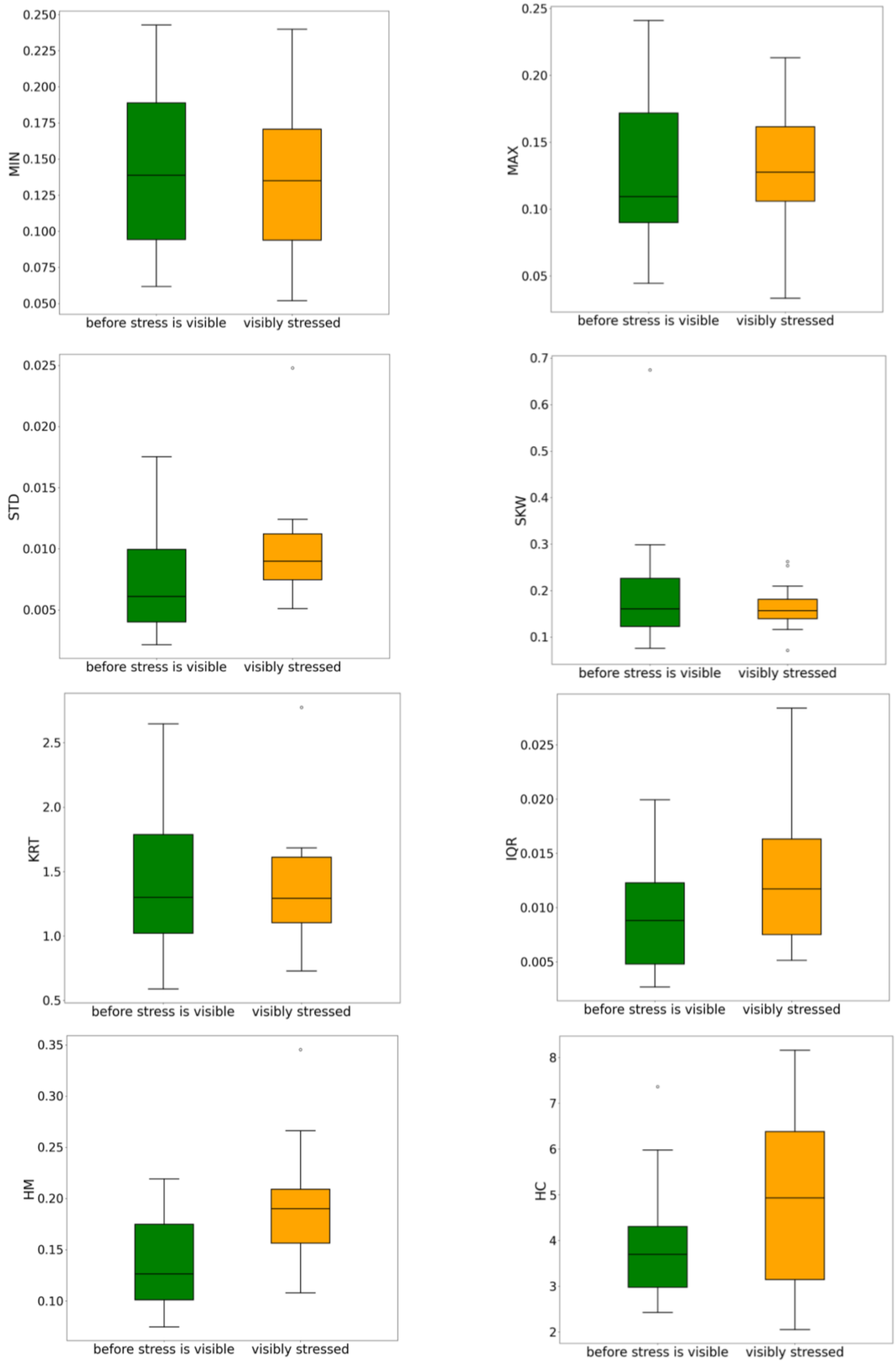

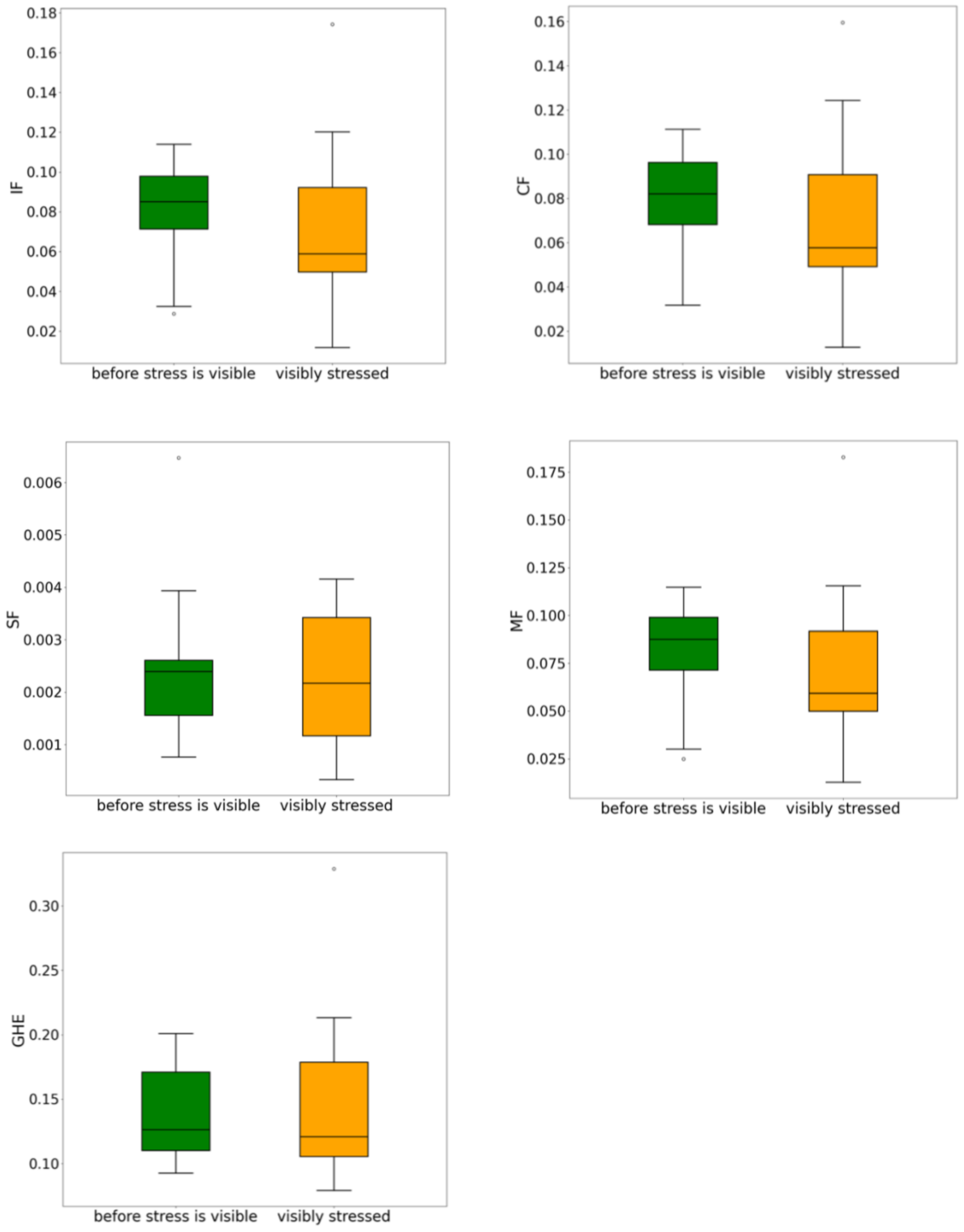


*Figure SM 3.1.5: Interquartile range of the features in the stressed apricots, in the six days before the first visible signs of wilting and on the following six days. All the features illustrated here did not show a significant difference between the two periods. Figure 5 shows the three features for which the difference was statistically significant.*

**SM 3.2 Frequency domain analysis**

**Tomato – Frequency domain features**

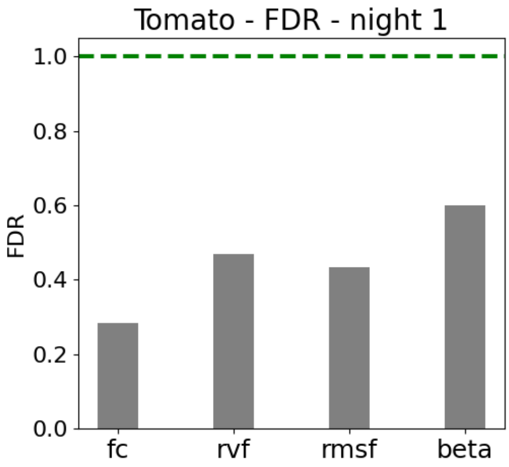


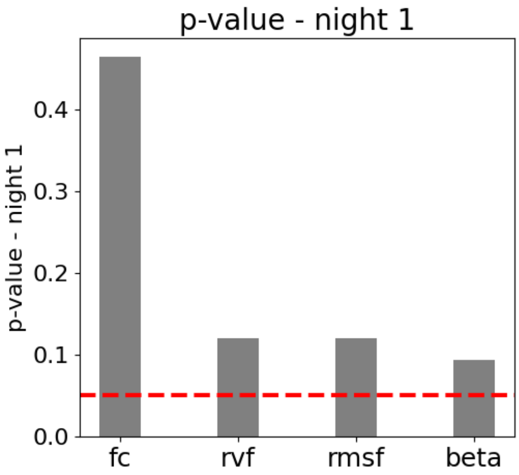


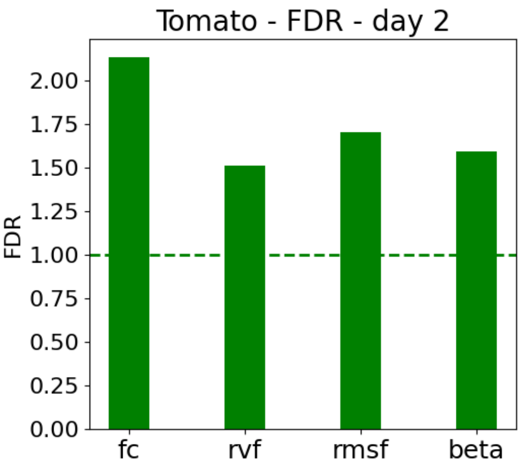


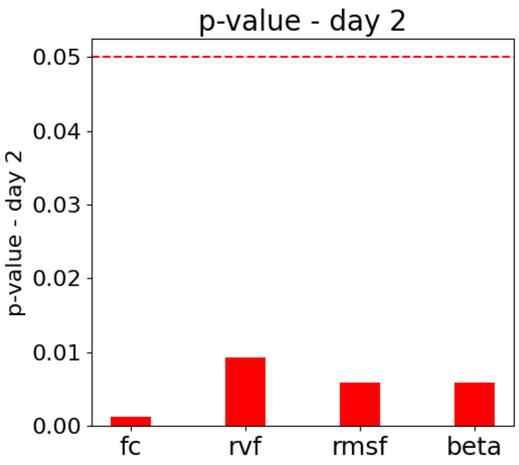


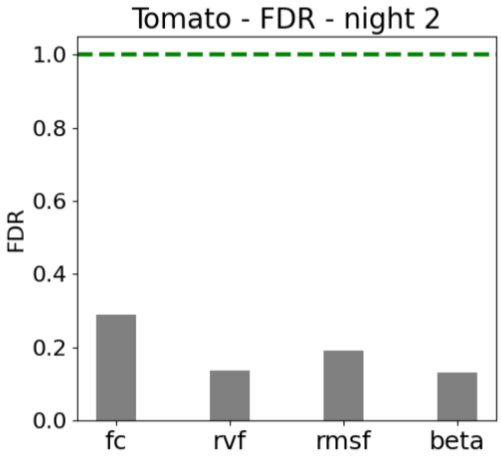


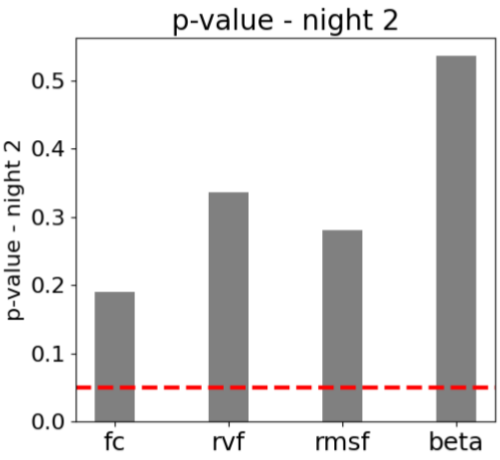

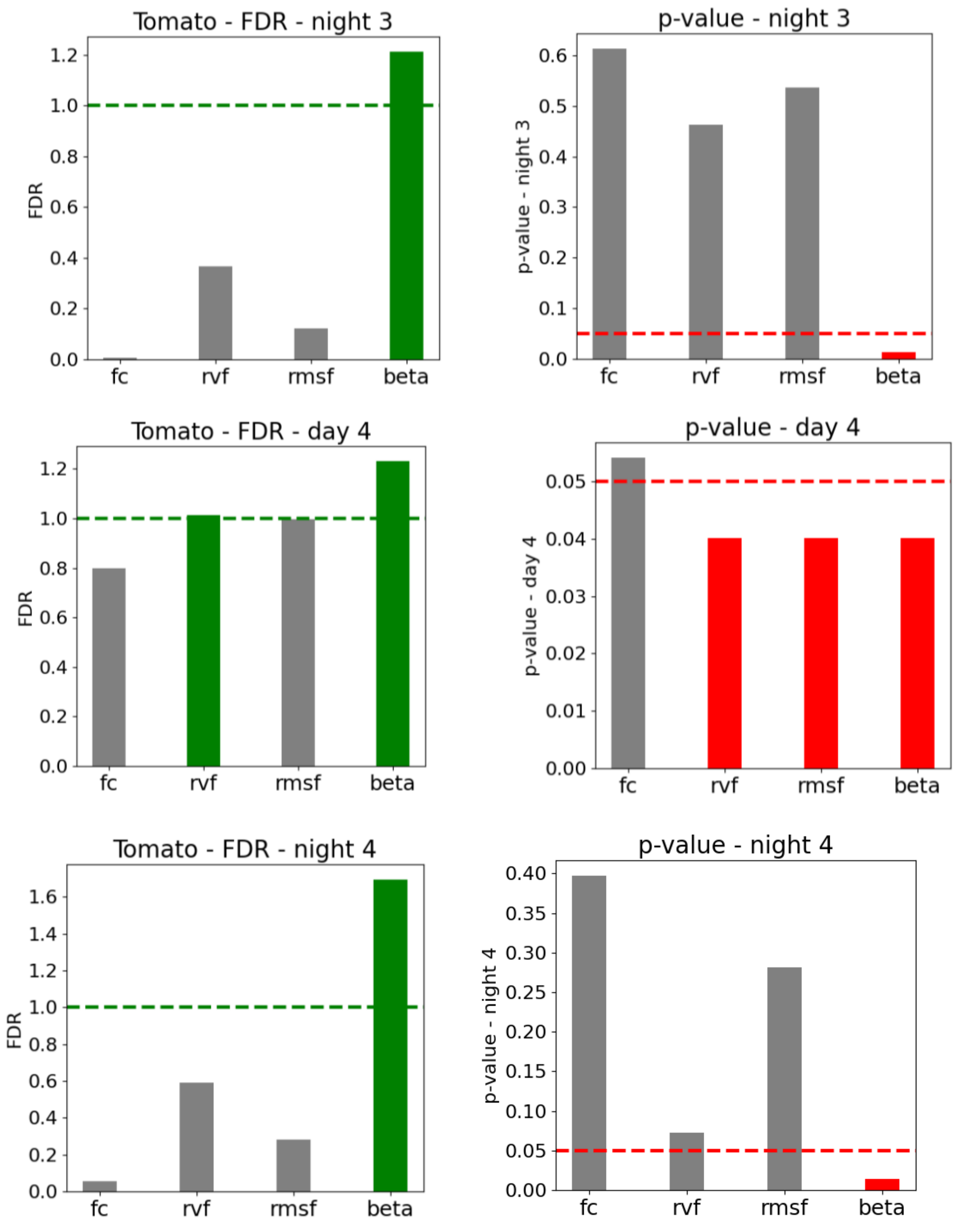

Tomato - FDR - night 3
FDR
1.2
1.0
0.8
0.6
0.4
0.2
0.0
fc
rvf
rmsf
beta
p-value - night 3
p-value - night 3
0.6
0.5
0.4
0.3
0.2
0.1
0.0
fc
rvf
rmsf
beta
Tomato - FDR - day 4
FDR
1.2
1.0
0.8
0.6
0.4
0.2
0.0
fc
rvf
rmsf
beta
p-value - day 4
p-value - day 4
0.05
0.04
0.03
0.02
0.01
0.00
fc
rvf
rmsf
beta
Tomato - FDR - night 4
FDR
1.6
1.4
1.2
1.0
0.8
0.6
0.4
0.2
0.0
fc
rvf
rmsf
beta
p-value - night 4
p-value - night 4
0.40
0.35
0.30
0.25
0.20
0.15
0.10
0.05
0.00
fc
rvf
rmsf
beta

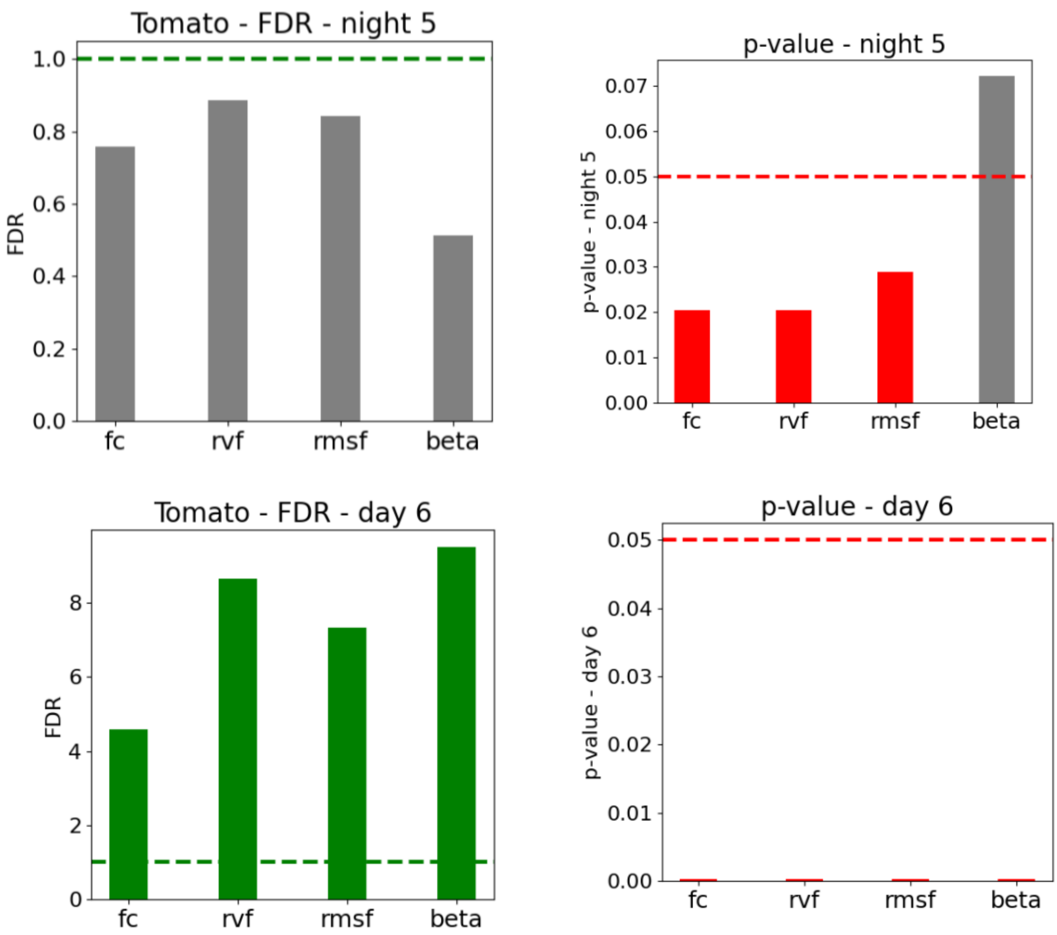


*Figure SM 3.2.1: FDR (on the left column) and p-value of the Mann-Whitney test (on the right column), showing the separation between control and stressed tomatoes, for the frequency-domain features computed on daylight and nighttime periods.*

**Apricot – Frequency domain features**

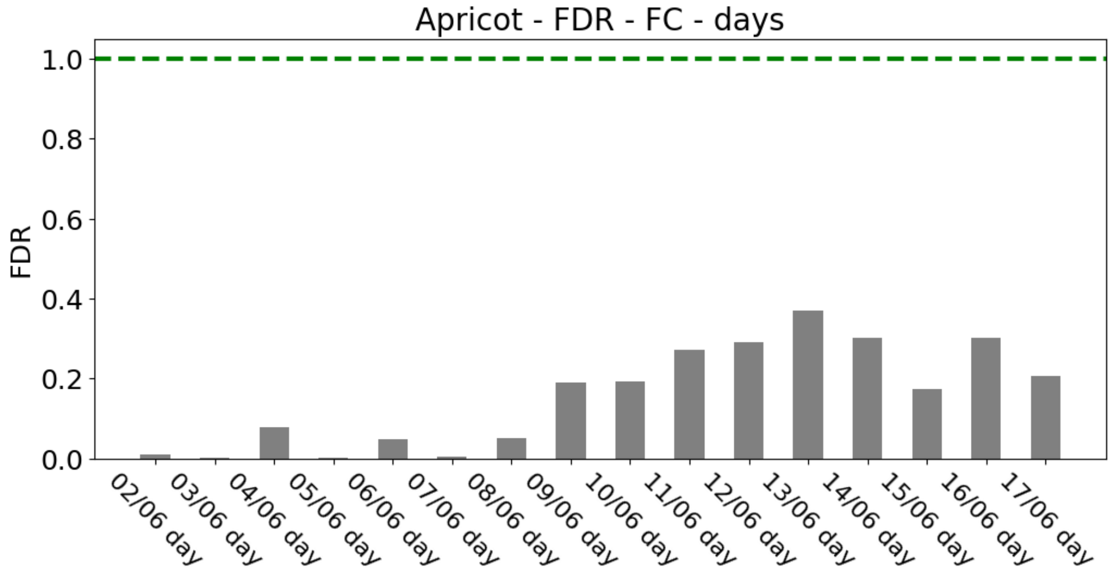


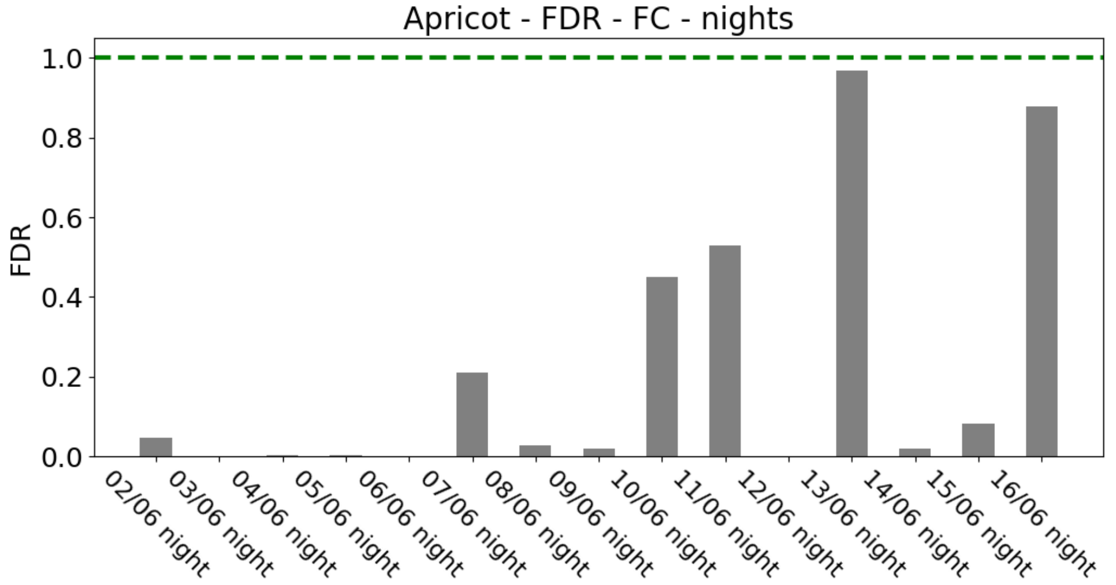


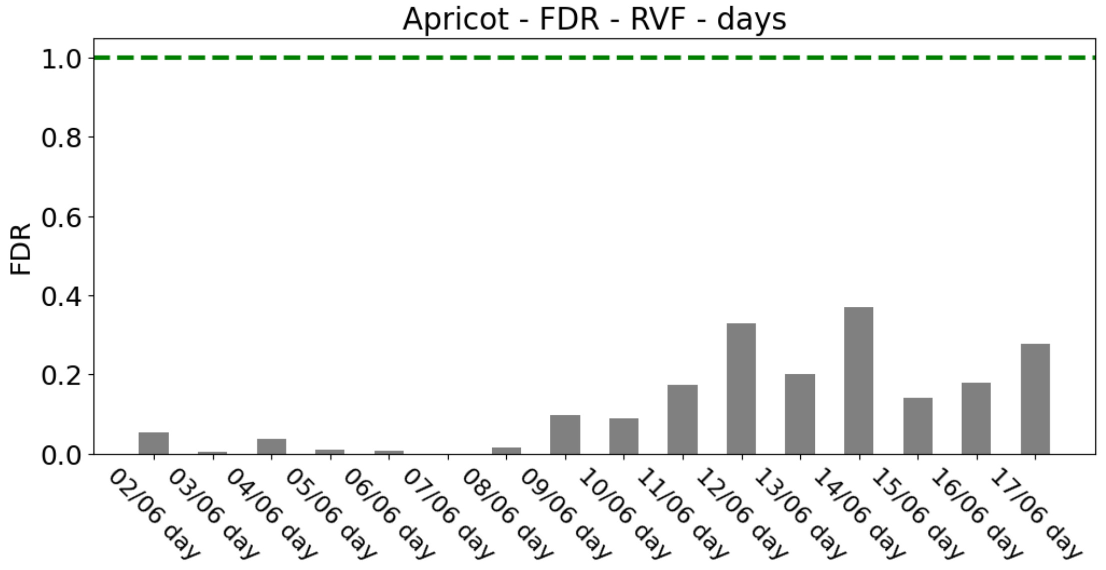

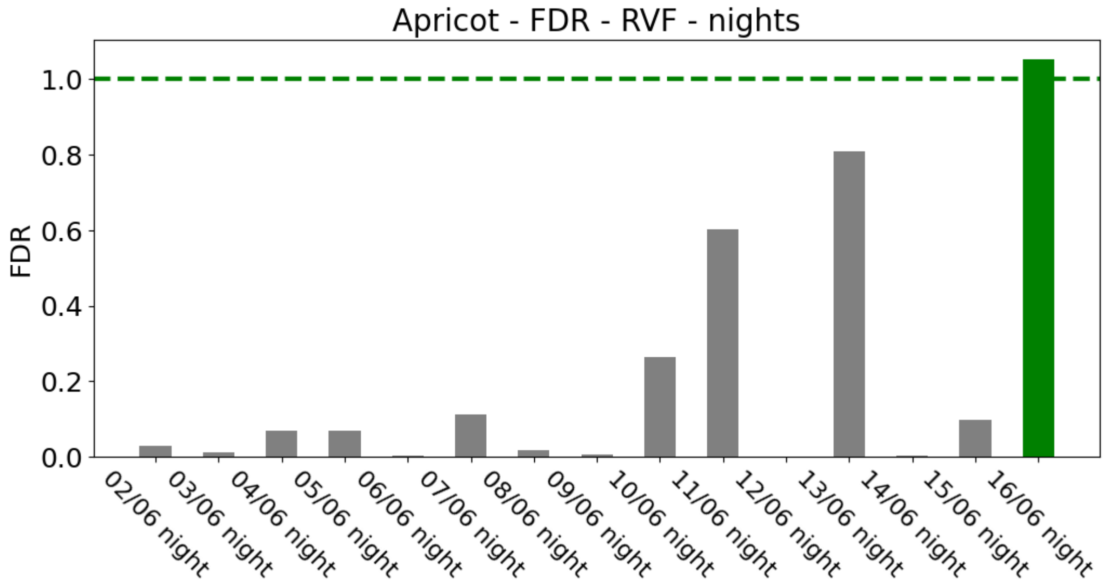
Apricot - FDR - RVF - nights
FDR
1.0
0.8
0.6
0.4
0.2
0.0
02/06 night
03/06 night
04/06 night
05/06 night
06/06 night
07/06 night
08/06 night
09/06 night
10/06 night
11/06 night
12/06 night
13/06 night
14/06 night
15/06 night
16/06 night

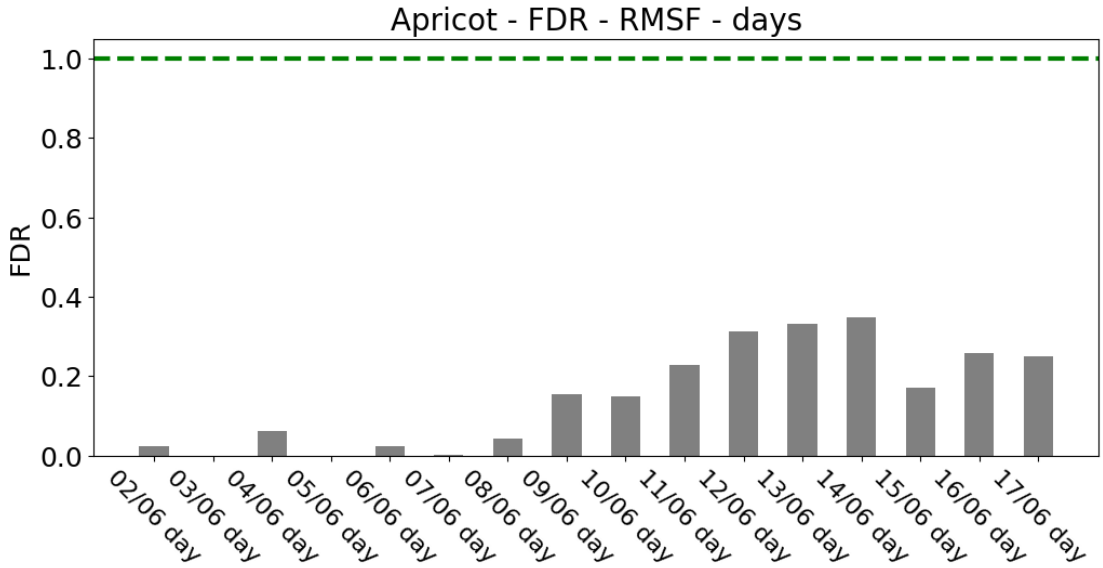
Apricot - FDR - RMSF - days
FDR
1.0
0.8
0.6
0.4
0.2
0.0
02/06 day
03/06 day
04/06 day
05/06 day
06/06 day
07/06 day
08/06 day
09/06 day
10/06 day
11/06 day
12/06 day
13/06 day
14/06 day
15/06 day
16/06 day
17/06 day

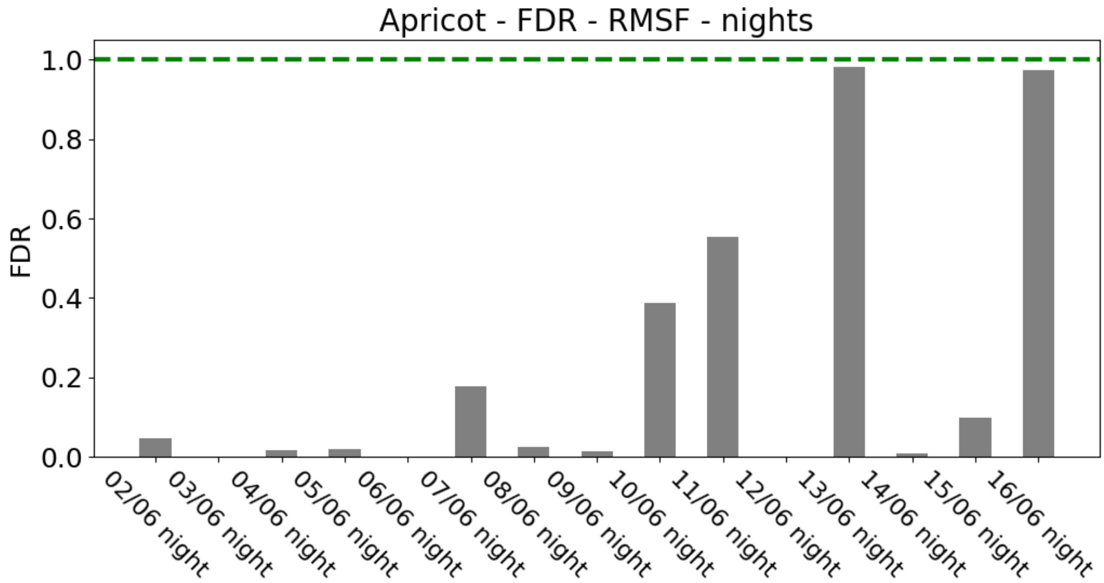
Apricot - FDR - RMSF - nights
FDR
1.0
0.8
0.6
0.4
0.2
0.0
02/06 night
03/06 night
04/06 night
05/06 night
06/06 night
07/06 night
08/06 night
09/06 night
10/06 night
11/06 night
12/06 night
13/06 night
14/06 night
15/06 night
16/06 night

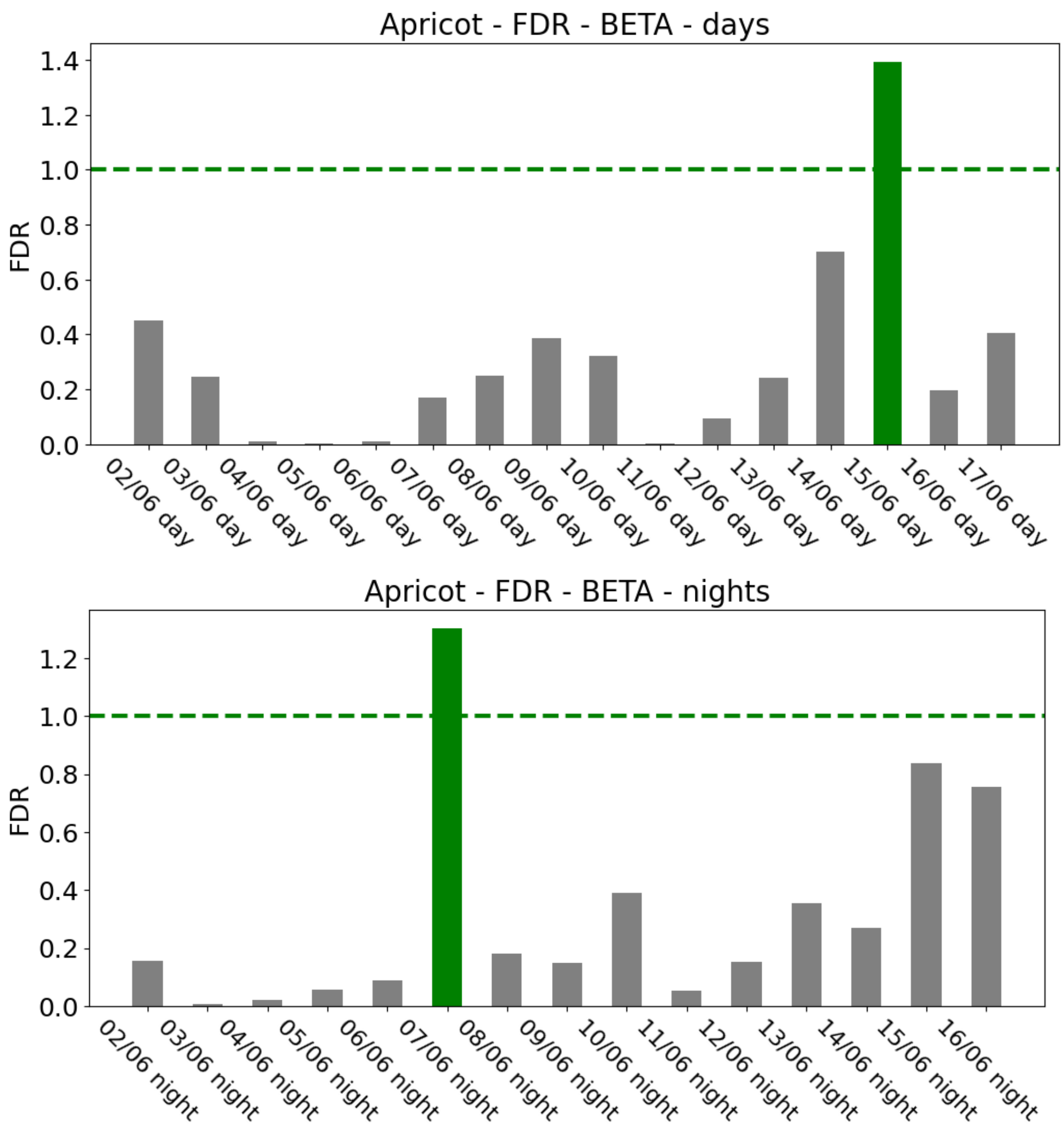


*Figure SM 3.2.2: FDR (on the left column) and p-value of the Mann-Whitney test (on the right column), showing the separation between control and stressed apricots, for the frequency-domain features computed on daylight and nighttime periods.*

**Tomato – Power Spectral Density Graphs**

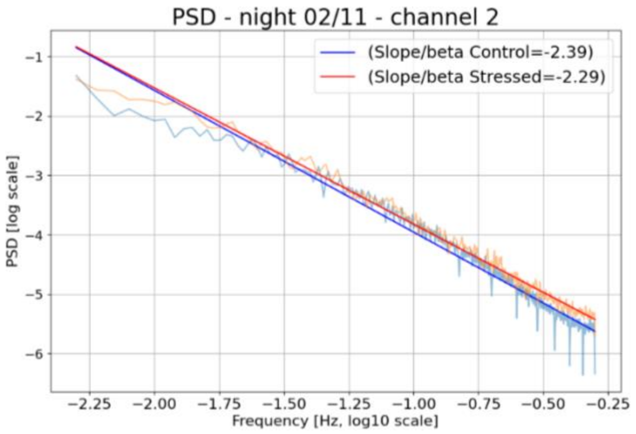


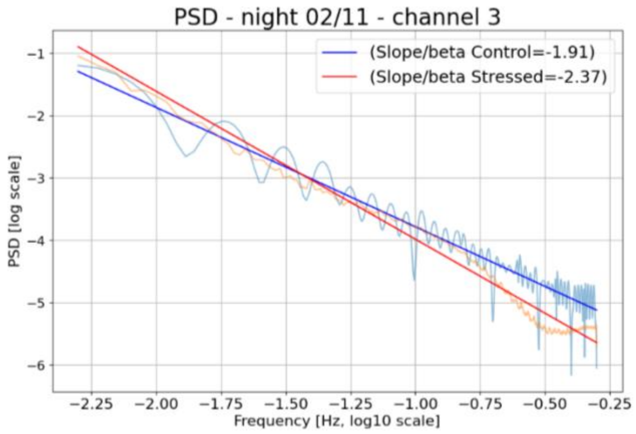


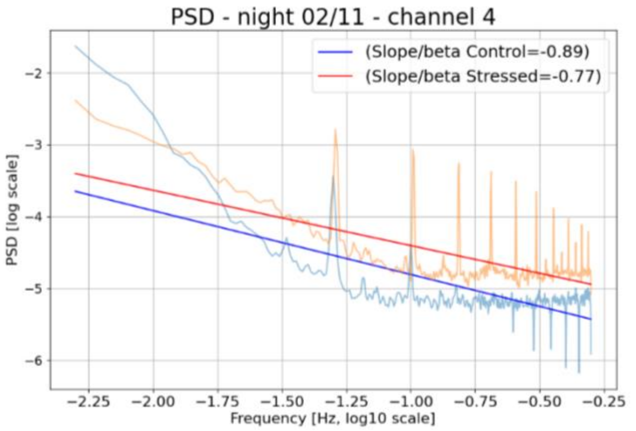


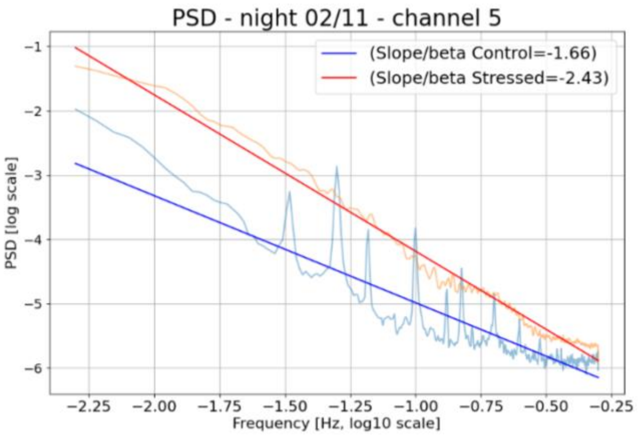


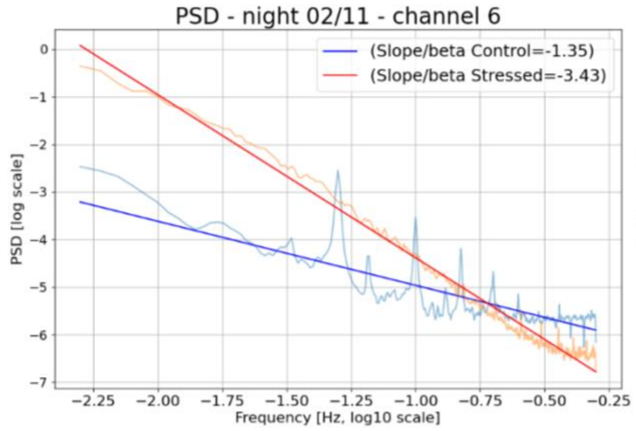


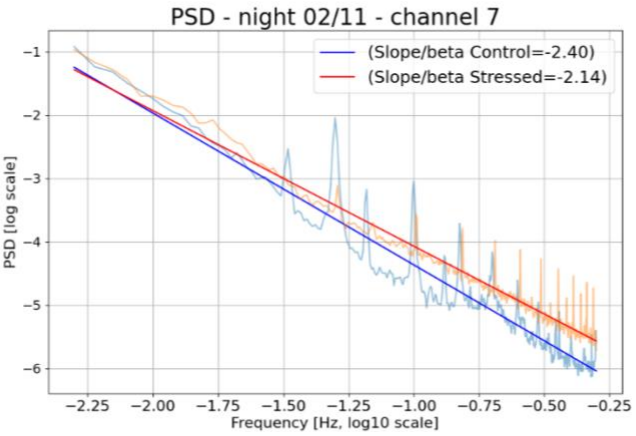


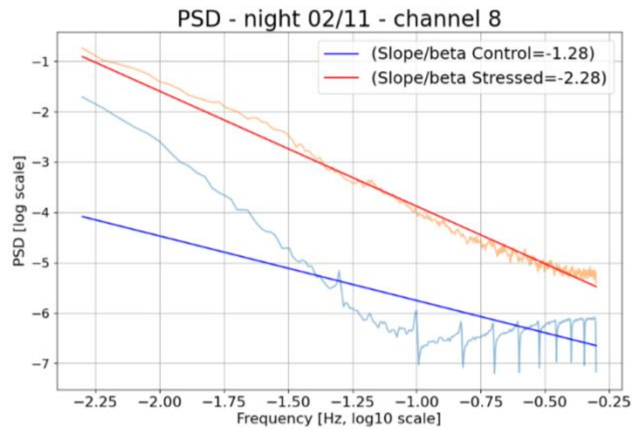

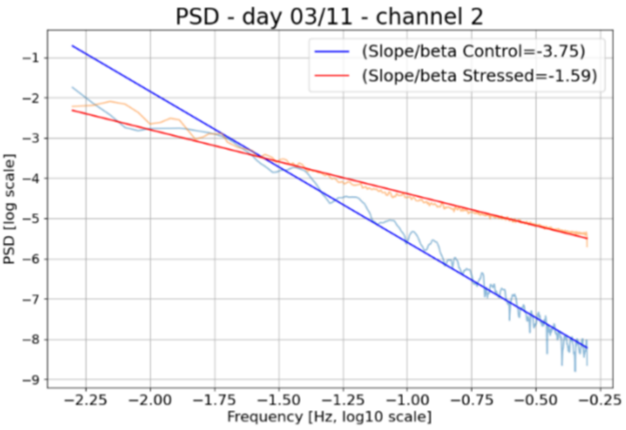
PSD - day 03/11 - channel 2
(Slope/beta Control=-3.75)
(Slope/beta Stressed=-1.59)
PSD [log scale]
Frequency [Hz, log10 scale]

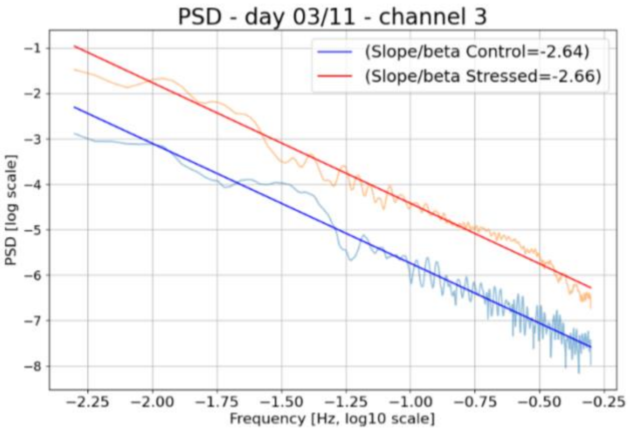
PSD - day 03/11 - channel 3
(Slope/beta Control=-2.64)
(Slope/beta Stressed=-2.66)
PSD [log scale]
Frequency [Hz, log10 scale]

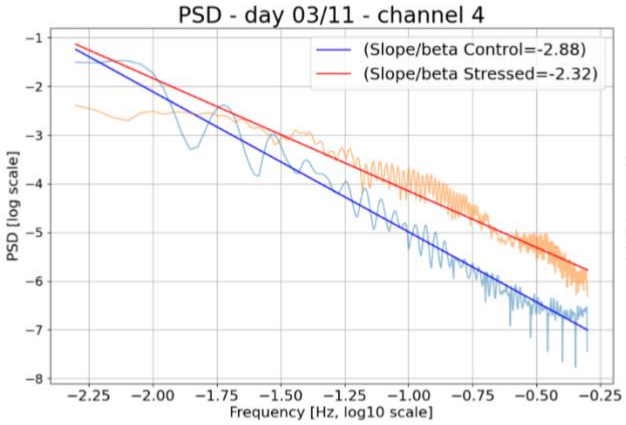
PSD - day 03/11 - channel 4
(Slope/beta Control=-2.88)
(Slope/beta Stressed=-2.32)
PSD [log scale]
Frequency [Hz, log10 scale]

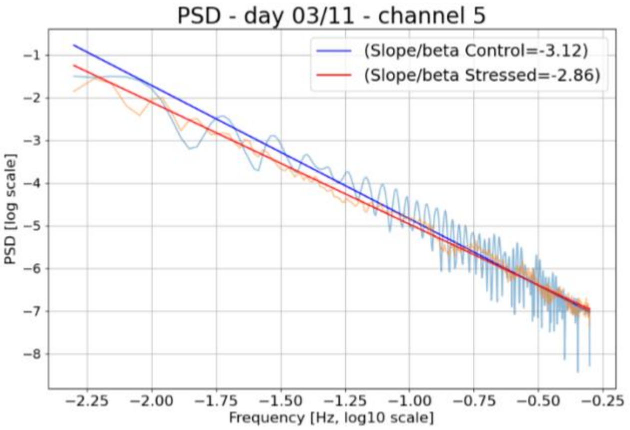
PSD - day 03/11 - channel 5
(Slope/beta Control=-3.12)
(Slope/beta Stressed=-2.86)
PSD [log scale]
Frequency [Hz, log10 scale]

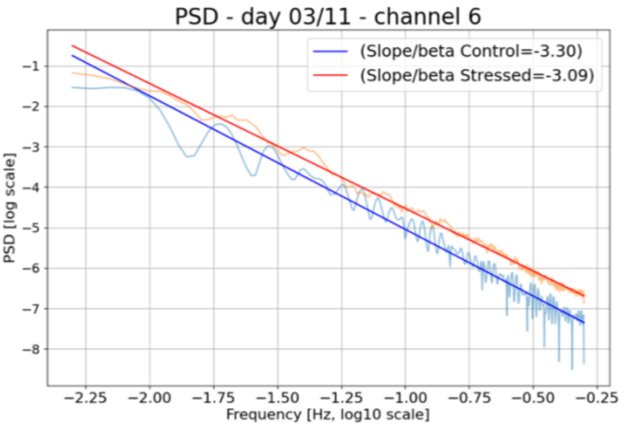
PSD - day 03/11 - channel 6
(Slope/beta Control=-3.30)
(Slope/beta Stressed=-3.09)
PSD [log scale]
Frequency [Hz, log10 scale]

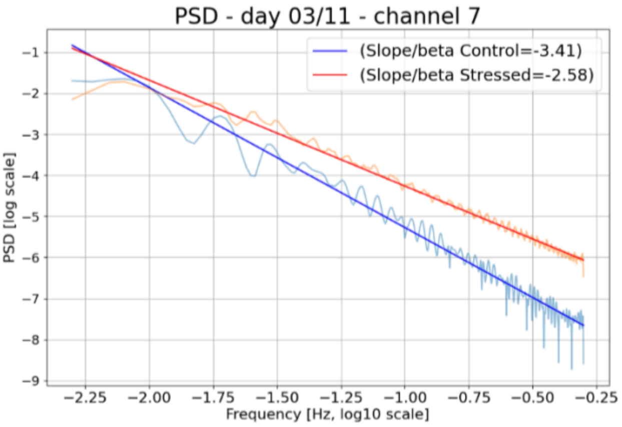
PSD - day 03/11 - channel 7
(Slope/beta Control=-3.41)
(Slope/beta Stressed=-2.58)
PSD [log scale]
Frequency [Hz, log10 scale]

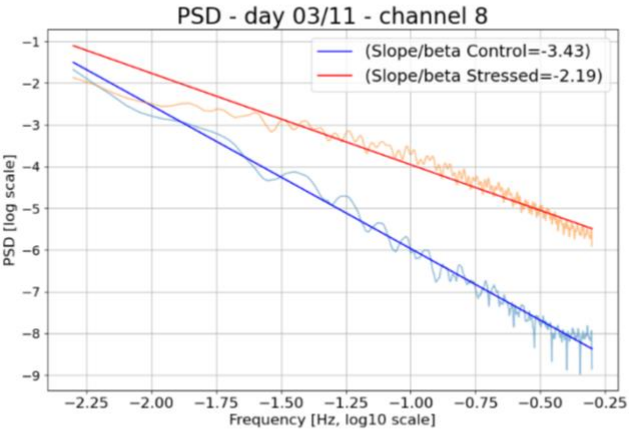
PSD - day 03/11 - channel 8
(Slope/beta Control=-3.43)
(Slope/beta Stressed=-2.19)
PSD [log scale]
Frequency [Hz, log10 scale]

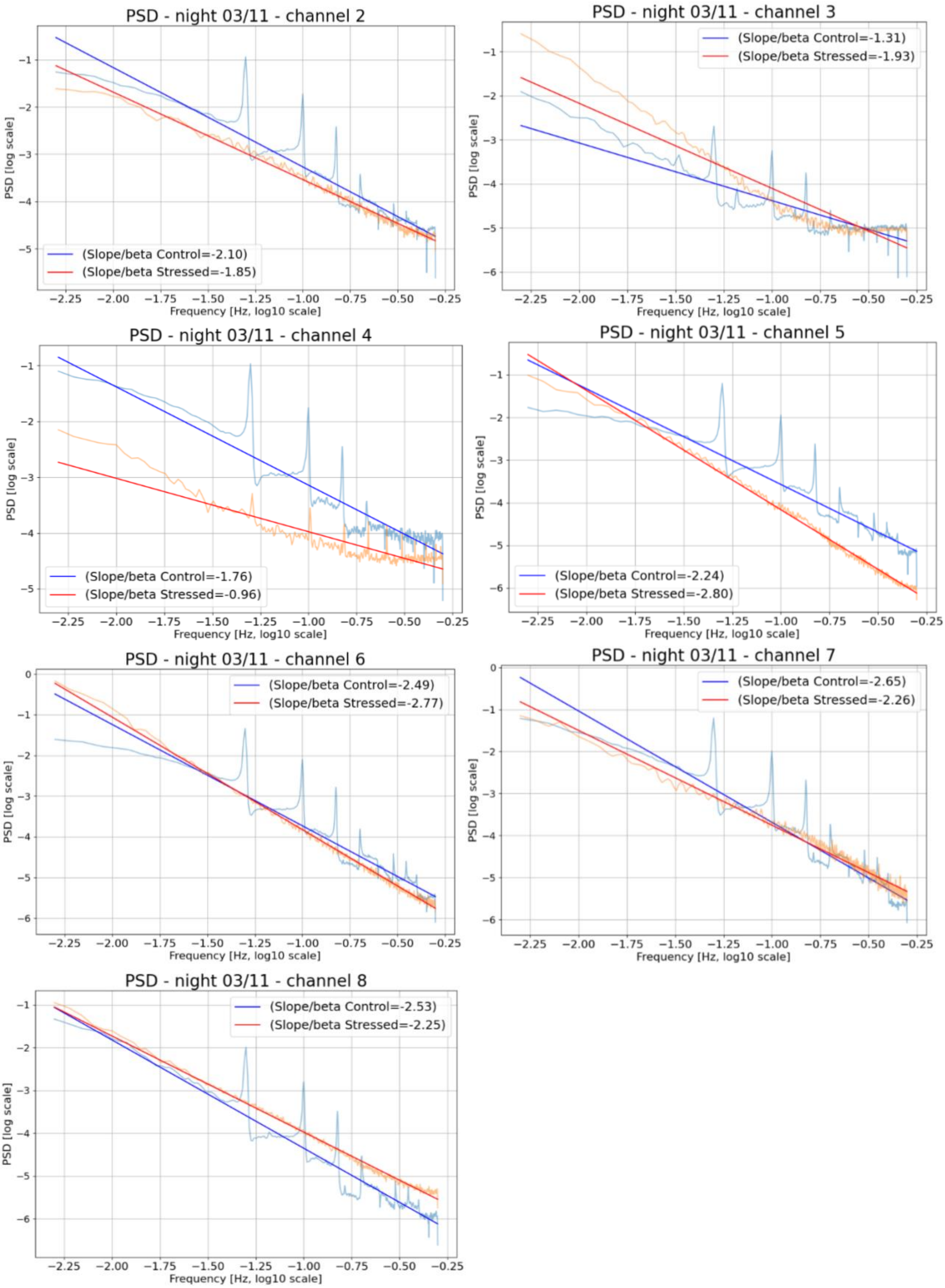
PSD - night 03/11 - channel 2
(Slope/beta Control=-2.10)
(Slope/beta Stressed=-1.85)
PSD - night 03/11 - channel 3
(Slope/beta Control=-1.31)
(Slope/beta Stressed=-1.93)
PSD - night 03/11 - channel 4
(Slope/beta Control=-1.76)
(Slope/beta Stressed=-0.96)
PSD - night 03/11 - channel 5
(Slope/beta Control=-2.24)
(Slope/beta Stressed=-2.80)
PSD - night 03/11 - channel 6
(Slope/beta Control=-2.49)
(Slope/beta Stressed=-2.77)
PSD - night 03/11 - channel 7
(Slope/beta Control=-2.65)
(Slope/beta Stressed=-2.26)
PSD - night 03/11 - channel 8
(Slope/beta Control=-2.53)
(Slope/beta Stressed=-2.25)
PSD [log scale]
Frequency [Hz, log10 scale]

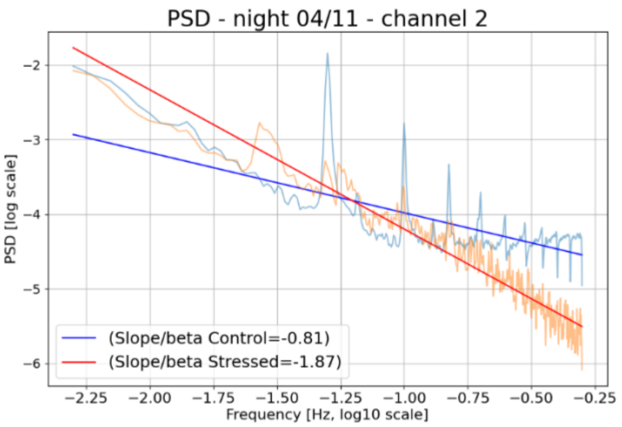
PSD - night 04/11 - channel 2
(Slope/beta Control=-0.81)
(Slope/beta Stressed=-1.87)
PSD [log scale]
Frequency [Hz, log10 scale]

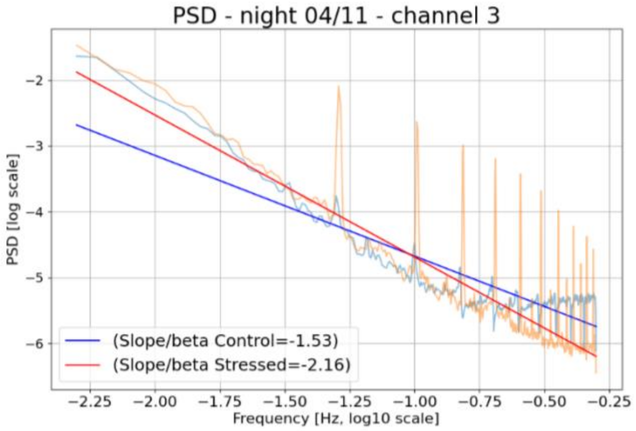
PSD - night 04/11 - channel 3
(Slope/beta Control=-1.53)
(Slope/beta Stressed=-2.16)
PSD [log scale]
Frequency [Hz, log10 scale]

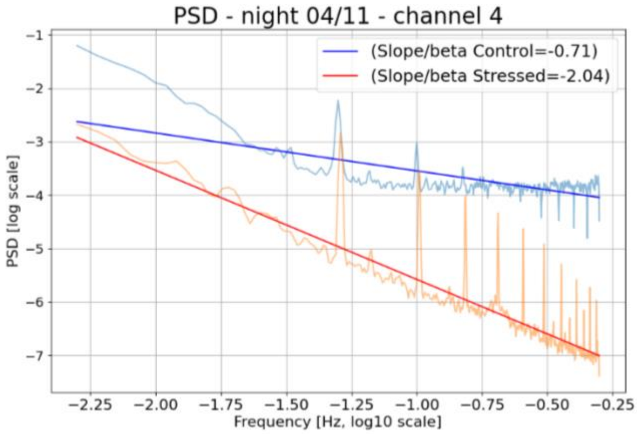
PSD - night 04/11 - channel 4
(Slope/beta Control=-0.71)
(Slope/beta Stressed=-2.04)
PSD [log scale]
Frequency [Hz, log10 scale]

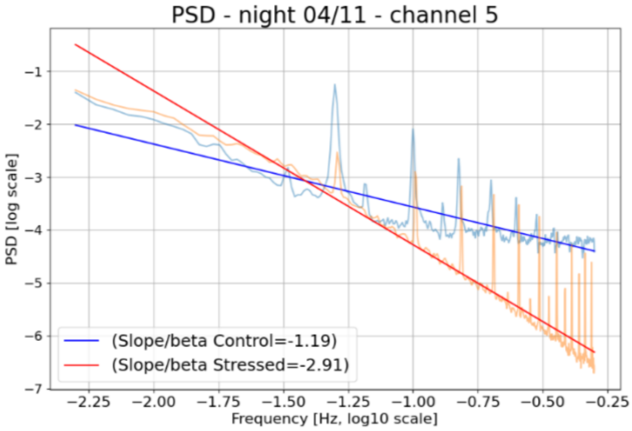
PSD - night 04/11 - channel 5
(Slope/beta Control=-1.19)
(Slope/beta Stressed=-2.91)
PSD [log scale]
Frequency [Hz, log10 scale]

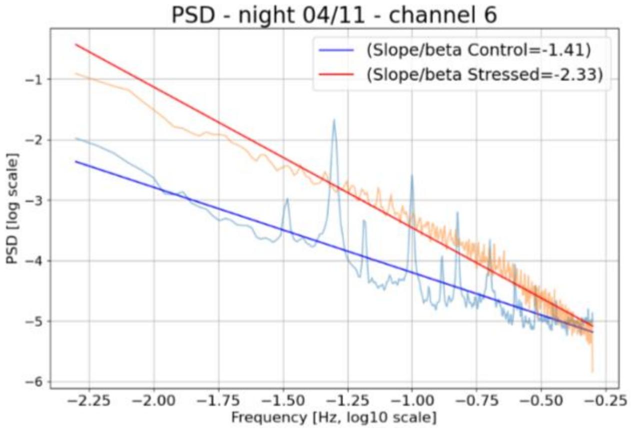
PSD - night 04/11 - channel 6
(Slope/beta Control=-1.41)
(Slope/beta Stressed=-2.33)
PSD [log scale]
Frequency [Hz, log10 scale]

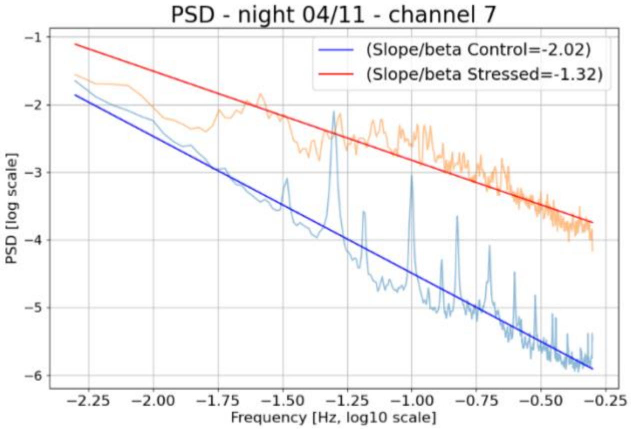
PSD - night 04/11 - channel 7
(Slope/beta Control=-2.02)
(Slope/beta Stressed=-1.32)
PSD [log scale]
Frequency [Hz, log10 scale]

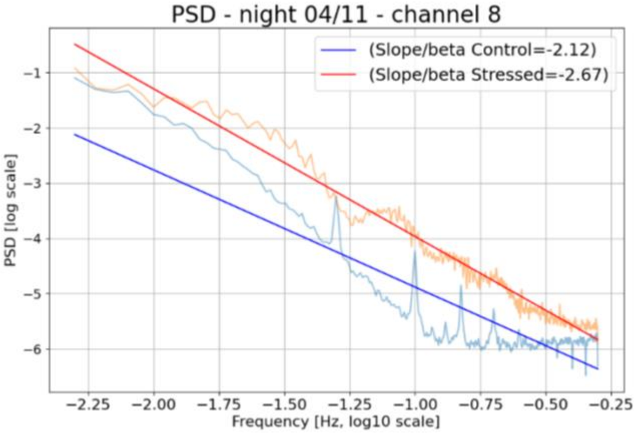
PSD - night 04/11 - channel 8
(Slope/beta Control=-2.12)
(Slope/beta Stressed=-2.67)
PSD [log scale]
Frequency [Hz, log10 scale]

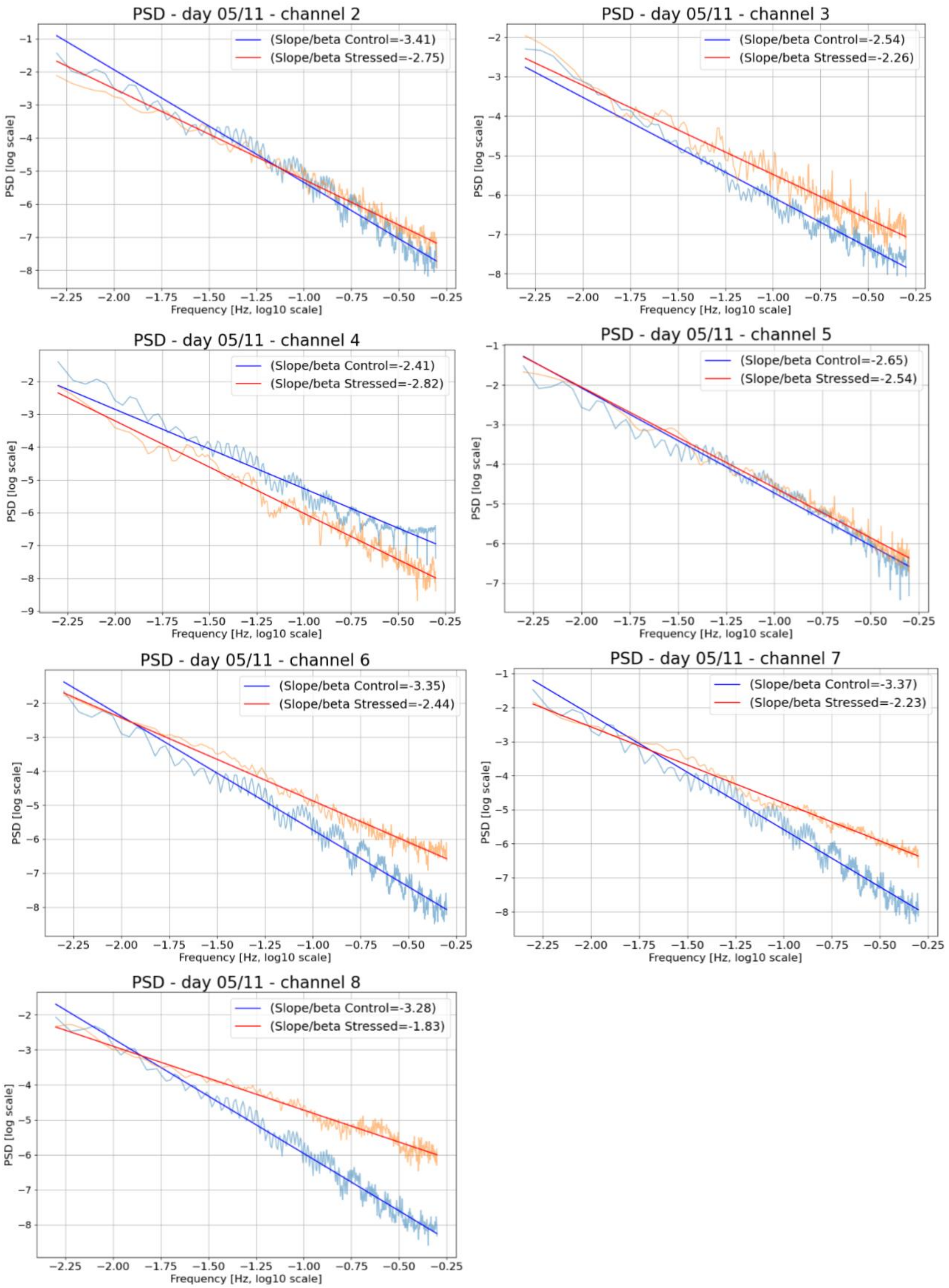
PSD - day 05/11 - channel 2
(Slope/beta Control=-3.41)
(Slope/beta Stressed=-2.75)
PSD [log scale]
Frequency [Hz, log10 scale]
PSD - day 05/11 - channel 3
(Slope/beta Control=-2.54)
(Slope/beta Stressed=-2.26)
PSD [log scale]
Frequency [Hz, log10 scale]
PSD - day 05/11 - channel 4
(Slope/beta Control=-2.41)
(Slope/beta Stressed=-2.82)
PSD [log scale]
Frequency [Hz, log10 scale]
PSD - day 05/11 - channel 5
(Slope/beta Control=-2.65)
(Slope/beta Stressed=-2.54)
PSD [log scale]
Frequency [Hz, log10 scale]
PSD - day 05/11 - channel 6
(Slope/beta Control=-3.35)
(Slope/beta Stressed=-2.44)
PSD [log scale]
Frequency [Hz, log10 scale]
PSD - day 05/11 - channel 7
(Slope/beta Control=-3.37)
(Slope/beta Stressed=-2.23)
PSD [log scale]
Frequency [Hz, log10 scale]
PSD - day 05/11 - channel 8
(Slope/beta Control=-3.28)
(Slope/beta Stressed=-1.83)
PSD [log scale]
Frequency [Hz, log10 scale]

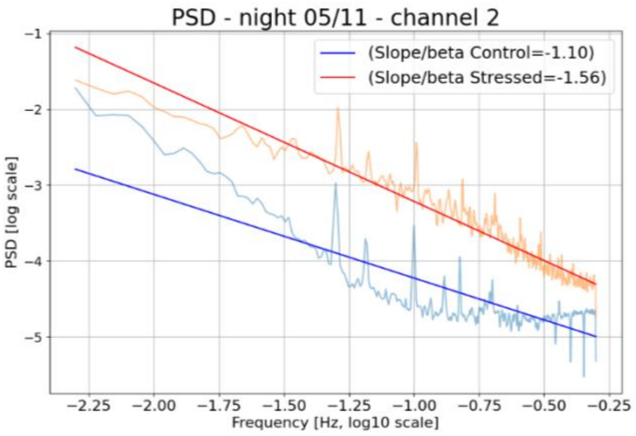
PSD - night 05/11 - channel 2
(Slope/beta Control=-1.10)
(Slope/beta Stressed=-1.56)
PSD [log scale]
Frequency [Hz, log10 scale]

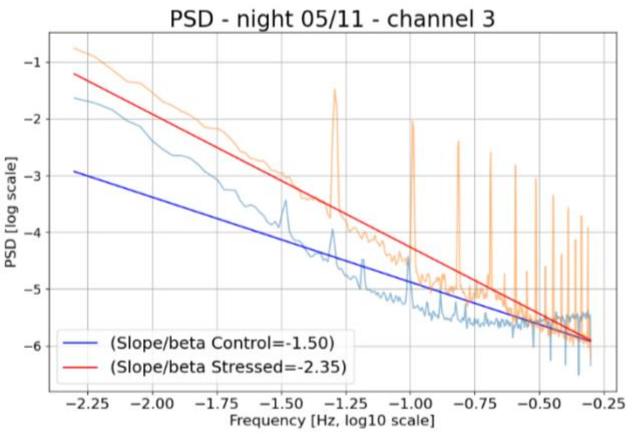
PSD - night 05/11 - channel 3
(Slope/beta Control=-1.50)
(Slope/beta Stressed=-2.35)
PSD [log scale]
Frequency [Hz, log10 scale]

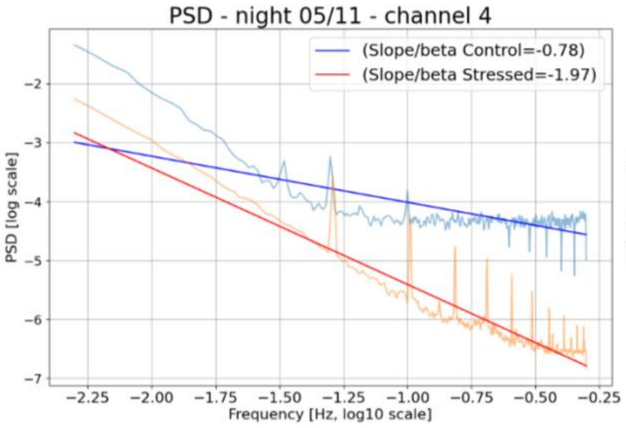
PSD - night 05/11 - channel 4
(Slope/beta Control=-0.78)
(Slope/beta Stressed=-1.97)
PSD [log scale]
Frequency [Hz, log10 scale]

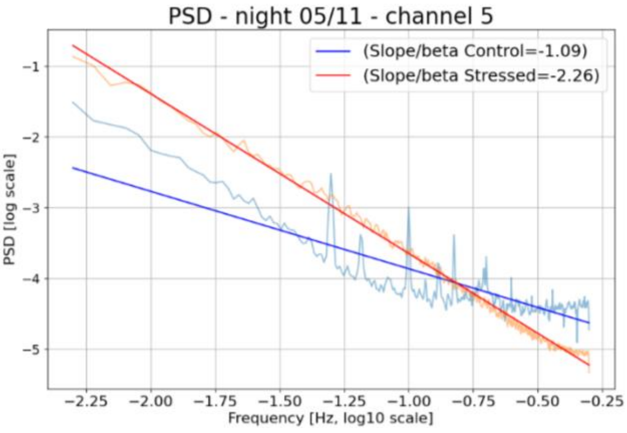
PSD - night 05/11 - channel 5
(Slope/beta Control=-1.09)
(Slope/beta Stressed=-2.26)
PSD [log scale]
Frequency [Hz, log10 scale]

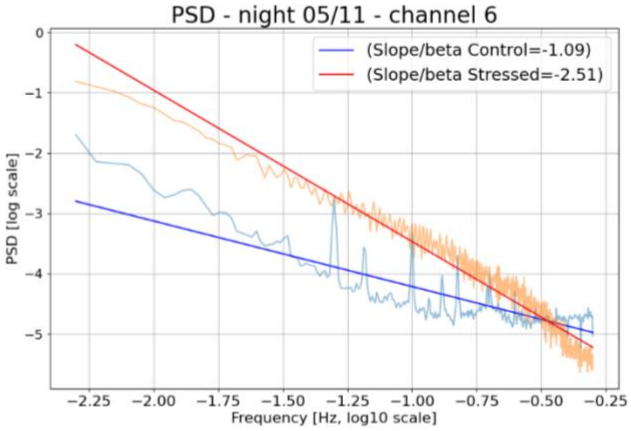
PSD - night 05/11 - channel 6
(Slope/beta Control=-1.09)
(Slope/beta Stressed=-2.51)
PSD [log scale]
Frequency [Hz, log10 scale]

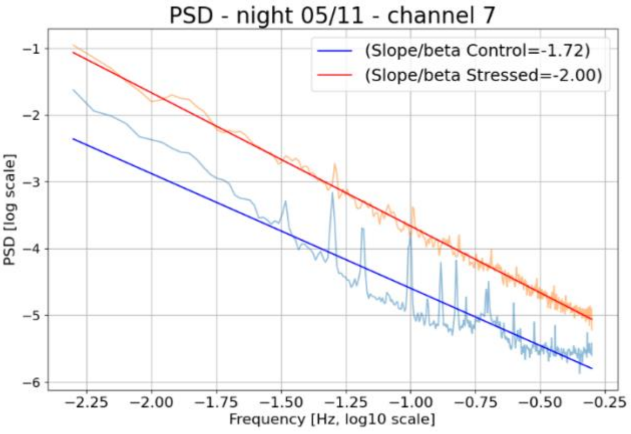
PSD - night 05/11 - channel 7
(Slope/beta Control=-1.72)
(Slope/beta Stressed=-2.00)
PSD [log scale]
Frequency [Hz, log10 scale]

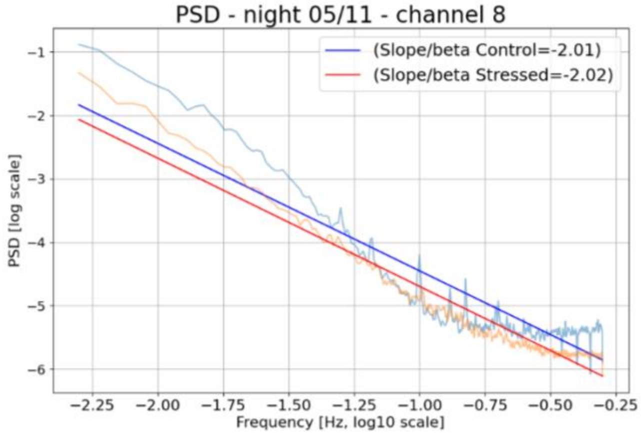
PSD - night 05/11 - channel 8
(Slope/beta Control=-2.01)
(Slope/beta Stressed=-2.02)
PSD [log scale]
Frequency [Hz, log10 scale]

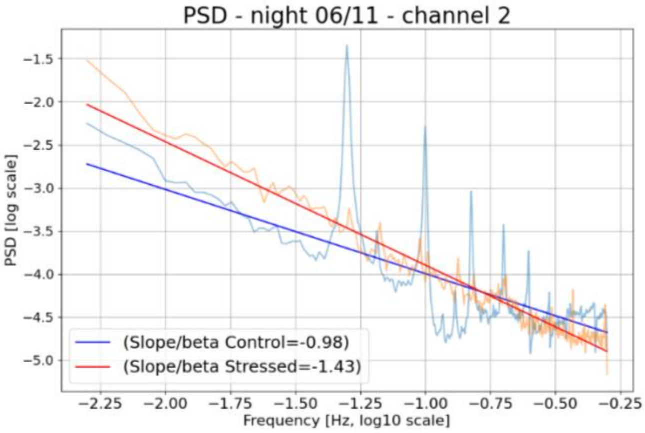
PSD - night 06/11 - channel 2
(Slope/beta Control=-0.98)
(Slope/beta Stressed=-1.43)
PSD [log scale]
Frequency [Hz, log10 scale]

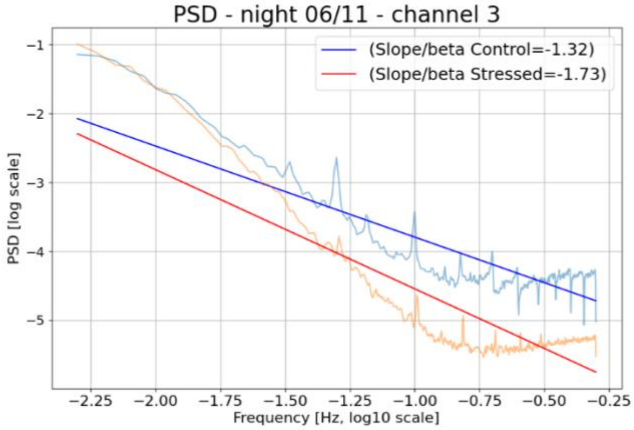
PSD - night 06/11 - channel 3
(Slope/beta Control=-1.32)
(Slope/beta Stressed=-1.73)
PSD [log scale]
Frequency [Hz, log10 scale]

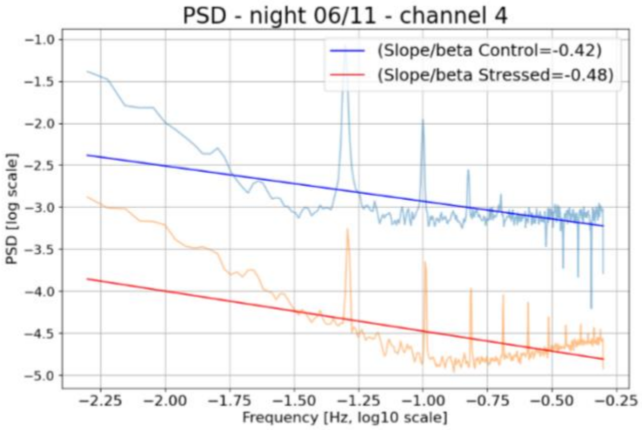
PSD - night 06/11 - channel 4
(Slope/beta Control=-0.42)
(Slope/beta Stressed=-0.48)
PSD [log scale]
Frequency [Hz, log10 scale]

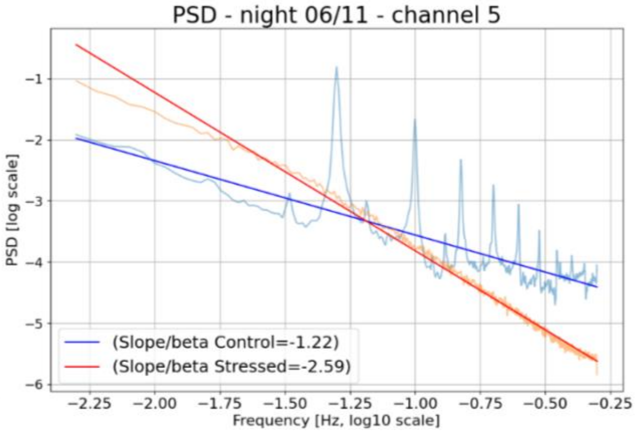
PSD - night 06/11 - channel 5
(Slope/beta Control=-1.22)
(Slope/beta Stressed=-2.59)
PSD [log scale]
Frequency [Hz, log10 scale]

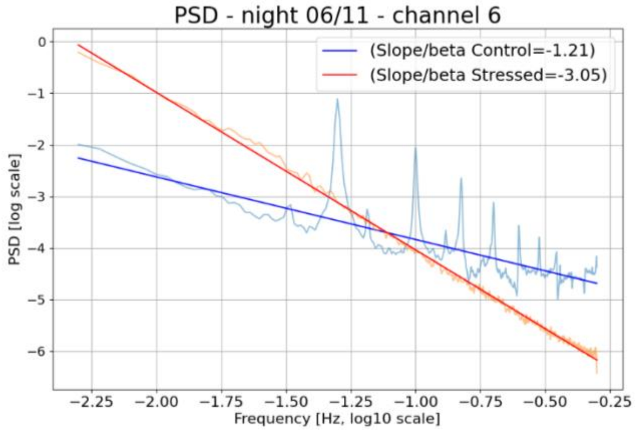
PSD - night 06/11 - channel 6
(Slope/beta Control=-1.21)
(Slope/beta Stressed=-3.05)
PSD [log scale]
Frequency [Hz, log10 scale]

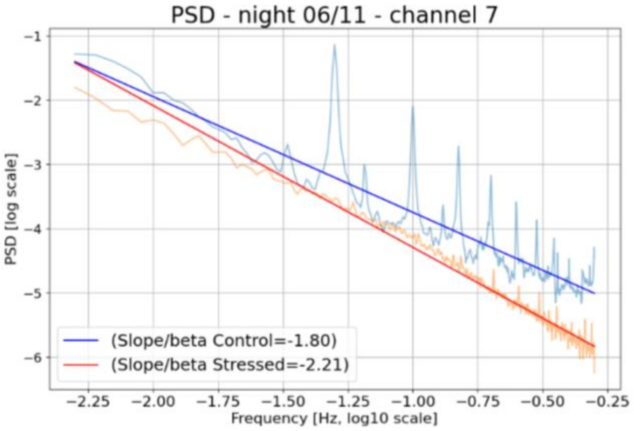
PSD - night 06/11 - channel 7
(Slope/beta Control=-1.80)
(Slope/beta Stressed=-2.21)
PSD [log scale]
Frequency [Hz, log10 scale]

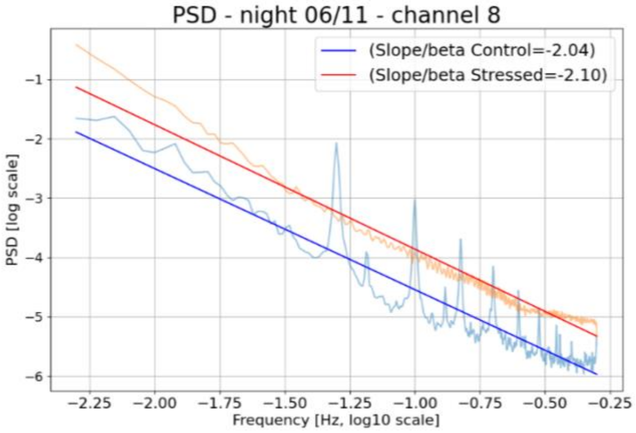
PSD - night 06/11 - channel 8
(Slope/beta Control=-2.04)
(Slope/beta Stressed=-2.10)
PSD [log scale]
Frequency [Hz, log10 scale]

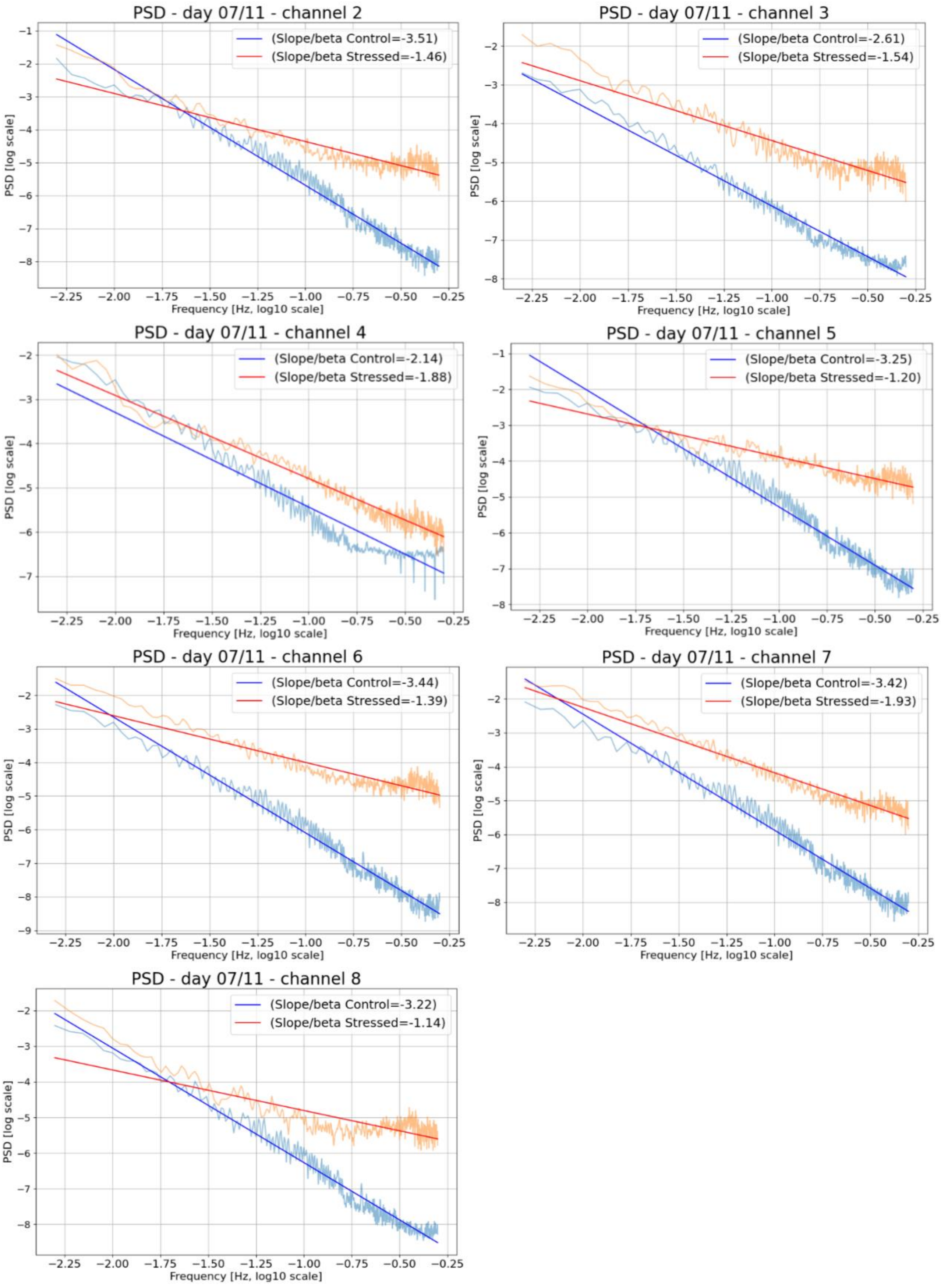


*Figure SM 3.2.3: Densities (PSDs) of the tomato plants, for each daylight and nighttime period. The daylight periods of 02/11, 04/11 and 06/11 could not be computed because of gaps in the data. The PSDs are plotted on a bilogarithmic plot and interpolated through a linear fitting line. A straight line, in the bilogarithmic plot, represents a power-law.*
