## Supplementary Material 4 for "Electrophysiological monitoring of plants: an exploratory study on drought stress"

**Supplementary Material 4 – Additional results for the XGB model**

Table SM4.1 reports the average accuracy values of the XGB model for each N-l combination and for both tomato plants (on the left) and apricot plants (on the right).

*Table SM4.1: accuracy values averaged over leave-one-out cross-validation results for the tomato plants (on the left) and the apricot plants (on the right).*

| N\l | 120 | 180 | 240 | 300 | 660 | 780 | 900 |
|---|---|---|---|---|---|---|---|
| 5 | 60.9% | 63.0% | 62.1% | 60.2% | 59.1% | 58.1% | 59.9% |
| 10 | 61.9% | 62.6% | 58.4% | 58.8% | 60.5% | 57.1% | 60.9% |
| 15 | 66.6% | 61.8% | 60.3% | 58.6% | 63.2% | 60.5% | 60.0% |
| 20 | 64.6% | 62.2% | 60.9% | 60.8% | 59.2% | 60.6% | 61.0% |
| 25 | 65.6% | 61.4% | 61.7% | 58.5% | 62.5% | 61.0% | 60.1% |
| 30 | 64.7% | 63.2% | 61.4% | 60.4% | 62.1% | 61.8% | 60.2% |
| 35 | 64.9% | 61.8% | 62.6% | 65.9% | 61.4% | 61.3% | 60.7% |

| N\l | 120 | 180 | 240 | 300 | 660 | 780 | 900 |
|---|---|---|---|---|---|---|---|
| 5 | 43.4% | 44.4% | 47.0% | 42.6% | 45.1% | 44.5% | 47.4% |
| 10 | 45.9% | 47.2% | 47.1% | 48.2% | 45.1% | 43.8% | 46.9% |
| 15 | 50.0% | 43.8% | 45.5% | 47.2% | 45.2% | 45.2% | 46.3% |
| 20 | 44.5% | 49.5% | 48.5% | 47.4% | 47.3% | 43.6% | 48.0% |
| 25 | 48.7% | 48.7% | 48.2% | 45.4% | 47.7% | 46.0% | 46.8% |
| 30 | 49.5% | 48.5% | 49.9% | 45.6% | 51.0% | 47.3% | 50.2% |
| 35 | 48.1% | 49.7% | 47.0% | 46.8% | 49.0% | 50.5% | 47.9% |

It is important to note that the values in Table SM4.1 result from an average of a number of accuracy matrices derived from the cross-validation procedure (seven for the tomato and six for the apricot). Therefore, it is crucial to consider the variability of each value, in terms of standard deviation, as presented in SM4.2.

*Table SM4.2: Standard deviation values of the accuracies among all the cross-validation trials for the tomato plants (on the left) and the apricot plants (on the right).*

| N\l | 120 | 180 | 240 | 300 | 660 | 780 | 900 |
|---|---|---|---|---|---|---|---|
| 5 | 28.5% | 27.5% | 28.9% | 28.9% | 28.1% | 27.4% | 27.4% |
| 10 | 28.6% | 28.9% | 27.2% | 26.8% | 29.3% | 26.6% | 29.2% |
| 15 | 30.1% | 28.5% | 28.2% | 26.5% | 29.9% | 28.3% | 28.9% |
| 20 | 29.0% | 29.0% | 28.3% | 29.3% | 28.2% | 29.6% | 28.7% |
| 25 | 29.8% | 29.0% | 28.8% | 25.7% | 30.4% | 30.9% | 29.4% |
| 30 | 30.1% | 28.3% | 29.0% | 27.1% | 30.1% | 29.4% | 30.2% |
| 35 | 30.1% | 28.4% | 29.0% | 30.0% | 28.7% | 29.5% | 30.2% |

| N\l | 120 | 180 | 240 | 300 | 660 | 780 | 900 |
|---|---|---|---|---|---|---|---|
| 5 | 8.40% | 8.40% | 6.10% | 7.40% | 5.30% | 5.80% | 6.80% |
| 10 | 7.80% | 5.00% | 5.60% | 5.20% | 7.10% | 7.80% | 7.90% |
| 15 | 5.90% | 8.10% | 8.70% | 7.70% | 9.20% | 6.00% | 7.50% |
| 20 | 9.40% | 5.60% | 3.90% | 7.40% | 9.30% | 7.40% | 10.00% |
| 25 | 8.50% | 5.00% | 7.70% | 5.80% | 8.00% | 8.80% | 8.10% |
| 30 | 9.40% | 6.50% | 5.70% | 8.50% | 10.10% | 9.50% | 10.10% |
| 35 | 8.80% | 4.60% | 7.50% | 6.20% | 9.30% | 7.00% | 9.10% |
